# Assessing chemical toxicity across Eukaryota using multimodal transformers

**DOI:** 10.64898/2026.09.23.753795

**Authors:** Styrbjörn Käll, Mikael Gustavsson, Patrik Svedberg, Mercedes Dalman, Jens Henriksson, Thomas Backhaus, Erik Kristiansson

## Abstract

Biodiversity is globally threatened by chemical pollution, yet toxicity data remain unavailable for millions of species and hundreds of thousands of chemicals, severely limiting our ability to assess ecological impacts. To meet this challenge, we developed TRIDENT-2, a multimodal artificial intelligence model for predicting chemical toxicity across evolutionarily diverse eukaryotic species. Trained on 560,780 toxicity assays spanning 82,775 chemicals, 6,793 species, and multiple exposure scenarios, TRIDENT-2 accurately predicts toxicity across Eukaryota with an average median absolute error ranging from 1.76 to 3.80 (fold-change). By jointly learning from chemical, biological, and experimental information, TRIDENT-2 remains accurate across broad chemical and taxonomic distances, allowing for toxicity assessment for species and chemicals beyond the current experimental evidence. Our findings demonstrate that artificial intelligence can help overcome longstanding data limitations in ecotoxicology, paving the way for improved decision-making and reducing chemical impacts on biodiversity and ecosystems.

## Main

The United Nations has identified climate change, chemical pollution and biodiversity loss as interlinked challenges in the triple planetary crisis (*Global Resources Outlook 2024*, 2024). Pollution is also independently identified as one of the five major drivers of biodiversity loss and one of the nine planetary boundaries (IPBES, 2019; Persson et al., 2022; Rockström et al., 2009; Sánchez-Bayo & Wyckhuys, 2019). Indeed, evidence of adverse effects from chemicals spans the eukaryotic tree of life and has e.g. been linked to the widespread decline of insects and pollinators (Liess et al., 2021; Nicholson et al., 2024; Sánchez-Bayo & Wyckhuys, 2019; Sgolastra et al., 2020), collapsing bird populations (Hallmann et al., 2014; Oaks et al., 2004; Swan et al., 2006), and altered species compositions in both aquatic and terrestrial biomes (Keck et al., 2025; Malaj et al., 2014; Posthuma et al., 2020; Schäfer et al., 2026). In Europe, chemical pollution is associated with the loss of approximately 20 % of aquatic species (*The European Environment — State and Outlook 2020*, 2020) and significant economic burdens are linked to the degradation of ecosystem services and the cost for remediation (Chagnon et al., 2015; Directorate-General for Environment, European Commission et al., 2026). Consequently, to safeguard biodiversity, ecosystems, and their functions, it is essential to understand how chemical pollution adversely affects living organisms.

Although ecotoxicological studies have been conducted for decades, they are primarily limited to standardized tests, where a small group of selected algae, aquatic invertebrates, fish, plants, and rodents make up the vast majority of the reported data (*OECD Guidelines for the Testing of Chemicals, Section 2*, 2026; Olker et al., 2022). Strong biases also exist for chemical data, and less than one hundred chemicals make up more than half of the ecotoxicological scientific literature (Kristiansson et al., 2021). In contrast, estimates put the number of eukaryotic species at 8.7 million (Mora et al., 2011) and the European Environmental Agency has estimated that more than 100,000 chemicals are present on the European market (*The Unknown Territory of Chemical Risks*, 2019) while more than 500,000 new chemicals are discovered annually (Llanos et al., 2019). Most chemicals and eukaryotic species are, therefore, untested and potential effects are unknown. Meanwhile, species sensitivity can vary by several orders of magnitude, even when the species are closely related, making assessment of chemical impact on ecosystems exceedingly difficult (Blanck et al., 1984; González-Vázquez et al., 2025).

Recently, toxicity predictions using artificial intelligence (AI)-based methods have shown promise as alternatives to empirical testing (Anand et al., 2024; Fan et al., 2021; Forastiere et al., 2026; Mayr et al., 2016; Posthuma et al., 2025; Raimondo & Barron, 2020; Togo et al., 2023; Yuan et al., 2026; Zubrod et al., 2024). In particular, state-of-the-art deep learning (DL) approaches significantly improve performance compared to conventional quantitative structure-activity relationship (QSAR) methods (Gustavsson et al., 2024). However, existing methods remain limited to a narrow number of species or are typically restricted to species groups used in standard environmental risk assessment (such as algae, crustaceans, fish, and rodents) . Existing computational methods for toxicity prediction, therefore, neglect major parts of the eukaryotic tree of life, making them unsuitable for environmental risk assessment at the ecosystem scale. Meanwhile, recent advances in multimodal AI enable integration of diverse sources of information into a single model, improving performance beyond what is possible with individual data types (Abramson et al., 2024; Brixi et al., 2026). As chemical toxicity emerges from complex biochemical interactions, multi-scale species biology, and exposure conditions, multimodal AI provides a compelling framework for advancing chemical toxicity assessment across evolutionarily diverse organisms.

Therefore, we developed TRIDENT-2, a multimodal transformer-based AI model to predict chemical toxicity towards aquatic and terrestrial eukaryotic species. The model was trained on 560,780 exposure assay resultsspanning 6,793 species and 82,775 chemicals. The training data also included a diverse set of toxicological effects, endpoints, organism life stages, and routes of administration. We show that the accuracy of toxicity predictions benefits greatly from joint learning from diverse data, enabling accurate extrapolation across Eukaryota. TRIDENT-2 markedly improves accuracy compared to the current state of the art, enabling estimates of toxicity beyond current experimental data for many previously untested chemicals and taxa. We conclude that multimodal AI can address fundamental knowledge gaps in toxicology, considerably improving our ability to assess chemical impacts on biodiversity and ecosystems. TRIDENT-2 is available as a free-to-use service through https://trident.serve.scilifelab.se.

## Results

### Toxicity data is diverse but sparse

We collected and harmonized toxicological data from four major data sources (ECOTOXicology Knowledgebase, OpendFoodTox, REACH dossiers and Registry of Toxic Effects of Chemical Substances (RTECS))/(US EPA, EFSA, ECHA, and CCOHS), together comprising 560,780 exposure test results, spanning four toxicological endpoints, 6,793 species, 82,775 chemicals, 10 toxicological effects, 128 organism life stages, 304 routes of administration, and 14 concentration units. Toxicity levels ranged over 10 orders of magnitude (**Fig. 1a-f, Supplementary Table S1 and Fig. S1-S2)**, and exposure durations ranged from hours to months. Species taxonomy was diverse, including 4 kingdoms, 44 phyla, 139 classes, 450 orders, 1,674 families, and 4,549 genera, in total covering 34.5 % of Eukaryota (**Fig. 1c**). The dataset exhibited skewness, with, e.g., rodent exposure assays making up 26.7 % of the total observations, being the only tested species for 84.4 % of the chemicals. Likewise, insects made up 24.5 % of all species, and mortality made up 50.2 % of the reported toxicological effects.

**Figure 1.**
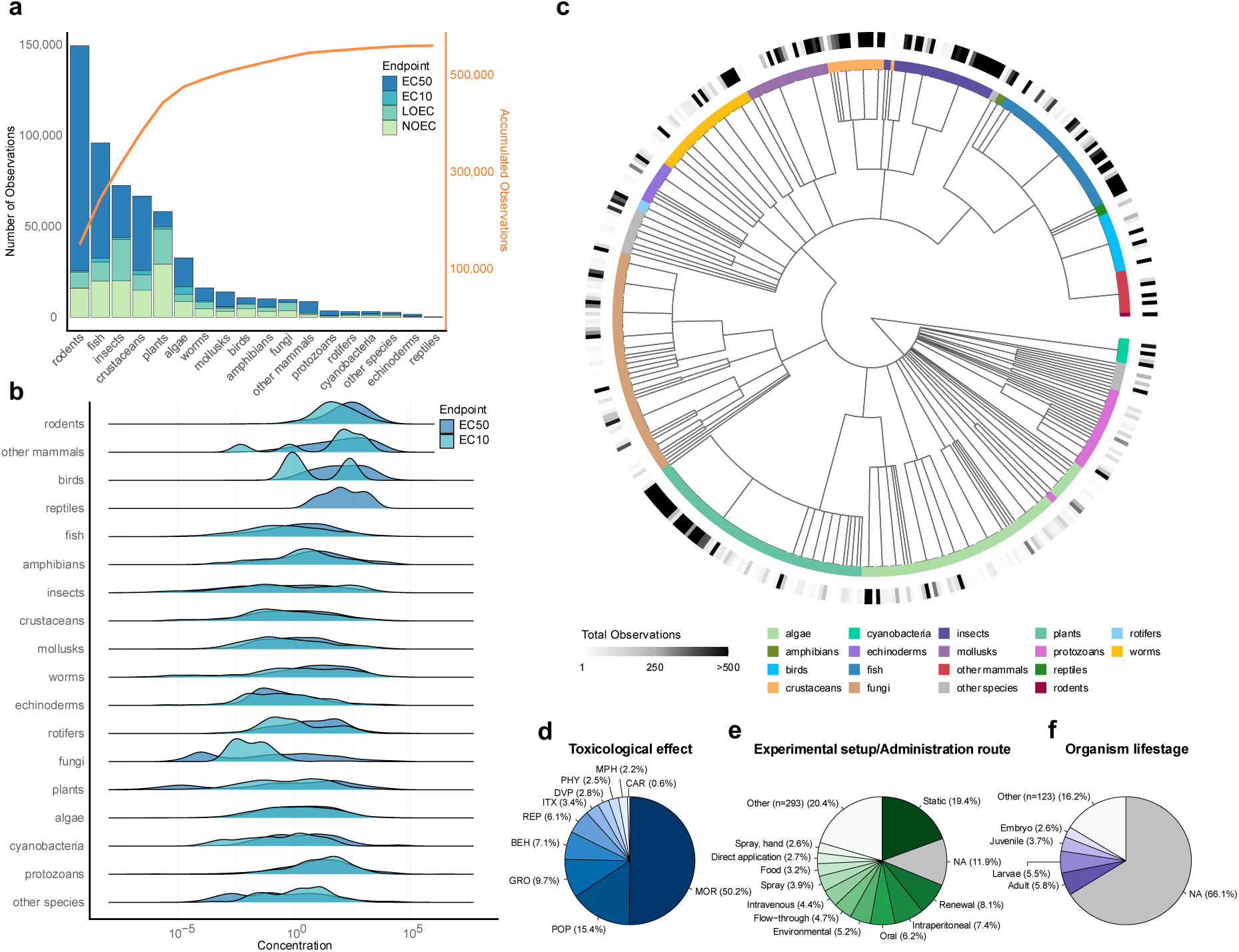
Dataset description. **(a)** Number of observations per taxon and toxicity endpoint, with associated cumulative count (right axis). Although not members of Eukaryota, cyanobacteria were kept as they are a common model organism. **(b)** EC50 (n=319,528) and EC10 (n=15,158) concentrations with 50 % and 10 % effect response distributions per taxon, see Supplementary Fig. S1 for No Observable Effect Concentration (NOEC) and Lowest Observable Effect Concentration (LOEC). **(c)** Taxonomic tree showing the taxonomic orders covered (n=450) by the 6,793 species in the dataset. Colored tiles indicate eukaryotic groups, and the intensity indicates the number of observations per order. **(d-f)** Distribution of toxicological effects, experimental setups/routes of administration, and organism life stages. Categories making up less than 5 % of the total observations were grouped into “Other”. “NA” indicates missing information. See Supplementary Fig. S2 for harmonized ratios. (BEH = Behavior. CAR = Carcinogenicity. DVP = Development. GRO = Growth. ITX = Intoxication. MOR = Mortality. MPH = Morphology. PHY = Physiology. POP = Population. REP = Reproduction.)

### Multimodal modelling of chemical toxicity

We developed a multimodal transformer model (TRIDENT-2) to predict chemical toxicity towards eukaryotic species (**Fig. 2**). The model is designed to simultaneously output four toxicity endpoints (EC_50_, EC_10_, LOEC, and NOEC) at the species level, based on chemical structure, toxicological effect (mortality, population, growth, behaviour, reproduction, intoxication, development, physiology, morphology, and carcinogenicity), and exposure scenario, including route of administration, organism life stage, exposure duration, and concentration unit.

**Figure 2.**
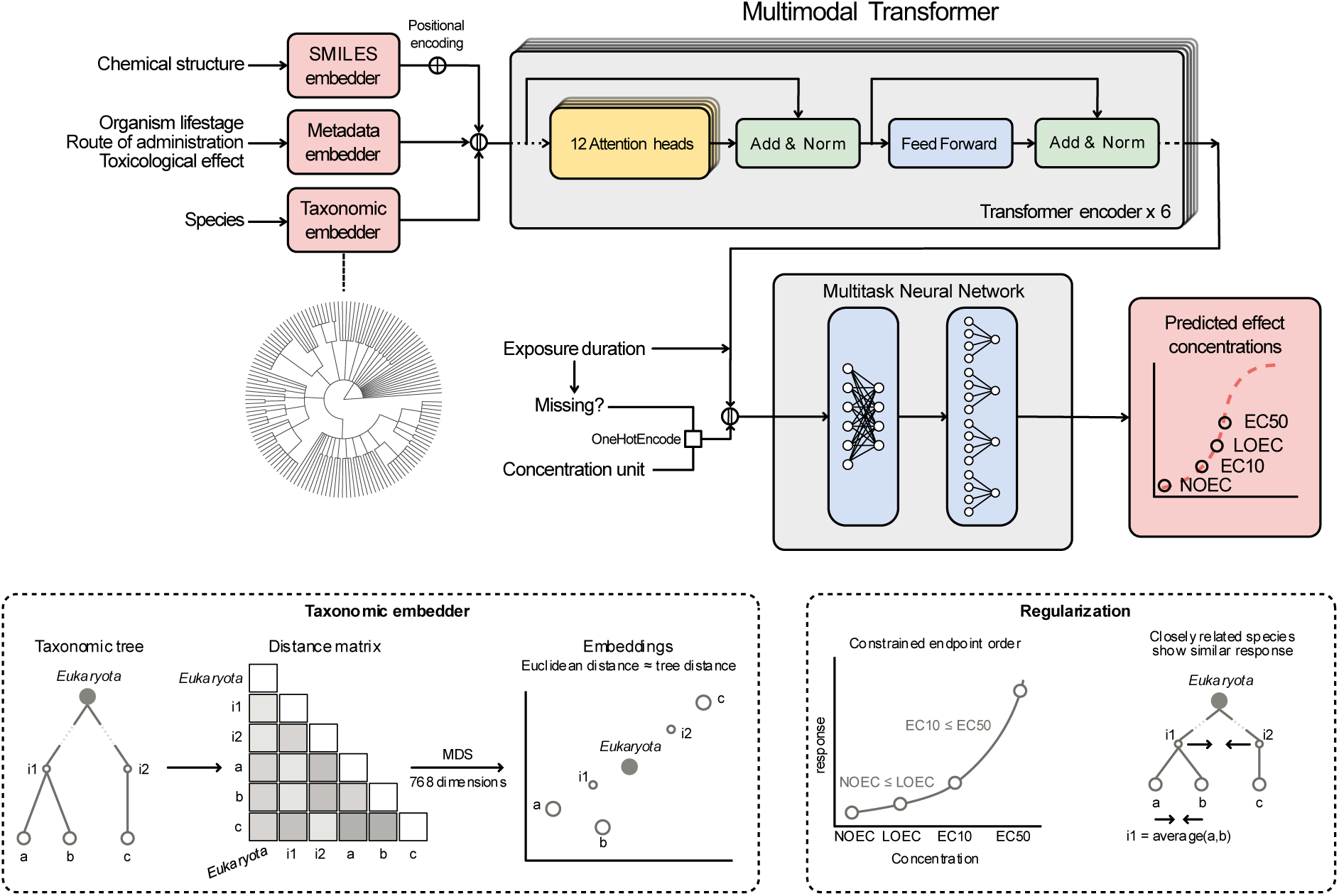
Model description. Chemical structures, toxicological effects, routes of administration, organism life stages, and species are embedded as 768-dimensional embedding vectors and fed to the multimodal transformer encoder (50,475,892 parameters). Chemicals, represented as SMILES strings, effects, route of administration, and organism life stages are embedded with learnable token embeddings. To respect taxonomic distances, species taxonomy is embedded with non-learnable embeddings from a metric MDS of the taxonomic distance matrix. A prepended CLS-embedding is extracted from the transformer after sequential updates through self-attention. The embedding is concatenated with the exposure duration (log10 hours), a one-hot encoding of the concentration unit, and a flag when the duration is missing. The multitask neural network (236,404 parameters) uses the concatenated representation to predict four toxicological endpoints: EC50, EC10, LOEC, and NOEC, using a shared hidden layer and four task-specific layers. During training, two regularization terms are added where endpoint predictions are penalized if ordered incorrectly, and predictions for evolutionarily similar species are pulled together through averages, propagating down the taxonomic tree to mitigate data sparsity and cause species with little data to learn from related species.

The model uses vector embeddings to capture relationships among data modalities (**Fig. 2**). Chemicals are encoded as SMILES strings (Simplified Molecular Input Line Entry System), which, together with toxicological effects, routes of administration, and organism life stages, are represented by embeddings inferred from the data. A fused species-level toxicity representation is generated by integrating information from the taxonomic tree, reflecting evolutionary relationships between taxa. Four neural networks with a shared hidden layer predict the toxicity for the four included endpoints. Task-specific regularization inhibits endpoints from violating their natural ordering and encourages similar toxicity predictions for closely related species.

### Stable prediction errors and high extrapolation capacity

Model performance was measured using three different sets of ten-fold cross-validation. The validation sets were either based on unseen species and chemical combinations (**Fig. 3a, b**), unseen species (**Fig. 3c**), or unseen chemicals (**Fig. 3d**).

**Figure 3.**
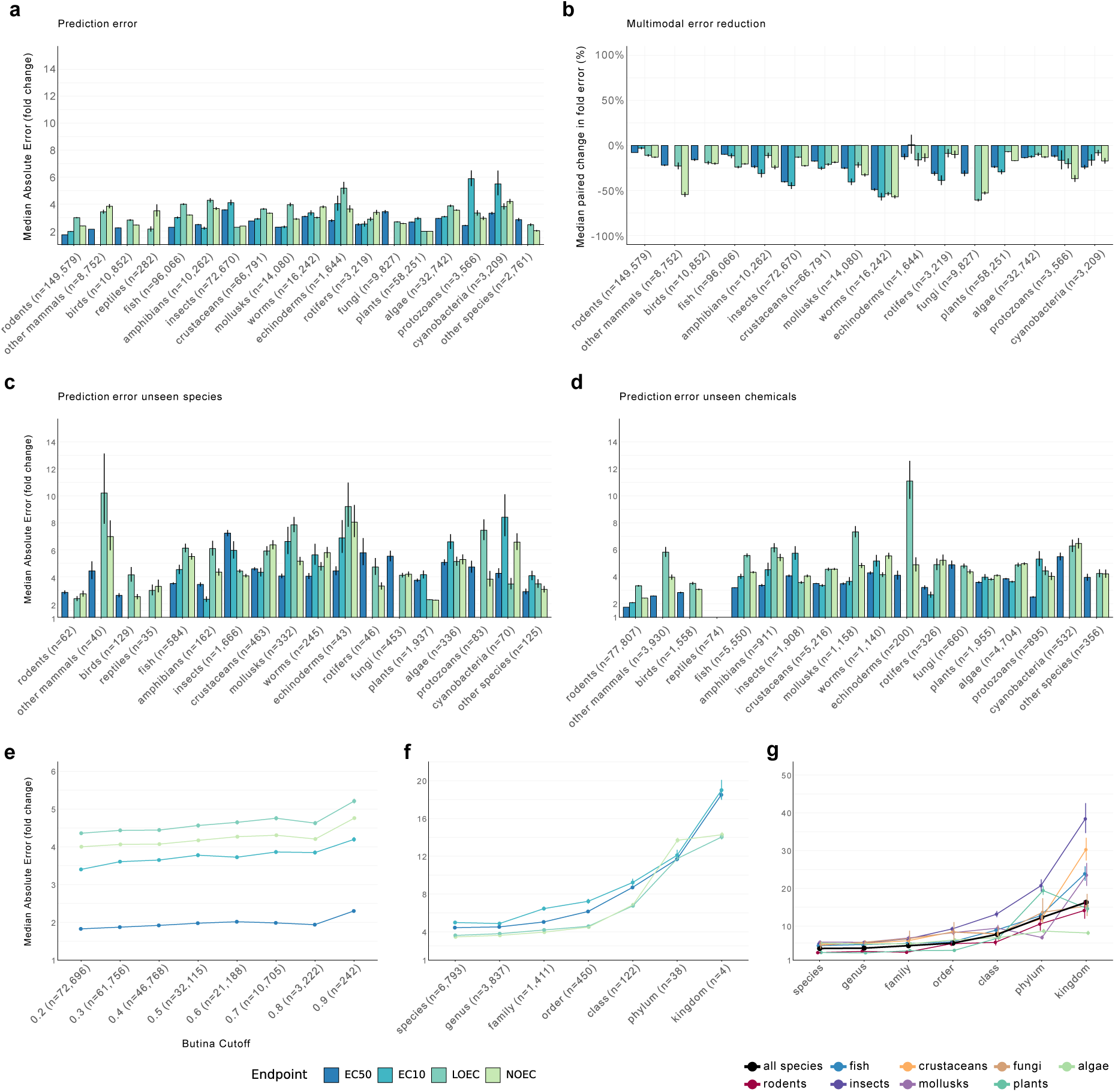
Model prediction error and extrapolation capability. Median Absolute Error (fold change) (MAE) from ten-fold cross-validation with error bars showing the Median Absolute Deviation (MAD) divided by √(n). **(a)** MAE when evaluating the model on unseen combinations of species and chemicals (for chemicals at Tanimoto similarity < 0.8, n is the number of observations in each species group, see also Supplementary Fig. S3). **(b)** Percentual change in MAE between the models trained on individual species groups and the multimodal model. Values were computed from paired differences in log10 absolute error and back-transformed to the fold-error scale. Negative values indicate lower prediction error for the multimodal model (n is the number of unique observations in each species group, see also Supplementary Fig. S4-S7). **(c)** MAE when evaluating the model on unseen species for each species group (n denotes the number of unique species in each group, see also Supplementary Fig. S8). **(d)** MAE when evaluating the model on unseen chemicals (Tanimoto similarity less than 0.8) for each species group (n denotes the number of unique chemicals in each group, see also Supplementary Fig. S9). **(e)** MAE when evaluating the model on unseen chemicals with decreasing Tanimoto similarity/Butina cutoff (n denotes the number of unique Butina clusters). The large difference in error between EC50 and other endpoints is partly due to the large amount of rodent data (see also Supplementary Fig. S10)**. (f)** MAE when evaluating the model on species from unseen genera, families, orders, classes, phyla, and kingdoms, respectively (n denotes the number of taxa at each rank, see also Supplementary Fig. S11). **(g)** MAE when evaluating the model on species from unseen genera, families, orders, classes, phyla, and kingdoms, stratified by eukaryotic group, averaged across endpoints (see also Supplementary Fig. S12). The first scatter point (species) indicates MAE for a random subset of unseen species.

When evaluated on unseen species and chemical combinations, the model showed robust performance for all evaluated eukaryotic groups, with average median absolute error fold changes (MAE) between 1.76 and 3.80 across the four endpoints (**Fig. 3a**). Prediction errors were, on average, lowest for species of rodents, plants, and other mammals (1.74-2.18), while the highest errors were found in echinoderms and cyanobacteria (3.31-3.64). Errors varied between endpoints, with slightly lower errors for EC_50_ and EC_10_ than for NOEC and LOEC, consistent with the variability observed in the data (**Supplementary Fig. S1**). Across toxicological effects, errors also remained stable, with only 20 out of 110 endpoint-effect combinations being above four MAE fold change and only three exceeding a value of six (**Supplementary Fig. S3**). In general, mortality (MOR), carcinogenicity (CAR), and population (POP) were associated with low errors, whereas physiology (PHY), morphology (MPH), and behavior (BEH) were associated with large errors. Some combinations of taxa and endpoints showed elevated prediction errors (e.g., EC_10_-fungi, EC_10_-protozoans, and EC_10_-cyanobacteria), likely due to low data availability. In contrast, rodents, represented by comprehensive data largely generated by highly standardized tests, showed low prediction errors overall.

On average, the multimodal model showed a 25.2 % reduction in prediction error compared to models trained individually on each eukaryotic group, thus demonstrating efficient extrapolation across the tree of life (**Fig. 3b, Supplementary Fig. S4-S7**). Performance gains were observed for all groups, even for rodents, fish, and plants, where data were most abundant. The largest improvements, however, were seen in less data-abundant groups, especially fungi, worms, and insects, where errors were reduced by an average of 53.9 %, 53.1 %, and 32.1 %, respectively. Similarly, although less prominent, errors decreased when including information on route of administration and organism life stage, indicating that they capture complementary information relevant to assessing toxicity (**Supplementary Fig. S13**).

To evaluate the capability to support ecosystem impact assessment beyond existing exposure data, we assessed errors for unseen species and chemical structures independently (**Fig. 3c-d, Supplementary Fig. S8-S9**). For unseen species, MAE ranged from 2.27 to 10.21 (**Fig. 3c**), with plants, birds, and rodents showing the lowest errors. For unseen chemicals, MAE was generally lower, ranging from 1.75 to 11.10 (**Fig. 3d**). This indicates that it is more difficult to estimate the toxicity of a previously untested species than an untested chemical in a well-studied species.

To further explore the model’s ability and limitations in extrapolating to untested chemicals and species, we evaluated errors from two independently trained sets of models with progressively stricter restrictions on similarity between species and chemicals in the training and validation sets. As chemical dissimilarity increased, errors rose across all endpoints (**Fig. 3e, Supplementary Fig. S10)**. However, the increase in errors was modest, even for largely dissimilar chemicals (1.82 to 2.30 for EC_50_ and 3.40 to 4.20 for EC_10_ at Butina cutoff 0.2 and 0.9, respectively). This indicates strong potential for assessing toxicity in untested chemicals, also from new structural classes. With increasing evolutionary distance, the average MAE increased monotonically, from 4.08 for species in untested genera to 16.30 in untested kingdoms (**Fig. 3f and Supplementary Fig. S11**). Errors were moderate up to taxonomic family and order levels (4.50-7.23 MAE), suggesting that extrapolations to previously untested species should not exceed too large evolutionary distances. Insects deviated earlier than other groups, suggesting that extrapolation in this part of the tree of life is especially difficult (MAE = 9.25 compared to, e.g. 5.81 for fish at the order level) (**Fig. 3g, Supplementary Fig. S12**). The correlation between error and distance, measured by the cosine similarity in the model’s embedding space, was also observed between predictions and the respective nearest neighbors in the training set. This distance can thus be used to obtain estimates of prediction reliability (**Supplementary Fig. S14**).

### Benchmarking shows improvements over the state-of-the-art

We benchmarked TRIDENT-2 against five other methods: three established toxicity prediction tools—ECOSAR (Wright et al., 2022), VEGA (Roncaglioni et al., 2022), and T.E.S.T. (Martin, 2020)—as well as TRIDENT-1 (Gustavsson et al., 2024), a transformer-based model trained for aquatic species groups, and a pairwise learning model (Posthuma et al., 2025).

We first compared the applicability of the methods, measured by the number of supported orders in Eukaryota, unique species, chemical structures, endpoints, and effects (**Fig. 4a-b**). TRIDENT-2 surpassed all competitors and encompasses 34.5 % of all existing eukaryotic orders, 209.3 % more orders than ECOSAR, 6,793 species, 5,526 more than the pairwise learning model, and 100 % of the chemicals, 57.5 % more than ECOSAR. For the other methods, predictions were limited to fish, worms, algae, invertebrates, and rats, and to a smaller selection of endpoints, effects, and chemicals, showing that TRIDENT-2 represents a major advance in assessing chemical toxicity.

**Figure 4.**
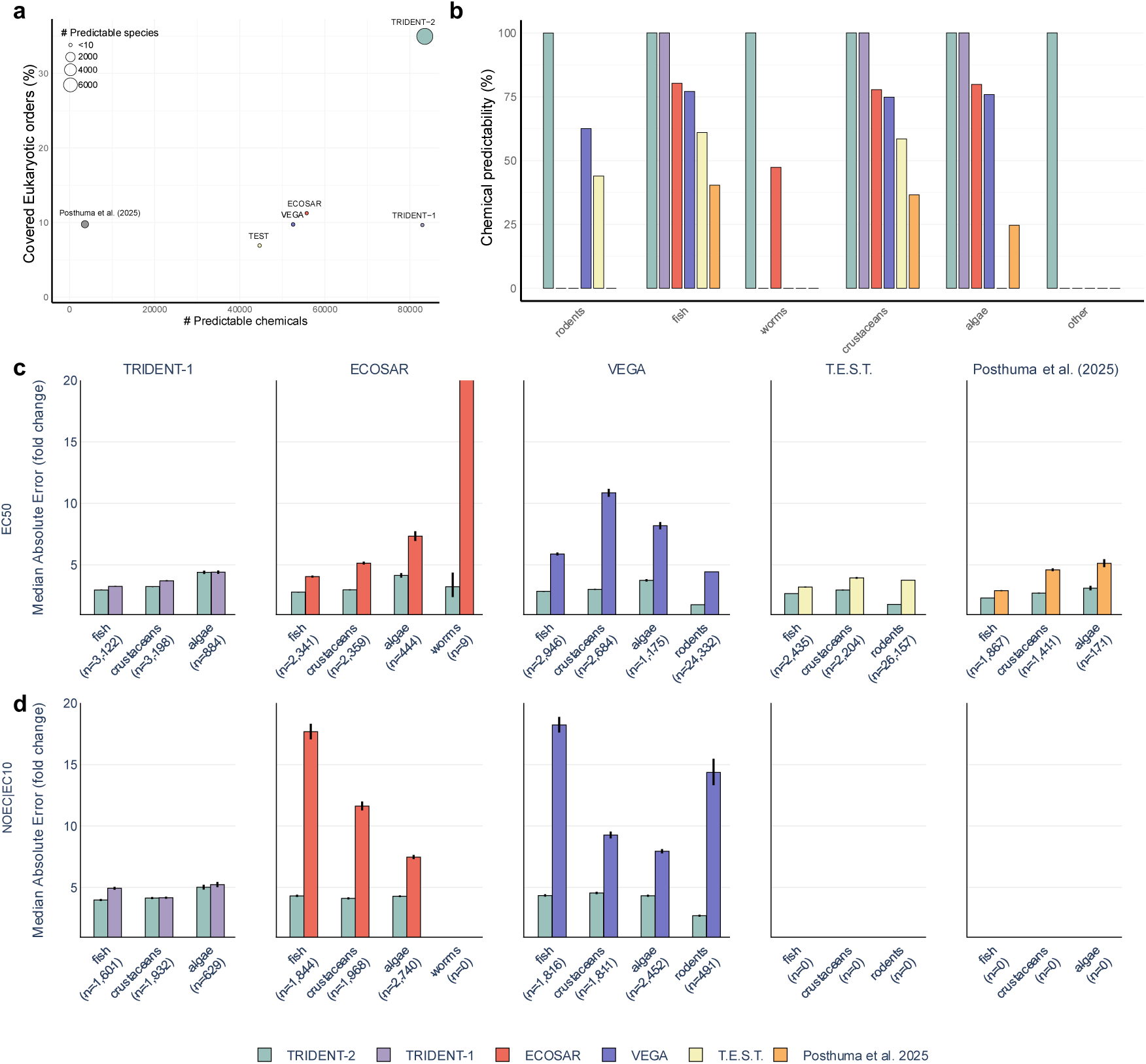
Model benchmarking. **(a)** The number of predictable chemicals from the full training set and the covered taxonomic orders in Eukaryota (%). Circle size reflects the number of predictable Eukaryotic species. **(b)** Chemical predictability (%) for the chemicals in the full training set. **(c)** Pairwise comparison of EC50 Median Absolute Error, MAE (fold change). Prediction error for ECOSAR-worms was 81.99 (MAD=+40.42/-27.07). **(d)** Pairwise comparison of combined EC10 and NOEC MAE (fold change). Performance improved for ECOSAR and VEGA when comparing only chemicals within their respective applicability domains (Supplementary Fig. S15 and Table S2-S3) and improved for all methods when considering only chemicals within their joint applicability domain (Supplementary Fig. S16 and Table S4-S5). Error bars show the Median Absolute Deviation (MAD) divided by √(n), where n is the number of unique chemicals in each comparison. Absent bars indicate the compared model cannot produce predictions for that combination of species group and endpoint. As only TRIDENT-2 could produce LOEC predictions, we only compared EC50 and pooled EC10 and NOEC predictions to retain sufficient data for comparison (n denotes the number of unique chemicals in each comparison).

Next, we conducted pairwise comparisons of prediction errors between the models (**Fig. 4c-d, Supplementary Table S2-S3**). TRIDENT-2 consistently outperformed all other models for both EC_50_ and EC_10_/NOEC predictions for all eukaryotic groups. Error differences were most prominent for EC_10_/NOEC predictions from ECOSAR and VEGA, with errors being 74-431 % larger than for TRIDENT-2. Meanwhile, T.E.S.T., TRIDENT-1, and Posthuma et al. 2025 performed roughly on par, with errors between 20-26 % higher for the best-performing group (EC_50_-fish) and 33-70 % higher for the worst-performing group (EC_50_-crustaceans). When comparing chemicals inside ECOSAR’s and VEGA’s respective strict applicability domains, predictability decreased while prediction errors improved, yet TRIDENT-2 remained superior (**Supplementary Fig. S15**, **Supplementary Table S2-S3**). Finally, we compared errors for the chemicals falling within the joint applicability domain of ECOSAR and VEGA, greatly reducing the number of chemicals in the comparison. Compared to the second-best model, TRIDENT-2 achieved 22 % and 38 % lower errors for EC_50_ and EC_10_/NOEC, respectively (**Supplementary Fig. S16, Supplementary Table S4-5**).

### Pesticide toxicity varies at the species level

To assess the impact of chemicals on biodiversity it is vital to obtain comprehensive toxicity data at the species-level. We therefore explored whether TRIDENT-2 could recover taxonomic toxicity patterns while resolving species-specific sensitivity across the eukaryotic tree. We chose pesticides as a relevant test case since they cover a diverse set of chemicals posing widespread threats to ecosystems and biodiversity through their toxicity, global use, and direct release into the environment (Brühl & Zaller, 2019; Schäfer et al., 2026). TRIDENT-2 was applied to 2,146 pesticides collected from the Pesticide Properties Database (PPDB) and the Biopesticide Database (BPDB) (Tzilivakis et al., 2026) to assess toxicity for 6,793 eukaryotic species, in total encompassing 14,577,778 species-chemical combinations. Predicted EC_50_ values for mortality, population, and growth effects were normalized across species and chemicals to improve comparability (see Methods and **Supplementary Fig. S17** for details).

Analysis of the toxicity at the taxonomic order level revealed distinct patterns associated with pesticide classes (**Fig. 5a**). For the six most common pesticide types, TRIDENT-2 identified homogeneous clusters of sensitive orders, consistent with their expected biological targets. Particularly, most herbicides and insecticides were predicted to be highly toxic across plant and insect orders, respectively, whereas specific insecticides showed elevated predicted toxicity across a broader range of orders. Several pesticide types produced heterogeneous clusters spanning multiple eukaryotic orders, indicating that toxicity was not restricted to their primary targets.

**Figure 5.**
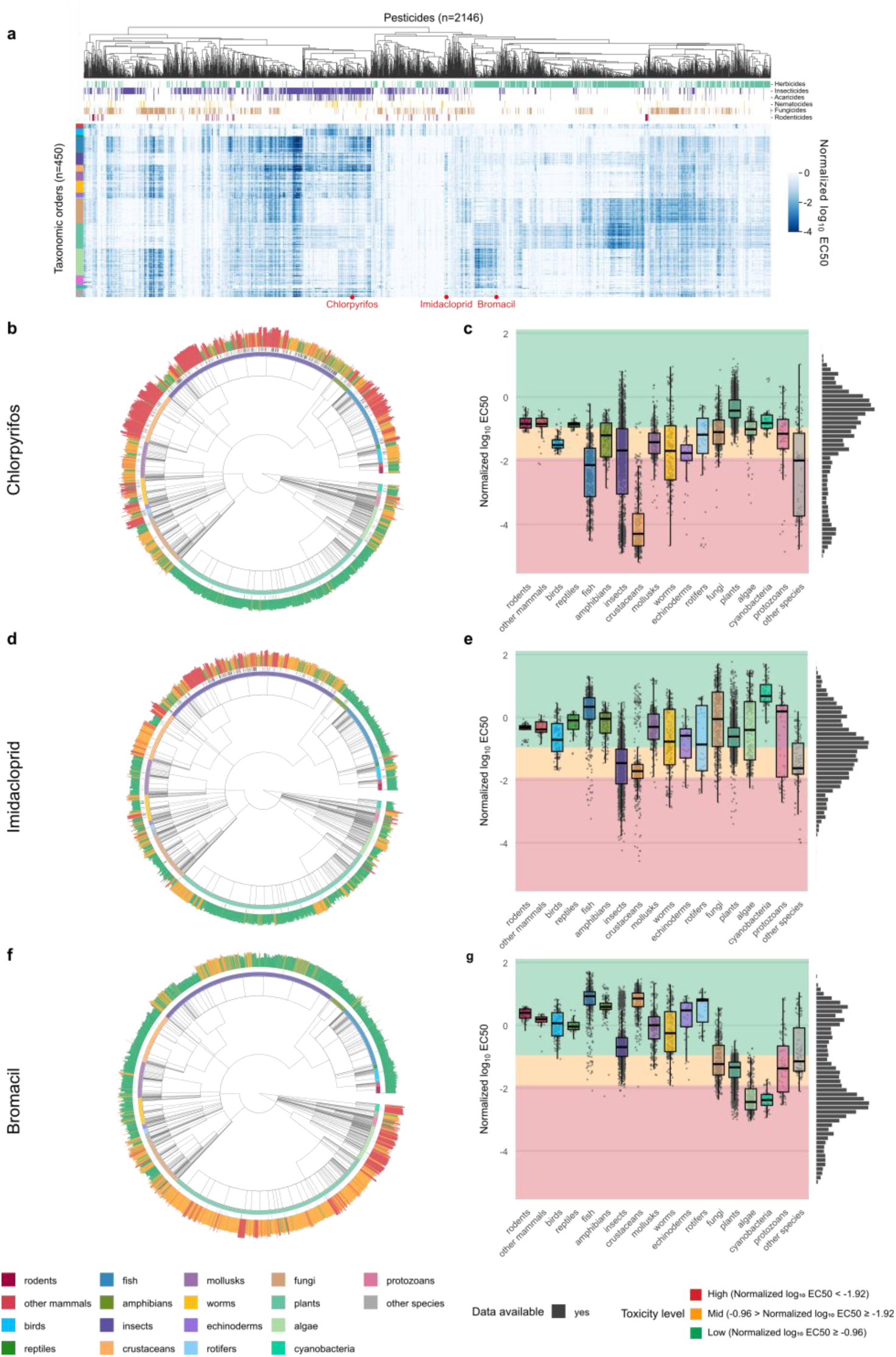
Analysis of model predictions across 6,793 eukaryotic species for 2,146 pesticides. **(a)** Model EC50 predictions across the 6,793 eukaryotic species in the dataset aggregated to the taxonomic order level (n=450) for 2,146 pesticides collected from the Pesticide Properties Database (PPDB) and the Biopesticides Database (BPDB). To increase comparability, the predicted EC50 was normalized for each species by the median prediction across 12,830 chemicals in the dataset (n=92,316,870 predictions, chemicals where only rodent data was available were excluded). Normalized predictions were averaged at the taxonomic order level. Chemicals were then grouped using hierarchical clustering based on Pearson correlation and average linkage. Column color bands indicate pesticide types from PPDB and BPDB and row color bands indicate eukaryotic groups. **(b, d, f)** Taxonomic trees showing predicted normalized EC50 per species as bars. Bar height indicates predicted normalized EC50, scaled by a constant to avoid negative bars, and colored based on the number of standard deviations from the mean prediction. Black tiles indicate species where experimental data is available (chlorpyrifos = 764/6,735; imidacloprid = 430/4,576; bromacil = 49/193; “number of species with data”/”number of observations”). Tree branches are cropped to the taxonomic order level. **(c, e, g)** Predicted normalized EC50 per species for the three pesticides: chlorpyrifos, imidacloprid, and bromacil for each species group. Each point is an individual species. Background color patches indicate the number of standard deviations from the mean prediction. Histograms illustrate aggregated distributions across all species.

Next, we explored predictions at the species level for three pesticides with different target organisms: Chlorpyrifos, imidacloprid, and bromacil (**Fig. 5b-g**, see **Supplementary Fig. S18** for a comparison between experimental data and predictions). Chlorpyrifos, an organophosphate insecticide, was, as expected, predicted to be most toxic towards aquatic arthropods, with crustaceans and insects followed by fish (Giddings et al., 2014), whereas birds and mammals were predicted to be considerably less sensitive (Barron & Woodburn, 1995). Predicted sensitivity varied by orders of magnitude within groups, e.g., among crustaceans and insects, consistent with experimental observations (Rubach et al., 2011, 2012). This information was largely lost when considering aggregated EC_50_ predictions at the order level or higher taxonomy. The neonicotinoid insecticide imidacloprid, a neurotoxin that binds to nicotinic acetylcholine receptors, appeared most toxic to insects. However, predicted EC_50_ varied greatly between species of insects, with mayflies, caddisflies and dipterans among the most sensitive (**Fig. 5d-e**). Moreover, certain species of crustaceans and worms also showed elevated responses, while, e.g., *Daphnia* were predicted insensitive, consistent with reported non-target effects of Imidacloprid (Malhotra et al., 2021; Morrissey et al., 2015; Touzout et al., 2021; Wan et al., 2025; Wang et al., 2019). Finally, toxicity for bromacil, a relatively understudied herbicide that disrupts photosynthesis by blocking the electron transport chain, was predicted to be high towards species of plants, algae, and cyanobacteria (**Fig. 5f-g**). However, certain species of insects and certain species of fungi were also predicted to be sensitive to bromacil for which no supporting experimental data was available. Ultimately, these examples demonstrate that TRIDENT-2 can capture broad, biologically meaningful patterns while retaining species-level resolution, enabling the identification of toxicological responses among species for which there is no or little empirical data.

## Discussion

In this study, we show that TRIDENT-2, a deep learning multimodal transformer model, can accurately predict chemical toxicity across thousands of aquatic and terrestrial species, spanning evolutionarily diverse branches of the eukaryotic tree of life. TRIDENT-2 extends toxicity assessment to species and chemicals for which experimental data are scarce or completely missing, effectively bridging several of the fundamental knowledge gaps within chemical risk assessment and enabling identification of previously overlooked environmental threats. This provides decision-makers with new evidence on poorly represented species, supporting more informed prioritization of chemicals and recognition of especially vulnerable organisms, enabling improved strategies for reducing chemical impacts on biodiversity.

### Expanding the scope of ecotoxicology using AI

TRIDENT-2 was trained and evaluated on data from more than half a million chemical exposures. Our evaluations show that TRIDENT-2 achieves consistently high predictive performance across both less studied species groups—e.g., reptiles, amphibians, birds, echinoderms, tunicates, sponges, bryozoans, fungi, rotifers, and protozoans—and groups that contain more commonly tested groups, such as rodents, fish, crustaceans, and algae. The performance is maintained over large chemical and taxonomic distances, with low errors also for species in unseen families and orders, and, in some cases, even classes and phyla. This achievement can largely be attributed to the comprehensiveness and diversity of the dataset used to train the model, covering more than 80,000 chemicals and 6,700 species, representing ∼35 % of all eukaryotic orders. However, despite the size of the training data, it remains small compared with the total chemical and biological diversity. Indeed, millions of species and hundreds of thousands of chemicals remain toxicologically uncharacterized, and within represented eukaryotic groups, coverage is highly uneven (**Fig. 1a, Supplementary Table S1**). For example, for protists, which are important contributors to global ecosystem function, only 83 out of 60,000 described species have toxicity data (Miao et al., 2020; Perrin & Dorrell, 2024). In contrast, a small number of chemicals and species account for a disproportionate fraction of all data, with, e.g., copper and *D. magna* constituting 4.29 % and 5.21 % of the data, respectively. This reflects fundamental constraints of the generation of ecotoxicological data, including the resources required for testing and the practical and ethical challenges associated with studying diverse species and chemicals (Kristiansson et al., 2021). Here, computational approaches offer tractable alternatives to exposure experiments, circumventing high costs, delays, and ethical issues. As this paper shows, accurate extrapolations over larger taxonomic distances enable toxicity assessment beyond current experimental evidence. However, it should be noted that the performance estimates will remain unknown (and likely reduced) for parts of the taxonomic tree where experimental data is completely lacking.

TRIDENT-2 can efficiently use information from data-rich species, chemicals, and exposure scenarios to improve predictions for data-poor combinations thereof. This enables novel experimental testing strategies aimed, for instance, at improving broad-scale toxicity assessment accuracy rather than optimizing individual chemicals. The results from TRIDENT-2 can pinpoint the experimental combinations likely to have the greatest impact, enabling the research community to target the most critical knowledge gaps. For example, the comparatively poor extrapolation across insect orders (**Fig. 3f-g**) suggests that expanding toxicity data within this part of the tree of life is likely to yield greater overall improvements compared to continued testing of already well-studied vertebrate groups. We, therefore, argue that a process where experimental and computational approaches are used in tandem would have significant advantages, enabling more efficient and sustainable generation of toxicological knowledge.

### Modeling toxicity is a multimodal problem

An adverse toxicological response depends on multiple factors, e.g. the chemical properties, the biochemical and physiological makeup of the exposed species, and the exposure conditions, including route of administration and duration. Accordingly, TRIDENT-2 is designed to be multimodal, and data from several sources is fused into a single representation of toxicity. We deliberately retained exposure durations, routes of administration, organism life stages, and toxicological effects, together with chemically and taxonomically diverse data to capture as much biological complexity as possible. Although some exposure combinations were sparse, with modest contributions to the training data, including additional data typically improved the predictive performance across all groups of species. This highlights the value of allowing models to train on diverse data to increase performance and implies a large benefit to sustainable development if diverse AI-ready toxicity data would be made available.

Our species representation and relationships were based primarily on taxonomic distance. While previous studies have shown that taxonomy alone is a poor predictor of toxicity, this is highly chemical-dependent and has never been analyzed on this scale (Coleman & Edmands, 2024). TRIDENT-2 incorporates taxonomy through its multimodal attention mechanism, where the model can choose how much of that signal it will use to guide a prediction. Thus, as our results clearly show, it enables extrapolation of information between species groups. This representation, however, does not fully capture the physiological and ecological differences that influence species-specific sensitivity to chemicals. It is, therefore, likely that incorporating additional biological information, such as species’ ecological, physiological, and morphological traits, would further improve toxicity predictions. However, such data are currently only available for a small fraction of the eukaryotic diversity (Yang et al., 2026). As data continues to expand, multimodal AI provides a natural way to incorporate more diverse information that will further improve accuracy and provide increased toxicological understanding.

To conclude, multimodal AI makes it possible to efficiently use available experimental toxicity data to bridge fundamental data gaps and accelerate a needed transition towards more informed chemical risk assessments, effectively moving from assessments based on standard test species to assessments based on ecological relevance. We argue that this new capability can lead to better decisions in chemical management and sustainability, including but not limited to, better prioritization for chemical substitution, increased vigilance towards chemicals with previously unidentified toxicity, and more rapid identification of chemical impacts. Ultimately, only by taking better, more informed decisions will we be able to avert the planetary crisis and ensure that we stay within the planetary boundaries.

## Methods

### Data acquisition, preprocessing, and representation

#### Data sources and initial processing

We compiled toxicity data from the Registry of Toxic Effects of Chemical Substances (RTECS), REACH dossiers (Registration, Evaluation, Authorization and Restriction of Chemicals), the U.S. Environmental Protection Agency ECOTOXicology Knowledgebase (ECOTOX), and EFSAs OpenFoodTox (European Food Safety Authority). RTECS data was downloaded in November 2021 and, after reformatting, contained 390,976 entries. The REACH dossiers were accessed in June 2023 tabulated into 279,012 individual study entries. The US EPA ECOTOX database was downloaded in May 2023 and contained 1.15 million entries. Data on pesticide registration data was downloaded from EFSA in March 2024 and contained 9,633 entries.

Endpoints, effects, routes of administration, and organism life stages were harmonized (**Supplementary Tables S6-12**). Toxicological endpoints were filtered to include no observed effect concentrations (NOEC), lowest observed effect concentration (LOEC), and 50% and 10% effect concentrations (EC_50_, EC_10_). Toxicological effects were mapped to 10 words: mortality (MOR), developmental toxicity (DVP), behavioral effects (BEH), reproductive toxicity (REP), growth effects (GRO), intoxication (ITX), physiological effects (PHY), morphological effects (MPH), population-level effects (POP), and carcinogenicity (CAR). Concentration units and concentrations were translated and filtered to include 14 possible values (in the order of frequency: mg/L, mg/kg, kg/m², %, ppm, mg/kg bw, mg/kg soil, mg/org, mg/L air, mg/kg diet, mg/kg org, mL/kg bw, mg/org/d, mg/kg bw/d). Exposure durations were converted to hours (h) using 1e-6h for missing values. Routes of administration were harmonized to 23 possible values (in the order of frequency: static, oral, renewal, intraperitoneal, environmental, flow through, spray, intravenous, food, direct application, subcutaneous, gavage, inhalation, culture media, granular, topical, intramuscular, soaking, dermal, drinking, in vitro, injection, parenteral). Life stages were similarly mapped to early, mid, and adult. Concentration or dose limit values and values falling outside the range 0-1e10 were removed. Concentrations and duration values were log_10_ transformed. Finally, duplicate experiments—defined as having identical chemical structure, species, life stage, route of administration, endpoint, effect type, concentration unit, concentration value, duration unit, and duration value—were consolidated. The resulting dataset comprised 560,780 observations.

#### Chemical descriptors

Chemical Abstract Service (CAS) registry numbers were validated and formatted to ensure consistency. SMILES (Simplified Molecular-Input Line-Entry System) were mapped through a multi-tiered approach including PubChem, PubMed and Chemical Identifier Resolver (CIR). All SMILES were canonicalized using *rdkit* v2025.5.2, removing stereochemical information. Missing, invalid or SMILES causing parsing errors were removed. All salts, hydrated-, metal-, and organometal SMILES were kept unaltered.

#### Species information and taxonomic annotation

Species information was semi-manually harmonized, using common names, scientific names, and ECOTOX group assignments. Entries where no species name could be identified or where the name was provided as an ECOTOX group name were either removed or represented at a higher taxonomic level. Entries reporting multiple species were expanded into separate records, with each species receiving its own entry while retaining all other experimental parameters. Missing species group assignments were imputed using species-to-group mappings.

Taxonomic information was retrieved from the NCBI Taxonomy database (accessed 2025-09-23) using *ete3 toolkit* v3.1.3. Species scientific names were first fuzzy matched against NCBI scientific names, followed by genus-level matches to retrieve NCBI taxonomic IDs. When available, species groups were used as additional context to validate matches and resolve ambiguities. Matched species were verified to ensure they belonged to *Eukaryota,* except for cyanobacteria which were retained as they are a common model organisms.

Complete taxonomic lineages were retrieved for all species, spanning nine hierarchical ranks: superkingdom, kingdom, phylum, subphylum, class, order, family, genus, and species. Subphylum was included as it divides certain conventional eukaryotic groups, such as crustaceans. Missing intermediary ranks were filled using a top-down (leaf to root) approach, propagating the ancestor downwards with a suffix. Entries without species or genus ancestral information were removed.

#### Taxonomic tree and taxonomic embeddings

A taxonomic tree was constructed incorporating all species using a branch length of one. Pairwise taxonomic distances between node pairs in the tree were calculated by taking the sum of branch lengths along the path connecting two nodes. The distance matrix was transformed into continuous vector representations using metric multidimensional scaling (MDS) executed for 10,000 iterations on a NVIDIA A100-SXM4-40GB GPU using *mdscuda* v0.1.3. This embeds each taxonomic ID into a 768-dimensional Euclidean space while preserving pairwise distances as faithfully as possible. To decrease sparseness in the Euclidian space and improve model understanding of node connections, we also optimized distances to intermediary nodes. Thus, MDS embeddings for all nodes in the tree, including lower taxonomic ranks, were obtained. Embedding quality was assessed by comparing taxonomic to Euclidean distances (**Supplementary Fig. S19**).

### Model architecture

The model predicts the four toxicological endpoints (EC_50_, EC_10_, LOEC, NOEC) simultaneously, based on chemical structure, target species, route of administration, organism life stage, exposure duration, concentration unit, and target toxicological effect. The model is comprised of three modules: an embedding module, a multimodal transformer, and a multitask neural network.

The embedding module embeds the chemical structure, taxa, route of administration, organism life stage, and toxicological effect into 768-dimensional vectors. Chemical structures, represented by a SMILES strings, are tokenized using a byte pair encoding (BPE) tokenizer and embedded using pretrained ChemBERTa-1 embeddings and absolute positional encoding (Chithrananda et al., 2020). Taxonomic embeddings are reprojected and normalized using a learnable transformation and LayerNorm layer and prepended to the SMILES token embeddings. Routes of administration, organism life stages, and toxicological effects are represented by individual tokens, embedded using learnable embeddings, and prepended to the SMILES embeddings. When route of administration or organism life stage is missing in the data, a unique “missing” token is used. A beginning-of-string (CLS) token is embedded and prepended to the input embeddings and all modalities are separated by a separation (SEP) token.

The multimodal transformer is a six-encoder layer RoBERTa transformer which applies self-attention to all input embeddings (Liu et al., 2019). We initialize the RoBERTa encoders with ChemBERTa-1 pretrained weights. ChemBERTa-1 was chosen as the backbone of the transformer after initial fine-tuning tests comparing ChemBERTa-1 (Chithrananda et al., 2020), ChemBERTa-2 (Ahmad et al., 2022), Molformer (Ross et al., 2022) and Molecule Attention Transformer (Maziarka et al., 2020), showed it having the best performance.

The CLS-embedding from the multimodal transformer is concatenated with the exposure duration, a one-hot encoding of the concentration unit and duration missingness flag and subsequently passed to a multitask neural network. The neural network has one hidden layer and four prediction heads that predict the individual toxicological endpoints. We use a dropout of 0.2 through the entire model and GELU activation functions in the neural network.

### Model training and validation

All models were built using PyTorch v2.1.2 and Huggingface transformers v4.39.3. Models were trained using mixed precision, evaluated in full precision and monitored using weights and biases (wandb) v0.17.9. All code was written in python v3.11.

#### Data splits and stratified sampling

To estimate the performance on untested chemical structures, ten-fold cross-validation based on Butina clusters was used. SMILES were encoded as Morgan fingerprints of size 1024 with radius 3 and clustered using rdkit Butina clustering (iterative cluster assignment using a Tanimoto dissimilarity cutoff). Entire clusters are randomly assigned to either the training or validation set to prevent data leakage. This was repeated for successive Butina cutoffs (0.2-0.9) to estimate errors across increasing chemical dissimilarity. To estimate the performance of untested species, we similarly implemented ten-fold cross-validation with unique species in each fold, ensuring no species overlap between training and validation sets. Here, we also separated species by higher taxonomic ranks (genus, family, order, class, phylum, and kingdom) to obtain performance estimates with increasing taxonomic distance. Finally, to estimate the predictive performance where partial experimental data has already been collected, we constructed ten-fold cross validation scenarios where either the Butina cluster or the species was included in the training data but not in combination.

#### Hyperparameter optimization

Hyperparameters were optimized using Bayesian optimization and grid-search. Candidate values were based on previous results (Gustavsson et al., 2024) and initial tests. Batch size and learning rate were tuned using Bayesian optimization for a random 80/20 train/validation split across 50 epochs over 72 iterations using wandb with hyperband early termination (Biewald, 2020) evaluated after a minimum of 15 epochs.

Grid-search was run on the ten best performing combinations from the Bayesian optimization using a five-fold cross-validation. Based on the mean and standard deviation of the folds we selected; batch size 256, learning rate 5e-5, and 100 epochs for training of subsequent models. Early tests determined that the impact of partially reinitializing the pretrained encoders, applying layer wise learning rate decay, and increasing the weight decay (L2 regularization) of the AdamW optimizer (Loshchilov & Hutter, 2019) had negligible impact on performance.

#### Loss function, regularization, sampling, and data augmentation

To optimize model parameters, we use a combination of a Mean Absolute Error loss function (L1Error) and two regularization terms (**equations -3**). Mean absolute error was chosen to reduce the impact of outliers. The first regularization term enforces predictions to follow a monotonous behavior, where LOEC and EC_50_ are always predicted as higher than NOEC and EC_10_ respectively. The second regularization term promotes evolutionarily similar species to respond similarly to specific chemicals, aggregating higher taxonomic ranks toward the mean of their descendant species. This encourages species to learn from their close relatives and therefore mitigates data sparsity for poorly represented species. The full loss function is given by:

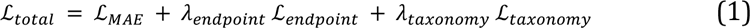

where ℒ*_endpoint_* is given by:

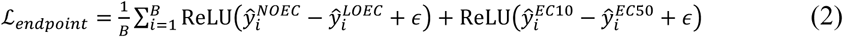

^and ℒ^*_taxonomy_*:

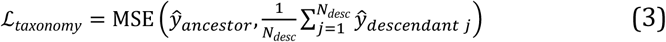

where *λ_endpoint_* and *λ_taxonomy_* are weighting hyperparameters for the two regularization terms (0.1 and 0.05 respectively), *B* is the batch size, *ε* is a small margin (10⁻³), *^y* the predicted log_10_-concentration for each endpoint, and *N_desc_* is the number of descendant species. We use the AdamW optimizer with default settings for gradient optimization, with a linear learning rate scheduler with 10% warmup steps and 90 % cool down steps.

To prevent the large skews in data coverage to bias the training, stratified sampling was implemented through a weighted sampler, inversely proportional to the square root of the number of SMILES, eukaryotic group, effect, and endpoint counts. Additionally, data augmentation was applied during sampling by randomly enumerating SMILES (i.e., not only the canonical form was not used in the training data) and by randomly masking route of administration, life stage, and effect with an independent probability of 0.1 for every sample.

#### Performance calculation

To assess model performance, predictions were evaluated using a hierarchical aggregation procedure. Technical replicates were first summarized by taking the median for each unique experimental setup defined by endpoint, effect, chemical, taxonomic rank, exposure duration, administration route, and organism life stage. Absolute error was then calculated for each setup, followed by averaging across exposure durations to obtain one error per endpoint–effect–chemical–species combination. For effect-specific analyses, this level of aggregation was retained. For analyses across effects, errors were subsequently combined using a weighted mean, with weights proportional to the number of observations for each endpoint-effect-chemical-species combination, yielding one error per endpoint-chemical-species combination.

To evaluate performance across species and chemicals while controlling their relative contributions, errors were further aggregated within species groups using three complementary approaches. Macro-averaging assigned equal weight to each chemical–species combination, whereas taxon-averaging first averaged across species and was used to assess performance for novel species; chemical-averaging first averaged across chemicals and was used to assess performance for novel chemicals. Median absolute error (MAE) was calculated from the resulting distributions, with each averaging strategy defining the unit receiving equal weight. Variability was quantified using the median absolute deviation standard error (MAD-SE), calculated as MAD divided by the square root of the corresponding sample size (see **Supplementary Text 1** for details).

#### Benchmarking

We compared TRIDENT-2 against three common QSAR methods—ECOSAR v.2.2, VEGA v1.2.3, and T.E.S.T. v5.1.1 (Martin, 2020; Roncaglioni et al., 2022; Wright et al., 2022)—a transformer-based model (TRIDENT-1) (Gustavsson et al., 2024), and a pairwise learning model (Posthuma et al., 2025). As only TRIDENT-2 could produce LOEC predictions, we only compared EC_50_ and pooled EC_10_ and NOEC predictions to retain sufficient data. For ECOSAR, VEGA, and T.E.S.T., we excluded all training and experimental data and transformed NOEC predictions into EC_10_ to make performance comparisons for untested chemicals (see **Supplementary Text 2-3** and **Supplementary Table S13**). To not overestimate errors for ECOSAR, VEGA and T.E.S.T. we only compared errors reported to be inside their respective applicability domains. For TRIDENT-1, we used cross-validation results for untested chemicals from the corresponding study and predicted chemicals in the dataset to estimate predictability. For Posthuma et al. 2025, we used cross-validation results from the corresponding study and executed the exact same cross-validation with TRIDENT-2. To compare species and eukaryotic order coverage, we strictly matched species scientific names to the dataset, and map species to extant eukaryotic orders in the NCBI Taxonomy database (n=1,455, accessed May 2026).

### Predicting pesticide toxicity across species

SMILES were gathered from the Pesticides Properties Database (PPDB) and Biopesticide Database (BPDB) (Tzilivakis et al., 2026) (accessed 2026-05-22), resulting in 2,146 SMILES. Species were assigned harmonized combinations of toxicological effect, concentration unit, and exposure duration based on eukaryotic group, habitat and frequency in the data which were used to predict toxicity for all species in the dataset (**Supplementary Text 4, Supplementary Table S14-S17**). To make predictions comparable across species, effects, durations, and concentration units, we also predicted toxicity for the 12,890 chemicals in the dataset that did not solely have rodent data. In total this resulted in 92,316,870 predictions. The median predicted EC_50_ per species was subtracted from the pesticide predictions, generating normalized EC_50_ values (**Supplementary Fig. S17**). To illustrate predictions in a heatmap, we averaged species predictions to the taxonomic order level and clustered pesticides using Pearson correlation and average linking. Finally, normalized predictions were categorized into three hazard levels (low, mid, high) based on the number of standard deviations from the mean of all predictions.

## Supporting information

Tabulated figures

## Data and code availability

### Data and materials availability

The code and data used to build, train, and evaluate the models presented in this study are available at https://github.com/StyrbjornKall/TRIDENT_2 and 10.5281/zenodo.22097847. The RTECS dataset is commercially available from BIOVIA for a fee or by way of subscription and cannot be shared. TRIDENT-2 is also provided through an online service at https://trident.serve.scilifelab.se/.

## Acknowledgements

**Acknowledgments:** the computations were enabled by resources provided by the national Academic infrastructure for Supercomputing in Sweden (NAISS) and the Swedish national infrastructure for computing (SNIC).

Funding

This work was supported by the SciLifeLab & Wallenberg Data Driven Life Science Program (grant: KAW2020.0239) (S.K., E.K., and J.H.) and by the FRAM centre for Future chemical Risk Assessment and Management at the University of Gothenburg (to S.K., M.G., P.S., and E.K.) and A Swedish Research Council for Sustainable Development (FORMAS) (M.G. grant: 2022-00528, E.K. grant: 2024-02047).

## Author information

### Contributions

**Author contributions:** conceptualization: S.K., M.G., and E.K. Methodology: S.K., M.G., and E.K. Validation: S.K., M.G., and E.K. Software: S.K., M.G., and P.S. Formal analysis: S.K., M.G. P.S., M.D., T.B., and E.K. Funding acquisition: S.K., M.G., and E.K. Data curation: S.K., M.G. and P.S. Investigation: S.K., M.G. and E.K. Supervision: E.K. and M.G. Project administration: S.K., M.G., and E.K. Writing—original draft: S.K., M.G., and E.K. Writing—review and editing: S.K., M.G., P.S., J.H., T.B., and E.K.

## Supplementary Materials

This PDF file includes:

Supplementary Results

Supplementary Methods

Supplementary Text 1 to Supplementary Text 4

Figures S1 to S19

Tables S2 to S17

Caption for Table S1

