## Supplementary Information for "Assessing chemical toxicity across Eukaryota using multimodal transformers"

**This PDF file includes:**

Supplementary Results

Supplementary Methods

Supplementary Text 1 to Supplementary Text 4

Figures S1 to S19

Tables S2 to S17

Caption for Table S1

### 1 Supplementary Results

**Table S1. Summary of ecotoxicological data used in the study, stratified by eukaryotic group and toxicity endpoint.** For each combination of eukaryotic group (including an aggregate "all taxa" category) and endpoint (EC50, EC10, LOEC, NOEC), the table reports: the number of unique species, genera, families, orders, classes, phyla, and kingdoms represented within each group; the toxic effect(s) captured (e.g., mortality [MOR], population [POP], morphology [MPH]); the number of observations (rows); the number of unique experimental setups (unique combinations of effect, chemical [SMILES], species, exposure duration, route of administration, and organism life stage); the number of unique chemicals tested (based on SMILES); the number of unique routes of administration and organism life stages; the number of distinct concentration units reported; mean ± standard error of the mean (SEM) for exposure concentration, computed separately for each concentration unit that had at least 10 observations and was reported consistently across all endpoints within the group; and mean ± SEM for exposure duration.

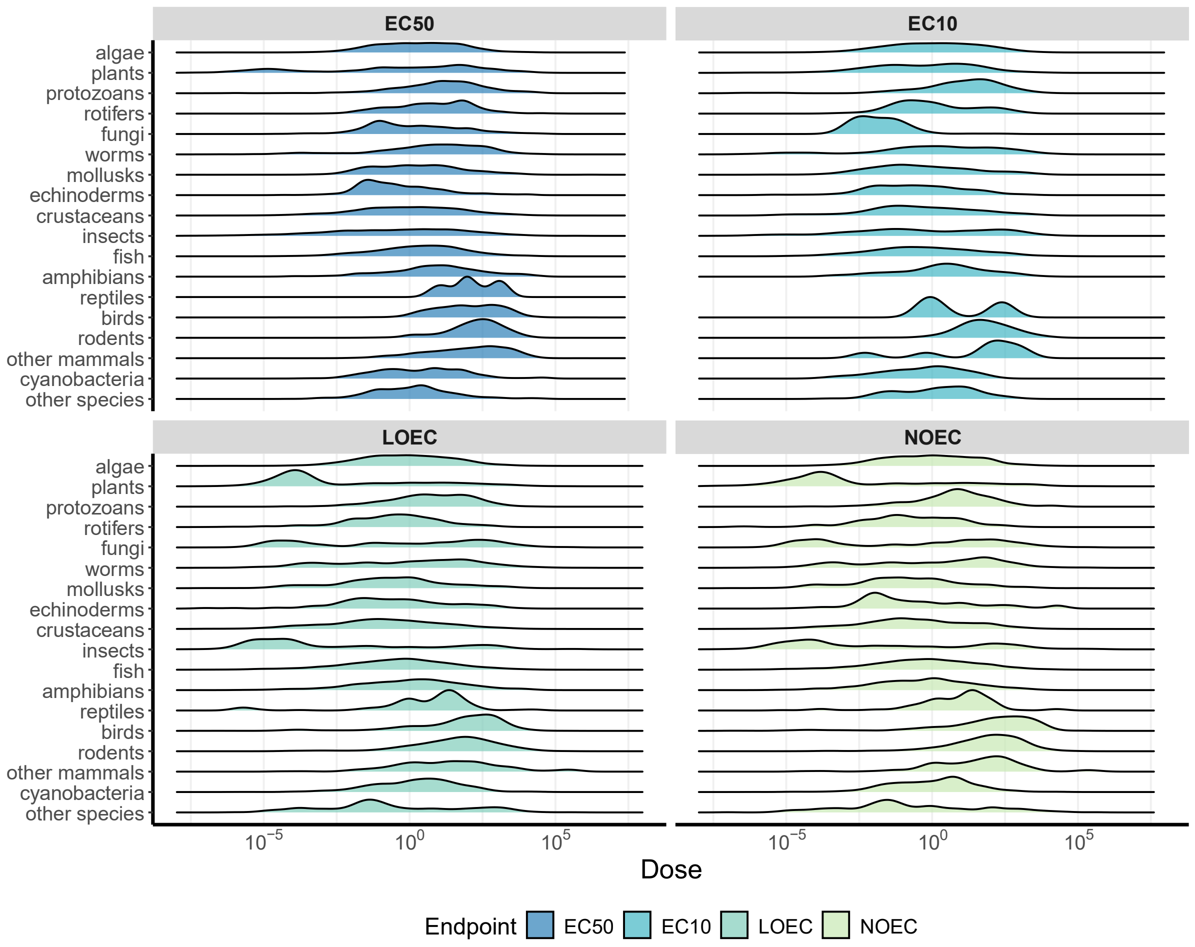

**Figure S1. Endpoint concentration profiles by eukaryotic group.** Ridgelines indicate the distribution of EC50 (n=319,528), EC10 (n=15,158), LOEC (n= 91,706), and NOEC (n=134,403) across the eukaryotic groups.

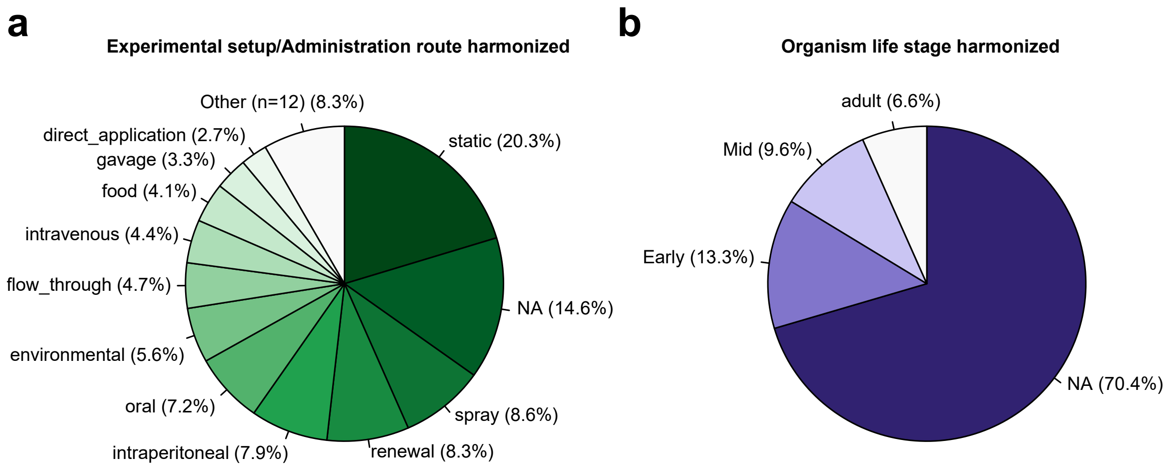

**Figure S2. Distribution of (a) experimental setups/routes of administration and (b) organism life stages after harmonization.** Categories making up less than 5 % of the total observations were grouped into "Other". "NA" indicates missing information.

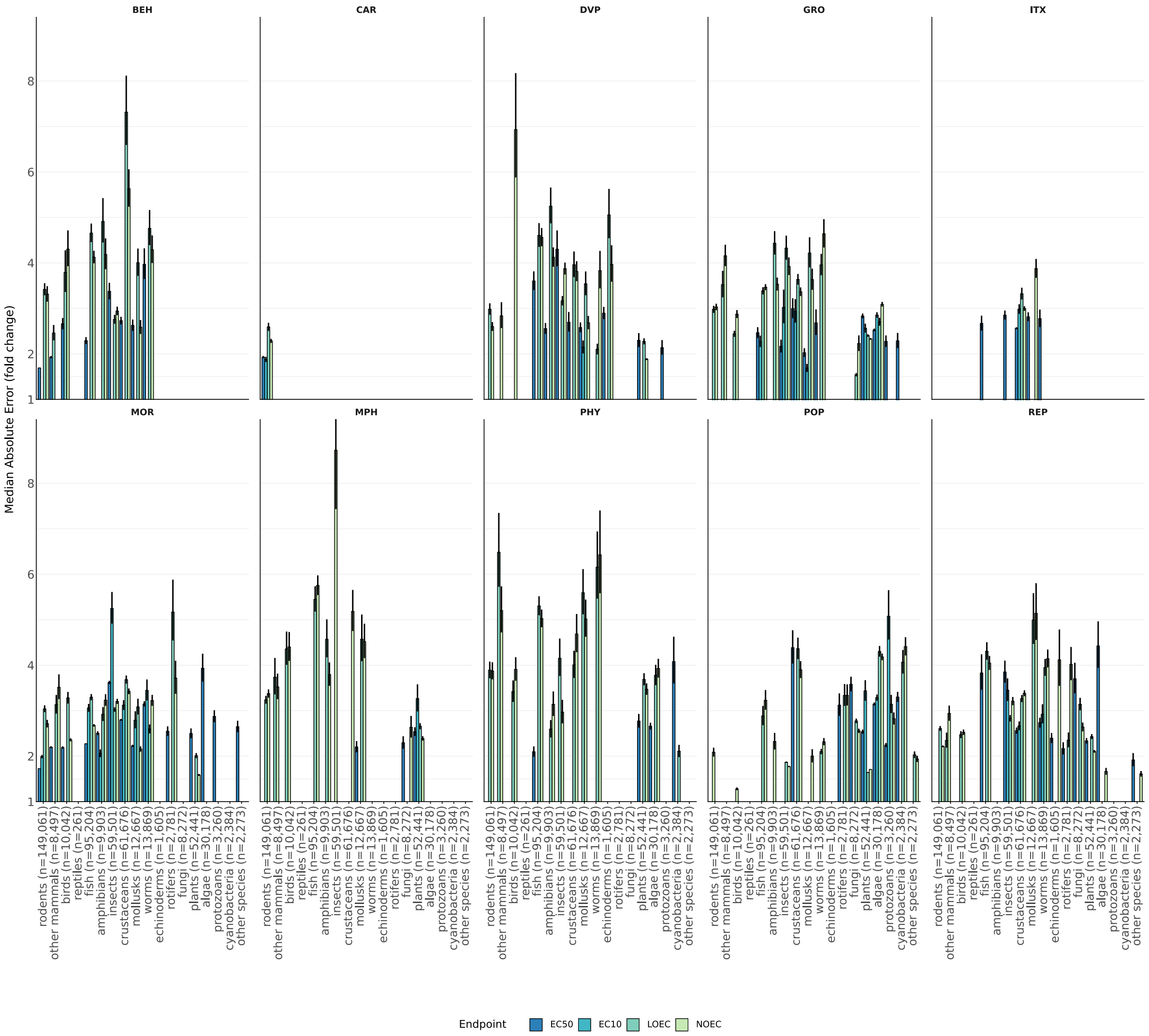

**Figure S3. Stratified Median Absolute Error fold change across effects.** Median Absolute Error (fold change) (MAE) from ten-fold cross-validations with error bars reflecting the Median Absolute Deviation (MAD) divided by √(n). MAE for EC50, EC10, LOEC and NOEC per toxicological effect when evaluating the model on novel combinations of species and chemicals (at Tanimoto similarity less than 0.8, n denotes the number of observations in each group). All combinations with number of observations below n=50 were removed.

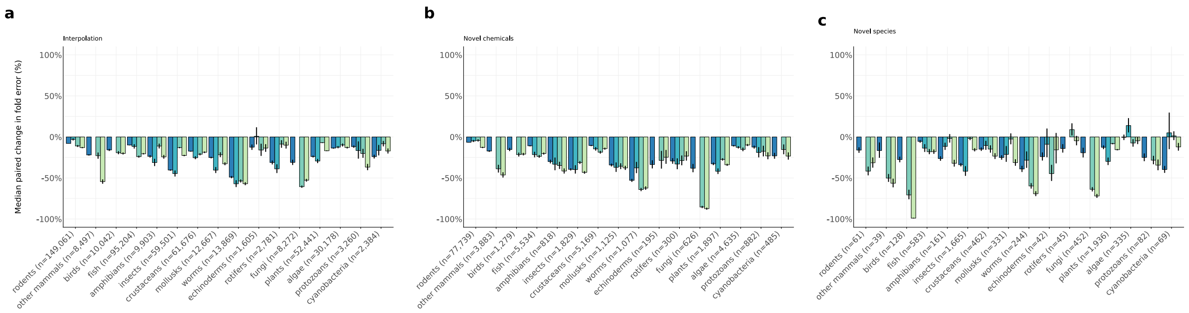

**Figure S4. Percentual change in Median Absolute Error fold change between the model and individual models trained each isolated eukaryotic group.** Values were computed from paired differences in log10 absolute error and back-transformed to the fold-error scale. Negative values indicate lower prediction error for the multimodal model. Error bars represent the back-transformed median absolute deviation (n denotes the number of unique observations in each group). Predictions were obtained from 10-fold cross-validation. **(a)** Percentual change in MAE between models trained on individual species groups and the multimodal model (same as Fig. 3b). **(b)** Percentual change in MAE between models trained on individual species groups and the multimodal model when evaluating the model on unseen chemicals (Tanimoto similarity less than 0.8). **(c)** Percentual change in MAE between models trained on individual species groups and the multimodal model when evaluating the model on unseen species.

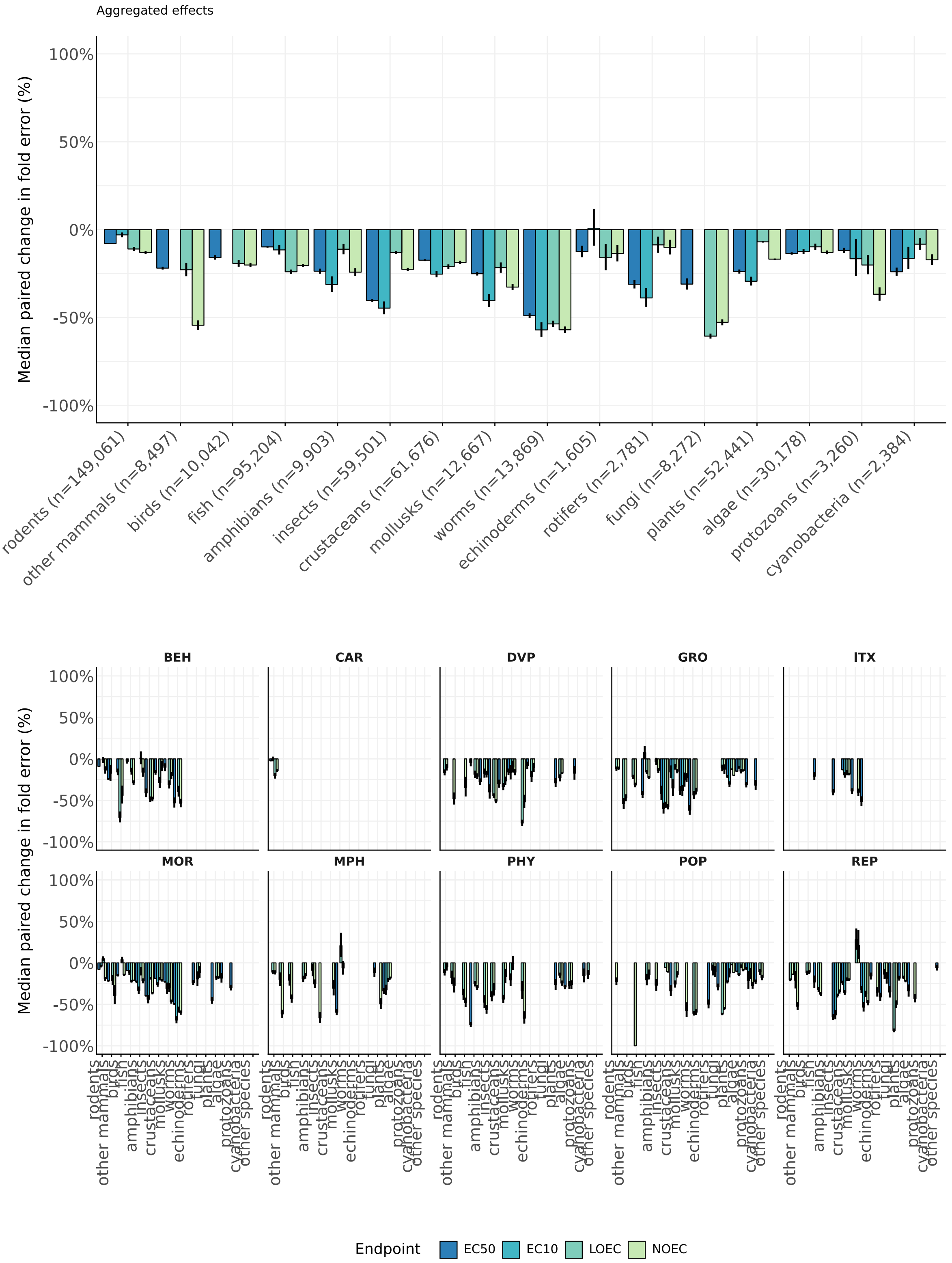

**Figure S5. Stratified percentual change in Median Absolute Error fold change (MAE) between the model and individual models trained each isolated eukaryotic group.** Values were computed from paired differences in log10 absolute error and back-transformed to the fold-error scale. Negative values indicate lower prediction error for the multimodal model. Error bars represent the back-transformed median absolute deviation. Predictions were obtained from 10-fold cross-validation.

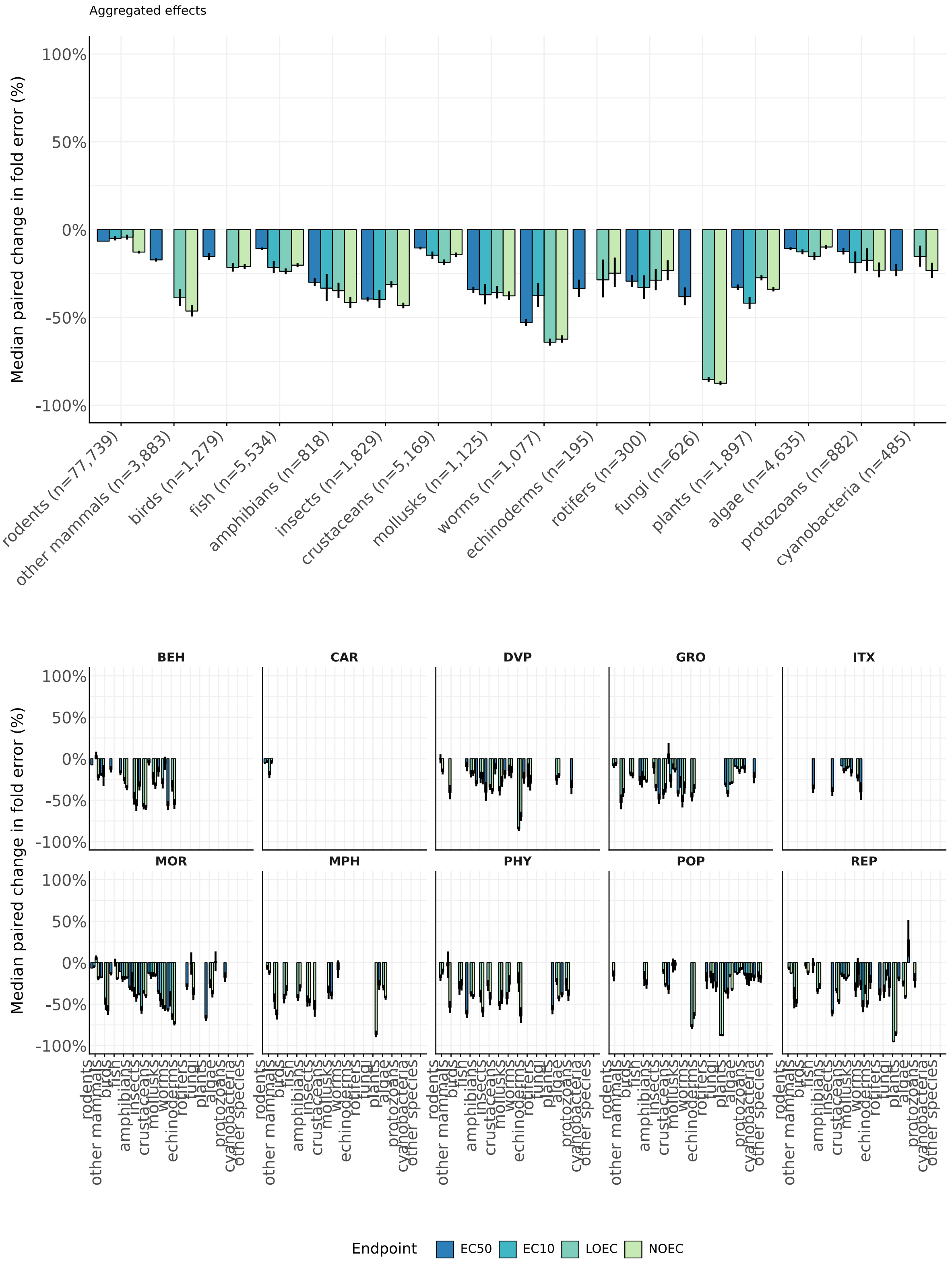

**Figure S6. Stratified percentual change in MAE between the model and individual models trained each isolated eukaryotic group when evaluating models on unseen chemicals.** Values were computed from paired differences in log10 absolute error and back-transformed to the fold-error scale. Negative values indicate lower prediction error for the multimodal model. Error bars represent the back-transformed median absolute deviation. Predictions were obtained from 10-fold cross-validation.

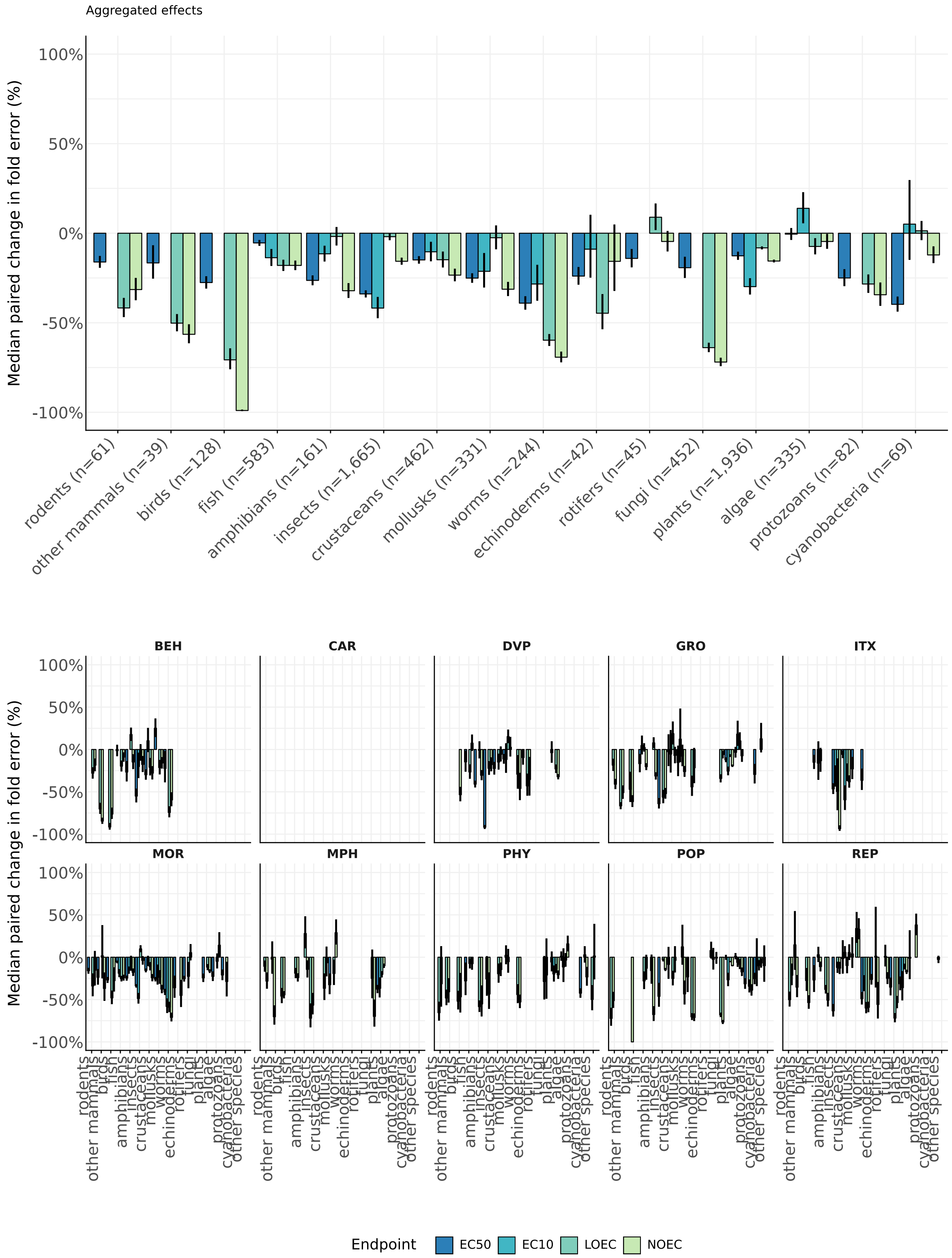

**Figure S7. Stratified percentual change in MAE between the model and individual models trained each isolated eukaryotic group when evaluating models on unseen species.** Values were computed from paired differences in log10 absolute error and back-transformed to the fold-error scale. Negative values indicate lower prediction error for the multimodal model. Error bars represent the back-transformed median absolute deviation. Predictions were obtained from 10-fold cross-validation.

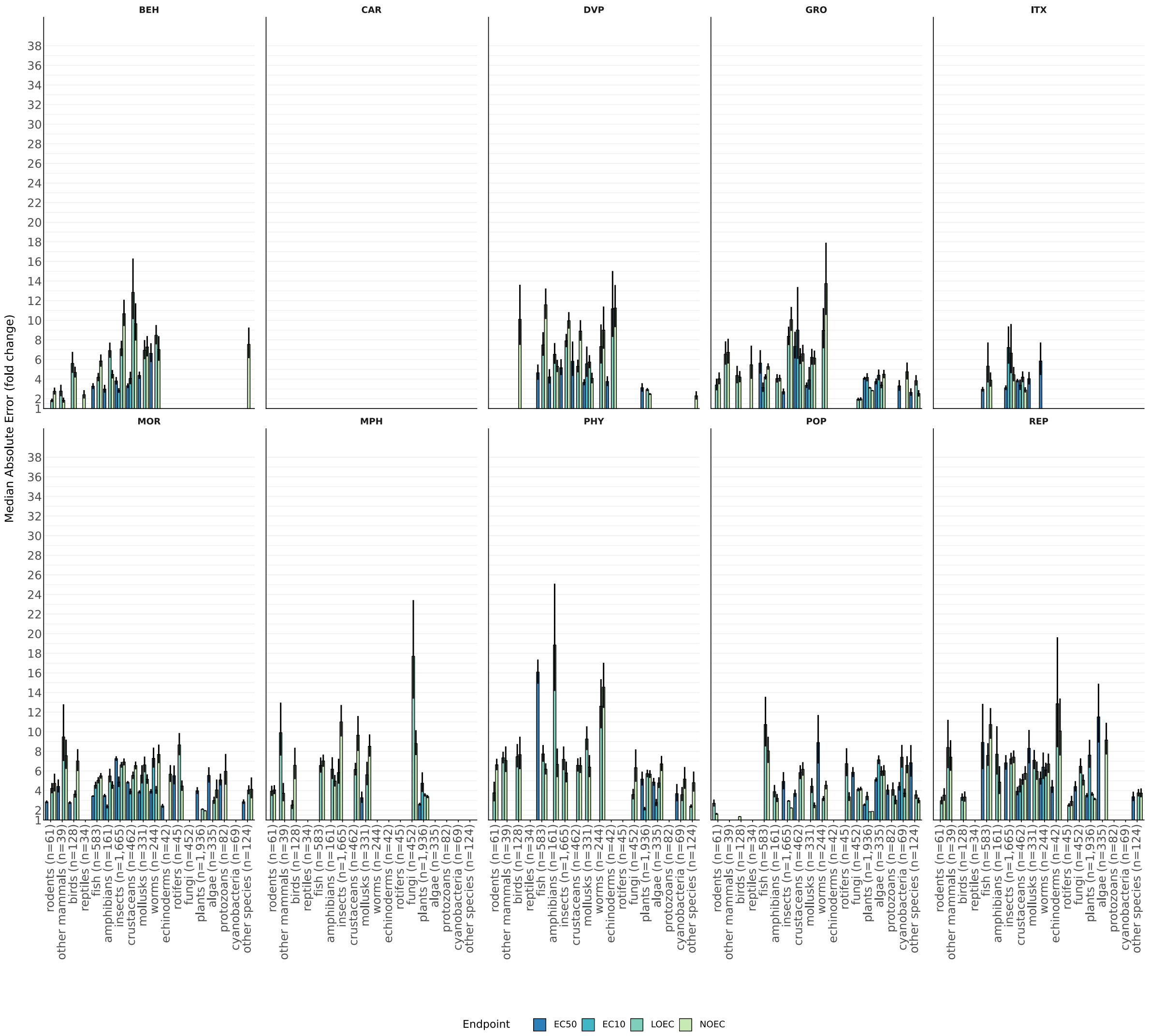

**Figure S8. Stratified Median Absolute Error fold change across effects for unseen species.** Median Absolute Error (fold change) (MAE) from ten-fold cross-validations with error bars reflecting the Median Absolute Deviation (MAD) divided by √(n). MAE for EC50, EC10, LOEC and NOEC per toxicological effect when evaluating the model on unseen species for each eukaryotic group (n denotes the number of unique species in each group). All combinations with number of species below n=10 were removed.

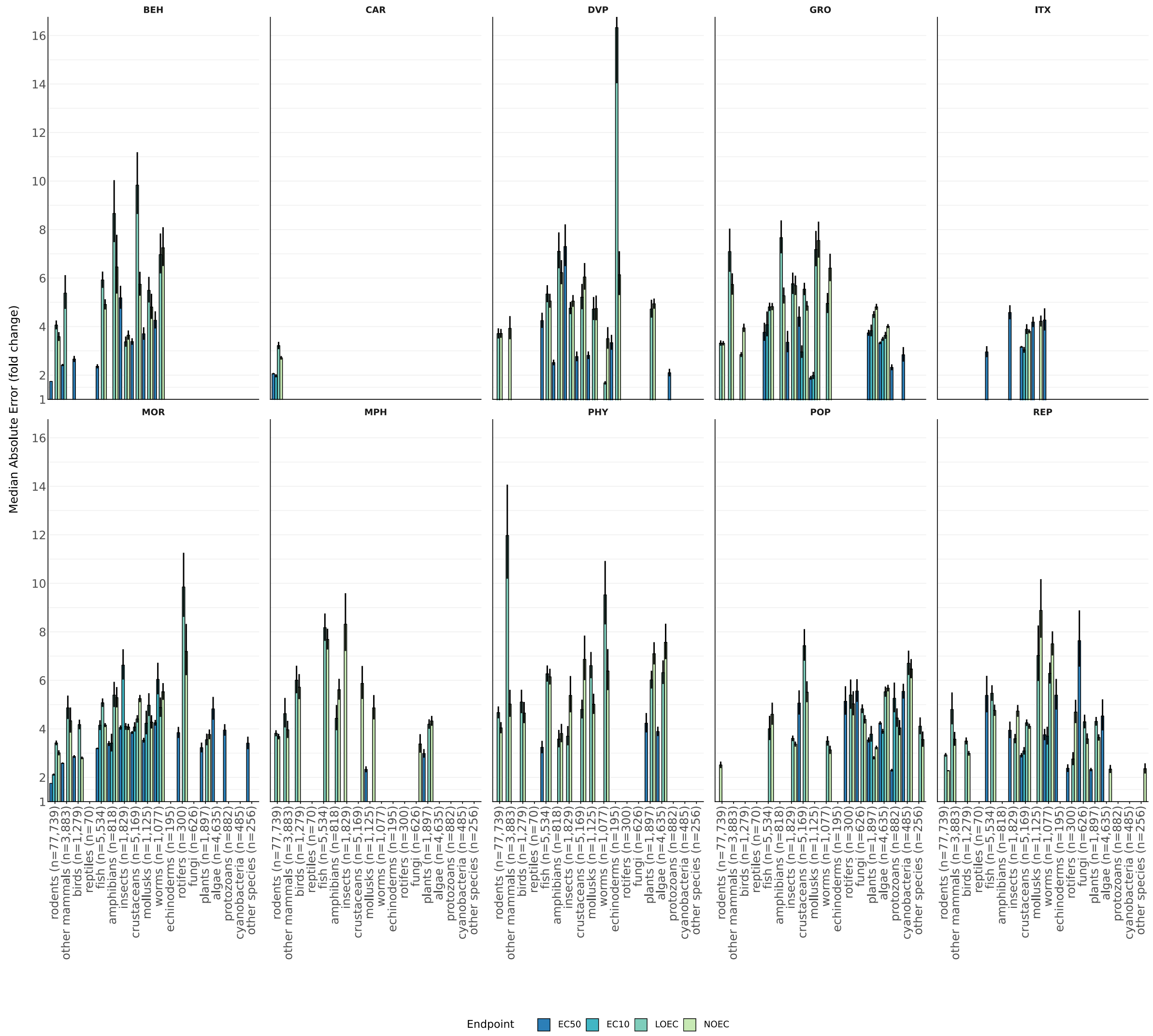

**Figure S9. Stratified Median Absolute Error fold change across effects for unseen chemicals.** Median Absolute Error (fold change) (MAE) from ten-fold cross-validations with error bars reflecting the Median Absolute Deviation (MAD) divided by √(n). MAE for EC50, EC10, LOEC and NOEC per toxicological effect when evaluating the model on unseen chemicals (Tanimoto similarity less than 0.8) for each eukaryotic group (n denotes the number of unique chemicals in each group). All combinations with number of chemicals below n=50 were removed.

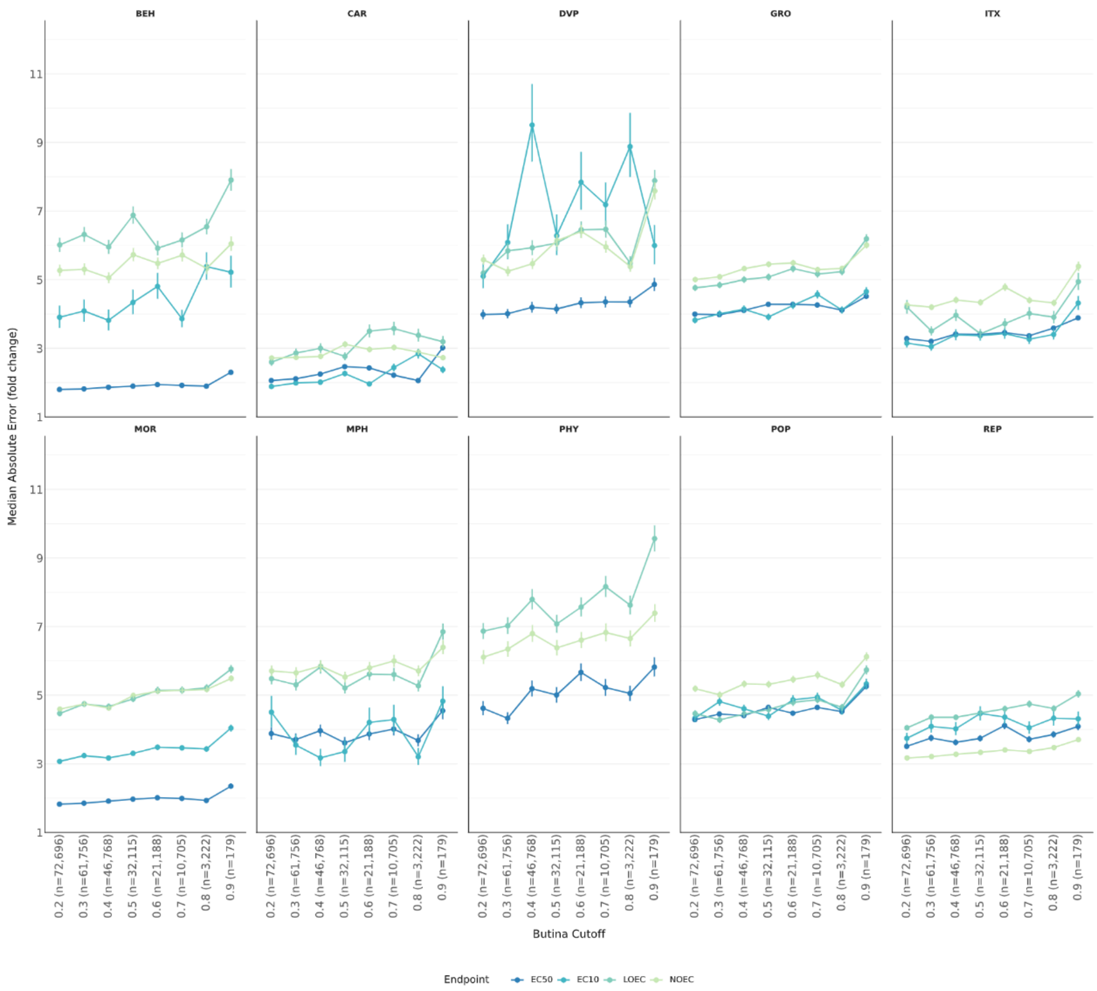

**Figure S10. Stratified Median Absolute Error fold change (MAE) propagation with increasing chemical structural distance.** MAE from ten-fold cross-validations when evaluating the model on unseen chemicals with decreasing Tanimoto similarity/Butina cutoff (n denotes the number of unique Butina clusters across all effects). The large difference in error between EC50-MOR and EC50-BEH and other endpoints is partly due to the large amount of rodent data. Error bars reflecting the Median Absolute Deviation (MAD) divided by √(n).

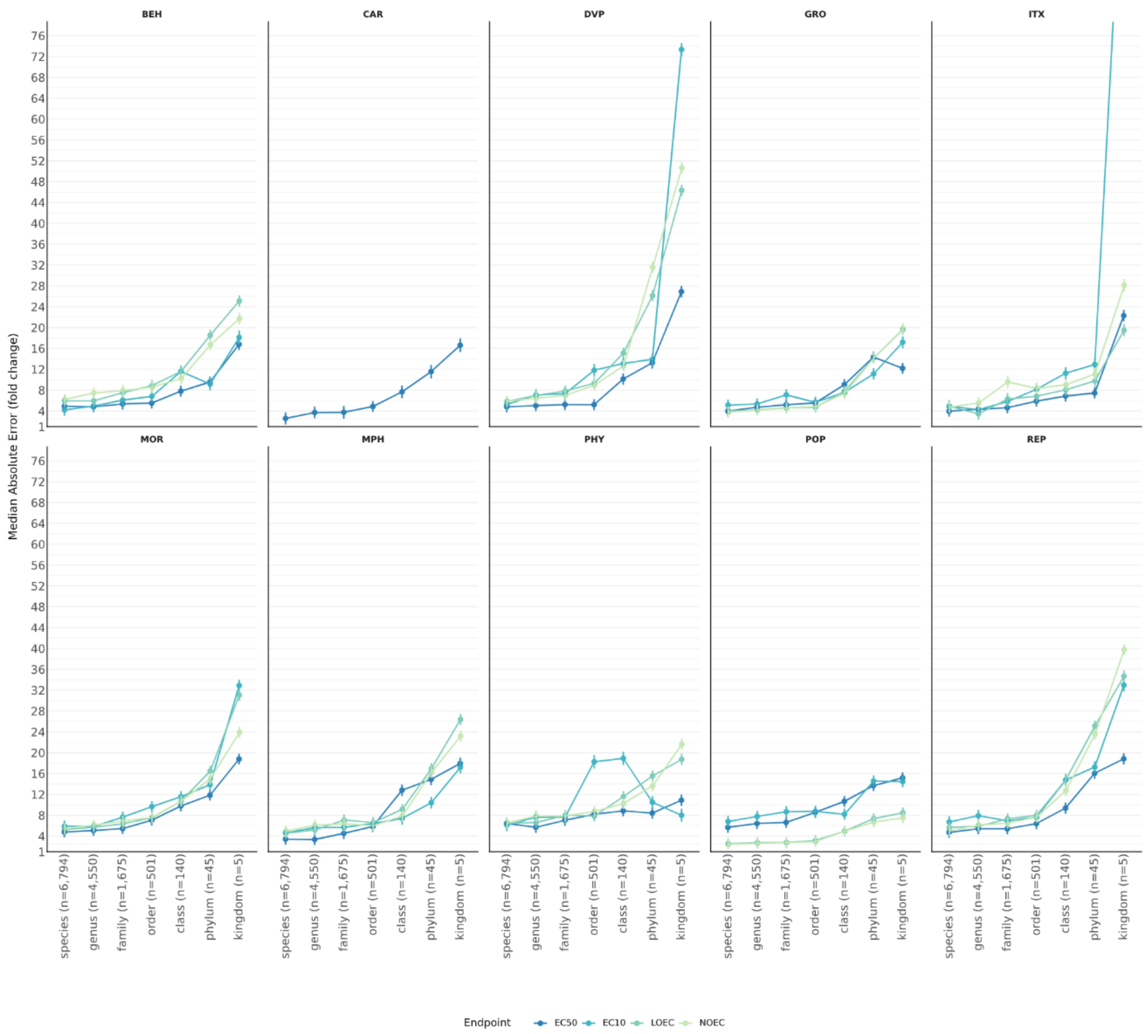

**Figure S11. Stratified Median Absolute Error fold change (MAE) propagation with increasing taxonomic distance.** MAE from ten-fold cross-validations when evaluating the model on species from unseen genera, families, orders, classes, phyla, and kingdoms, respectively (n denotes the number of taxa at each rank across all effects). The first scatter point (species) in each panel indicates MAE for a random subset of unseen species. Error bars reflecting the Median Absolute Deviation (MAD) divided by √(n).

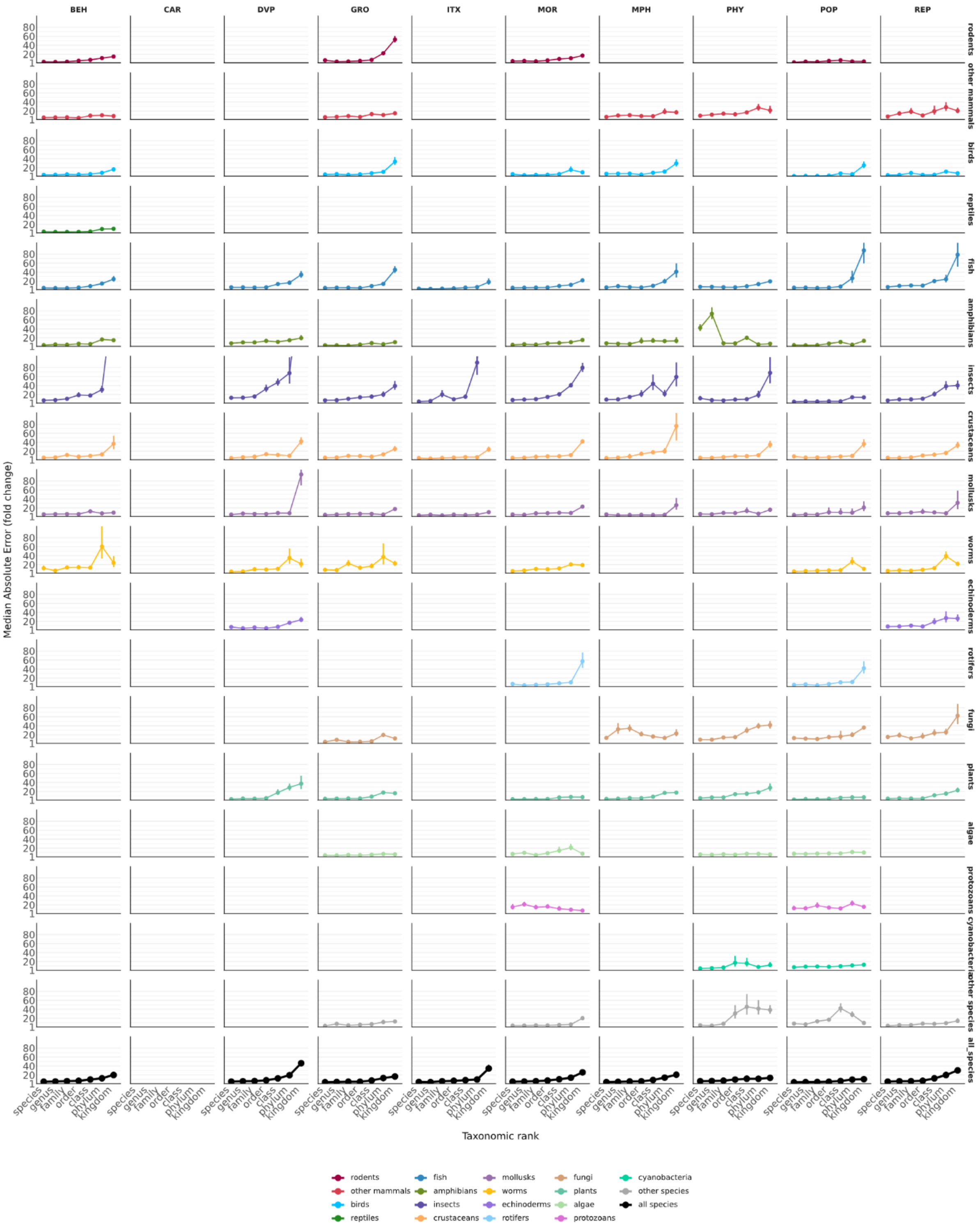

**Figure S12. Median Absolute Error fold change (MAE) propagation with increasing taxonomic distance stratified by eukaryotic group and toxicological effect.** MAE from ten-fold cross-validations when evaluating the model on species from unseen genera, families, orders, classes, phyla, and kingdoms, respectively aggregated across endpoints (EC50, EC10, LOEC, NOEC). The first scatter point (species) in each panel indicates MAE for a random subset of unseen species within each respective group. Error bars reflecting the Median Absolute Deviation (MAD) divided by √(n).

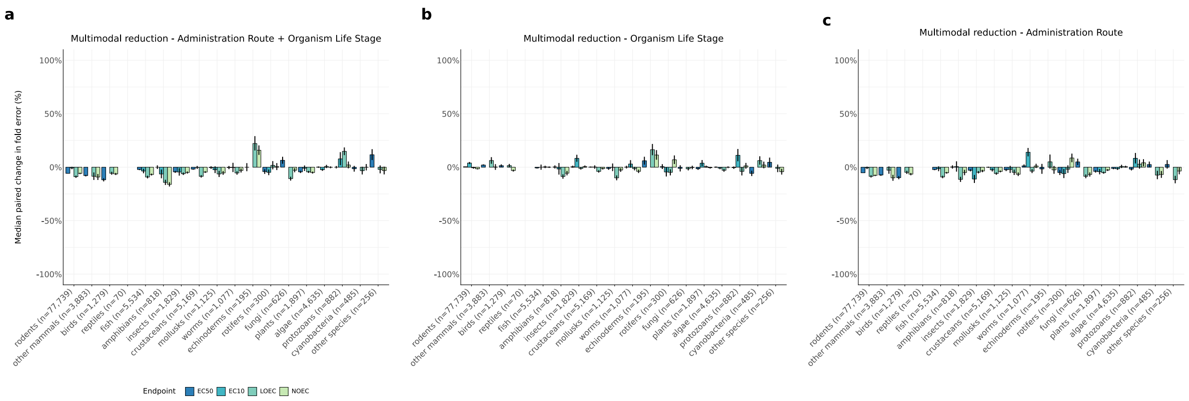

**Figure S13. The impact of organism lifestage and route of administration metadata on model prediction errors for unseen chemicals. (a)** Percentual change in prediction error between the model and a model trained without metadata (organism life stage and administration route). **(b)** Percentual change in prediction error between the model and a model trained without organism life stage. **(c)** Percentual change in prediction error between the model and a model trained without route of administration. Values were computed from paired differences in log10 absolute error and back-transformed to the fold-error scale. Negative values indicate lower prediction error for the multimodal model. Error bars represent the back-transformed median absolute deviation. Predictions were obtained from 10-fold cross-validation.

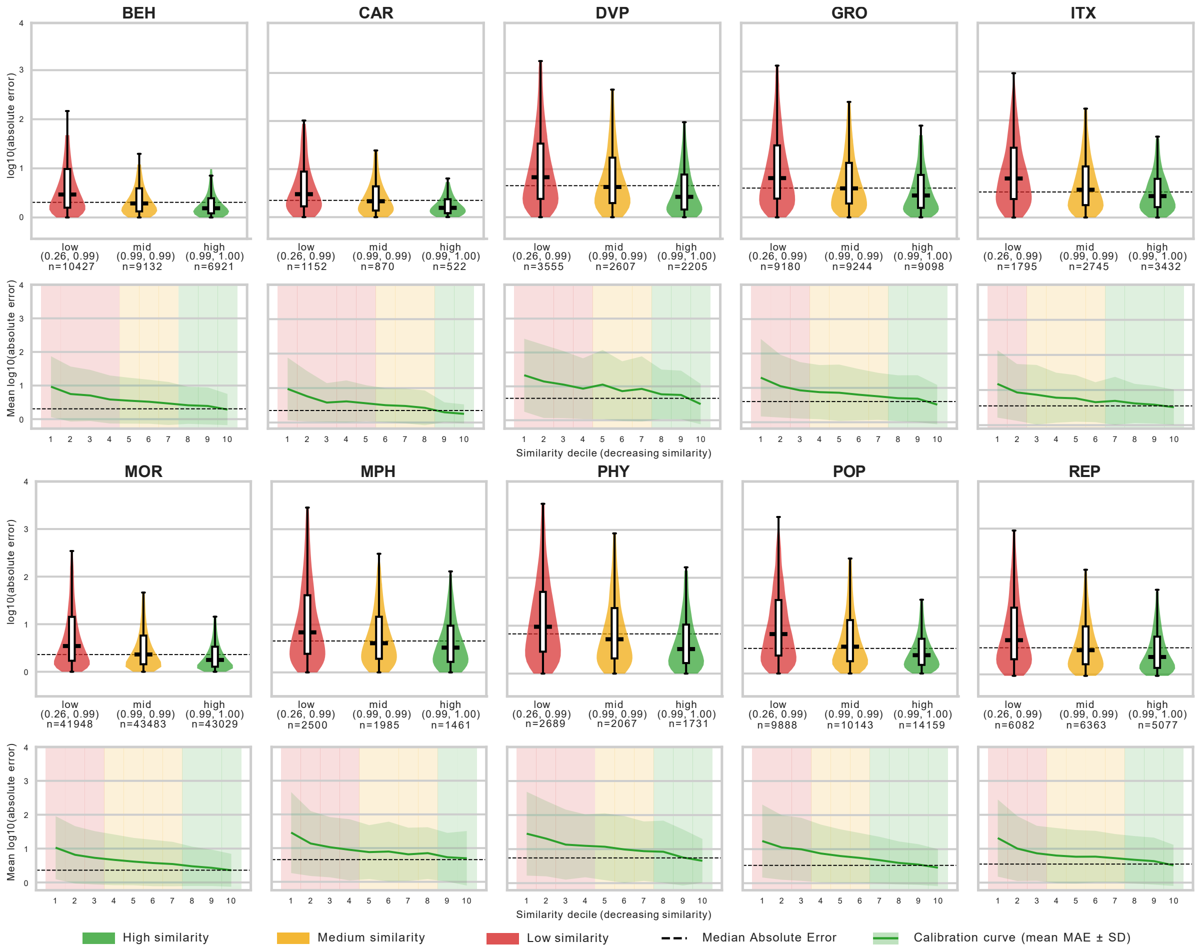

**Figure S14. Median Absolute Error (MAE) distribution with varying cosine similarity to the training data.** Ten-fold cross-validation was used to obtain error estimates for held-out chemicals (Tanimoto similarity less than 0.8) and CLS-embeddings were saved for each validation fold and its respective training set. kNN (k=5) cosine similarity was calculated for each validation entry by taking the mean similarity to its five closest CLS-embeddings in the corresponding training set. Higher values indicate a prediction lies in a region of the embedding space well represented in training data. The 33rd and 67th percentile of the kNN cosine similarity was computed across the whole dataset yielding three similarity intervals (green = [1.00,0.9949), yellow = [0.9949, 0.9872), red = [0.9872, -1], the minimum observed kNN similarity was 0.26). Violins show the distribution of the kNN cosine similarity within each interval for each toxicological effect. Dashed lines mark the mean MAE per effect. Calibration curves show kNN cosine similarity deciles and their associated mean MAE ± standard deviation (SD). Decile background is coloured by which interval its mean similarity falls into. The MAE is on log10 scale.

**Table S2. Pairwise Median Absolute Error fold change (MAE) comparison for EC50 prediction.** Prediction were obtained from each compared model (see Methods, Supplementary Text 2, and Supplementary Table 13) to estimate EC50 MAE for each species/groups yielding predictions for each model. Errors were calculated for all predictable chemicals (same as Fig. 4c) and for chemicals falling inside each tools' respective applicability domain (only ECOSAR and VEGA differ). Bold values indicate lowest error in the comparison.

| Comparison EC50 | Eukaryotic group (n chemicals in comparison) | MAE TRIDENT-2 | MAE other model |
|---|---|---|---|
| **Predictable chemicals** | | | |
| vs TRIDENT-1 | fish (n=3,122) | **2.96 (+0.03, -0.03)** | 3.26 (+0.04, -0.04) |
| vs TRIDENT-1 | crustaceans (n=3,198) | **3.23 (+0.04, -0.04)** | 3.71 (+0.05, -0.05) |
| vs TRIDENT-1 | algae (n=884) | **4.39 (+0.13, -0.13)** | 4.41 (+0.13, -0.13) |
| vs ECOSAR | fish (n=2,341) | **2.78 (+0.04, -0.03)** | 4.05 (+0.08, -0.08) |
| vs ECOSAR | crustaceans (n=2,359) | **2.98 (+0.04, -0.04)** | 5.14 (+0.11, -0.11) |
| vs ECOSAR | algae (n=444) | **4.14 (+0.18, -0.17)** | 7.33 (+0.4, -0.38) |
| vs ECOSAR | worms (n=9) | **3.22 (+1.15, -0.85)** | 81.99 (+40.42, -27.08) |
| vs VEGA | fish (n=2,946) | **2.84 (+0.03, -0.03)** | 5.88 (+0.12, -0.12) |
| vs VEGA | crustaceans (n=2,684) | **3.01 (+0.04, -0.04)** | 10.85 (+0.33, -0.32) |
| vs VEGA | algae (n=1,175) | **3.74 (+0.09, -0.09)** | 8.18 (+0.3, -0.29) |
| vs VEGA | rodents (n=24,332) | **1.75 (+0.0, -0.0)** | 4.43 (+0.03, -0.03) |
| vs TEST | fish (n=2,435) | **2.66 (+0.03, -0.03)** | 3.19 (+0.05, -0.05) |
| vs TEST | crustaceans (n=2,204) | **2.95 (+0.04, -0.04)** | 3.94 (+0.07, -0.07) |
| vs TEST | rodents (n=26,157) | **1.79 (+0.0, -0.0)** | 3.75 (+0.02, -0.02) |
| vs Posthuma | fish (n=1,867) | **2.3 (+0.02, -0.02)** | 2.9 (+0.04, -0.04) |
| vs Posthuma | crustaceans (n=1,411) | **2.7 (+0.04, -0.04)** | 4.59 (+0.12, -0.11) |
| vs Posthuma | algae (n=171) | **3.1 (+0.18, -0.17)** | 5.13 (+0.33, -0.31) |
| **Predictable chemicals inside applicability domain** | | | |
| vs TRIDENT-1 | fish (n=3,122) | **2.96 (+0.03, -0.03)** | 3.26 (+0.04, -0.04) |
| vs TRIDENT-1 | crustaceans (n=3,198) | **3.23 (+0.04, -0.04)** | 3.71 (+0.05, -0.05) |
| vs TRIDENT-1 | algae (n=884) | **4.39 (+0.13, -0.13)** | 4.41 (+0.13, -0.13) |
| vs ECOSAR | fish (n=2,324) | **2.77 (+0.03, -0.03)** | 4.08 (+0.08, -0.08) |
| vs ECOSAR | crustaceans (n=2,346) | **2.96 (+0.04, -0.04)** | 5.15 (+0.11, -0.11) |
| vs ECOSAR | algae (n=444) | **4.14 (+0.18, -0.17)** | 7.42 (+0.41, -0.39) |
| vs ECOSAR | worms (n=9) | **3.22 (+1.15, -0.85)** | 81.99 (+40.42, -27.08) |
| vs VEGA | fish (n=1,592) | **2.47 (+0.03, -0.03)** | 3.96 (+0.09, -0.08) |
| vs VEGA | crustaceans (n=1,372) | **2.54 (+0.04, -0.04)** | 4.32 (+0.1, -0.1) |
| vs VEGA | algae (n=439) | **2.81 (+0.09, -0.09)** | 5.34 (+0.27, -0.26) |
| vs VEGA | rodents (n=20,908) | **1.74 (+0.0, -0.0)** | 4.35 (+0.03, -0.03) |
| vs TEST | fish (n=2,435) | **2.66 (+0.03, -0.03)** | 3.19 (+0.05, -0.05) |
| vs TEST | crustaceans (n=2,204) | **2.95 (+0.04, -0.04)** | 3.94 (+0.07, -0.07) |
| vs TEST | rodents (n=26,157) | **1.79 (+0.0, -0.0)** | 3.75 (+0.02, -0.02) |
| vs Posthuma | fish (n=1,867) | **2.3 (+0.02, -0.02)** | 2.9 (+0.04, -0.04) |
| vs Posthuma | crustaceans (n=1,411) | **2.7 (+0.04, -0.04)** | 4.59 (+0.12, -0.11) |
| vs Posthuma | algae (n=171) | **3.1 (+0.18, -0.17)** | 5.13 (+0.33, -0.31) |

**Table S3. Pairwise Median Absolute Error fold change (MAE) comparison for NOEC/EC10 prediction.** Prediction were obtained from each compared model (see Methods, Supplementary Text 2, and Supplementary Table 13)) to estimate NOEC/EC10 MAE for each species/groups yielding predictions for each model. Errors were calculated for all predictable chemicals (same as Fig. 4d) and for chemicals falling inside each tools' respective applicability domain (only ECOSAR and VEGA differ). Bold values indicate lowest error in the comparison.

| Comparison NOEC\|EC10 | Eukaryotic group (n chemicals) | MAE TRIDENT-2 | MAE other model |
|---|---|---|---|
| **Predictable chemicals** | | | |
| vs TRIDENT-1 | fish (n=1,601) | **3.99 (+0.08, -0.08)** | 4.94 (+0.11, -0.11) |
| vs TRIDENT-1 | crustaceans (n=1,932) | **4.14 (+0.08, -0.08)** | 4.17 (+0.08, -0.08) |
| vs TRIDENT-1 | algae (n=629) | **5.02 (+0.2, -0.19)** | 5.24 (+0.21, -0.2) |
| vs ECOSAR | fish (n=1,844) | **4.32 (+0.09, -0.09)** | 17.67 (+0.64, -0.62) |
| vs ECOSAR | crustaceans (n=1,968) | **4.12 (+0.08, -0.08)** | 11.63 (+0.36, -0.35) |
| vs ECOSAR | algae (n=2,740) | **4.29 (+0.07, -0.07)** | 7.48 (+0.17, -0.17) |
| vs VEGA | fish (n=1,816) | **4.35 (+0.09, -0.09)** | 18.23 (+0.64, -0.62) |
| vs VEGA | crustaceans (n=1,811) | **4.55 (+0.09, -0.09)** | 9.26 (+0.28, -0.27) |
| vs VEGA | algae (n=2,452) | **4.33 (+0.08, -0.08)** | 7.95 (+0.18, -0.17) |
| vs VEGA | rodents (n=491) | **2.7 (+0.08, -0.08)** | 14.36 (+1.11, -1.03) |
| vs TEST | fish (n=0) | — | — |
| vs TEST | crustaceans (n=0) | — | — |
| vs TEST | rodents (n=0) | — | — |
| vs Posthuma | fish (n=0) | — | — |
| vs Posthuma | crustaceans (n=0) | — | — |
| vs Posthuma | algae (n=0) | — | — |
| **Predictable chemicals inside applicability domain** | | | |
| vs TRIDENT-1 | fish (n=1,601) | **3.99 (+0.08, -0.08)** | 4.94 (+0.11, -0.11) |
| vs TRIDENT-1 | crustaceans (n=1,932) | **4.14 (+0.08, -0.08)** | 4.17 (+0.08, -0.08) |
| vs TRIDENT-1 | algae (n=629) | **5.02 (+0.2, -0.19)** | 5.24 (+0.21, -0.2) |
| vs ECOSAR | fish (n=1,844) | **4.32 (+0.09, -0.09)** | 17.67 (+0.64, -0.62) |
| vs ECOSAR | crustaceans (n=1,968) | **4.12 (+0.08, -0.08)** | 11.63 (+0.36, -0.35) |
| vs ECOSAR | algae (n=2,740) | **4.29 (+0.07, -0.07)** | 7.48 (+0.17, -0.17) |
| vs VEGA | fish (n=103) | **3.2 (+0.2, -0.19)** | 4.65 (+0.45, -0.41) |
| vs VEGA | crustaceans (n=518) | **3.18 (+0.1, -0.09)** | 5.86 (+0.26, -0.24) |
| vs VEGA | algae (n=508) | **3.17 (+0.1, -0.09)** | 5.46 (+0.23, -0.22) |
| vs VEGA | rodents (n=39) | **2.7 (+0.36, -0.32)** | 20.05 (+3.86, -3.23) |
| vs TEST | fish (n=0) | — | — |
| vs TEST | crustaceans (n=0) | — | — |
| vs TEST | rodents (n=0) | — | — |
| vs Posthuma | fish (n=0) | — | — |
| vs Posthuma | crustaceans (n=0) | — | — |
| vs Posthuma | algae (n=0) | — | — |

**Table S4. Pairwise Median Absolute Error fold change (MAE) comparison for EC50 prediction in the QSAR applicability domain intersect.** Predictions were obtained from each compared model (see Methods, Supplementary Text 2, and Supplementary Table 13) to estimate EC50 MAE for each species/groups. Chemicals where either ECOSAR or VEGA flagged predictions as outside of their applicability domain for fish, crustaceans and algae were removed and only chemicals inside the tools' joint applicability domain were included for benchmarking. Values are the same as Supplementary Fig. S15. Bold values indicate lowest error in the comparison.

| Comparison EC50 | Eukaryotic group (n chemicals) | MAE TRIDENT-2 | MAE other model |
|---|---|---|---|
| vs TRIDENT-1 | fish (n=879) | **2.28 (+0.03, -0.03)** | 2.48 (+0.04, -0.04) |
| vs TRIDENT-1 | crustaceans (n=868) | **2.44 (+0.04, -0.04)** | 2.62 (+0.05, -0.05) |
| vs TRIDENT-1 | algae (n=37) | **3.91 (+0.48, -0.43)** | 4.17 (+0.56, -0.49) |
| vs ECOSAR | fish (n=1,189) | **2.36 (+0.03, -0.03)** | 2.99 (+0.06, -0.06) |
| vs ECOSAR | crustaceans (n=1,104) | **2.4 (+0.04, -0.04)** | 3.43 (+0.08, -0.08) |
| vs ECOSAR | algae (n=56) | **2.67 (+0.26, -0.24)** | 3.89 (+0.44, -0.39) |
| vs ECOSAR | worms (n=0) | — | — |
| vs VEGA | fish (n=1,189) | **2.36 (+0.03, -0.03)** | 3.59 (+0.08, -0.08) |
| vs VEGA | crustaceans (n=1,103) | **2.39 (+0.04, -0.04)** | 4.05 (+0.11, -0.1) |
| vs VEGA | algae (n=57) | **3.18 (+0.34, -0.3)** | 5.79 (+0.82, -0.72) |
| vs VEGA | rodents (n=0) | — | — |
| vs TEST | fish (n=1,156) | **2.33 (+0.03, -0.03)** | 2.64 (+0.04, -0.04) |
| vs TEST | crustaceans (n=1,047) | **2.41 (+0.04, -0.04)** | 3.11 (+0.07, -0.06) |
| vs TEST | rodents (n=0) | — | — |
| vs Posthuma | fish (n=97) | **2.23 (+0.11, -0.1)** | 3.11 (+0.26, -0.24) |

**Table S5. Pairwise Median Absolute Error fold change (MAE) comparison for NOEC/EC10 prediction in the QSAR applicability domain intersect.** Prediction were obtained from each compared model (see Methods, Supplementary Text 2, and Supplementary Table 13) to estimate NOEC/EC10 MAE for each species/groups applicable for each model. Chemicals where either ECOSAR or VEGA flagged predictions as outside of their applicability domain for fish, crustaceans and algae were removed and only chemicals inside the tools' joint applicability domain were included for benchmarking. Values are the same as Supplementary Fig. S16. Bold values indicate lowest error in the comparison. ECOSAR ChV predictions were converted into NOEC by division with the square root of 2.

| Comparison NOEC\|EC10 | Eukaryotic group (n chemicals) | MAE TRIDENT-2 | MAE other model |
|---|---|---|---|
| vs TRIDENT-1 | fish (n=66) | **3.75 (+0.34, -0.32)** | 3.88 (+0.33, -0.3) |
| vs TRIDENT-1 | crustaceans (n=363) | 3.14 (+0.11, -0.11) | **2.96 (+0.1, -0.1)** |
| vs TRIDENT-1 | algae (n=78) | **3.39 (+0.27, -0.25)** | 3.75 (+0.39, -0.35) |
| vs ECOSAR | fish (n=94) | **3.78 (+0.31, -0.29)** | 8.01 (+0.99, -0.88) |
| vs ECOSAR | crustaceans (n=492) | **3.17 (+0.1, -0.1)** | 9.07 (+0.5, -0.47) |
| vs ECOSAR | algae (n=503) | **3.17 (+0.1, -0.09)** | 5.13 (+0.21, -0.2) |
| vs ECOSAR | worms (n=0) | — | — |
| vs VEGA | fish (n=94) | **3.78 (+0.31, -0.29)** | 5.48 (+0.6, -0.54) |
| vs VEGA | crustaceans (n=492) | **3.17 (+0.1, -0.1)** | 5.86 (+0.26, -0.25) |
| vs VEGA | algae (n=503) | **3.17 (+0.1, -0.09)** | 5.48 (+0.23, -0.22) |
| vs VEGA | rodents (n=0) | — | — |
| vs TEST | fish (n=0) | — | — |
| vs TEST | crustaceans (n=0) | — | — |
| vs TEST | rodents (n=0) | — | — |
| vs Posthuma | fish (n=0) | — | — |
| vs Posthuma | crustaceans (n=0) | — | — |

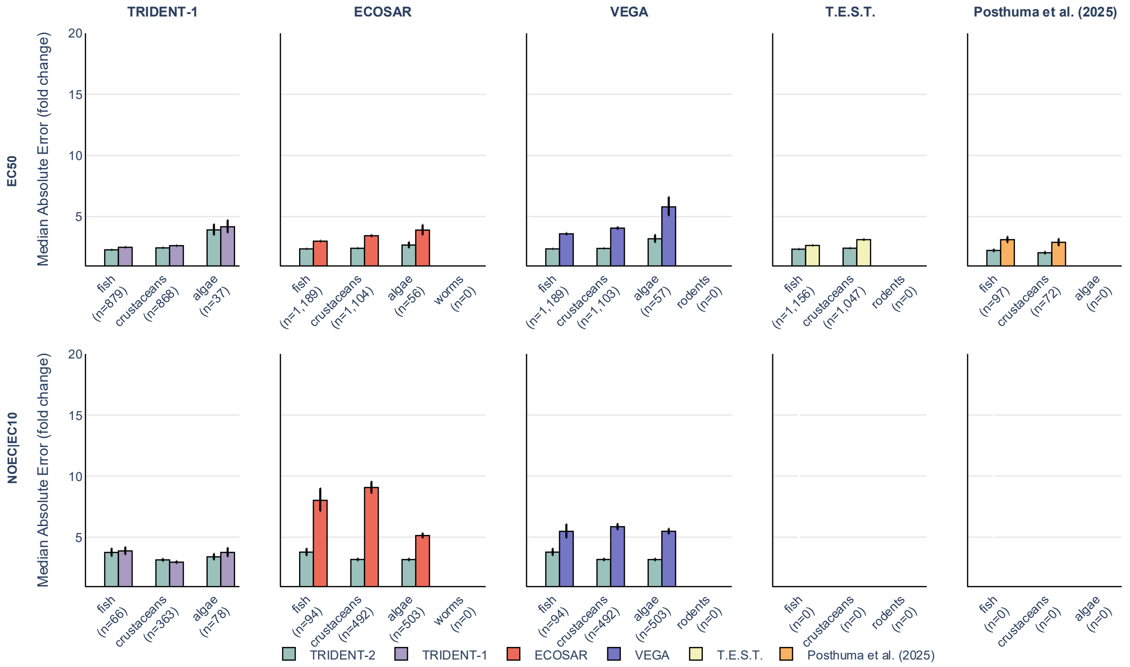

**Figure S15. Pairwise Median Absolute Error fold change (MAE) comparison for EC50 and NOEC/EC10 predictions inside each QSAR's respective applicability domain.** Prediction were obtained from each compared model (see Methods, Supplementary Text 2, and Supplementary Table 13) to estimate NOEC/EC10 MAE for each species/groups yielding predictions for each model. Predictions where either tool flagged predictions as outside of their applicability domain were removed (chenges only apply to ECOSAR and VEGA as all other tools do not yield such a distinction). ChV predictions were converted into NOEC by division with the square root of 2.

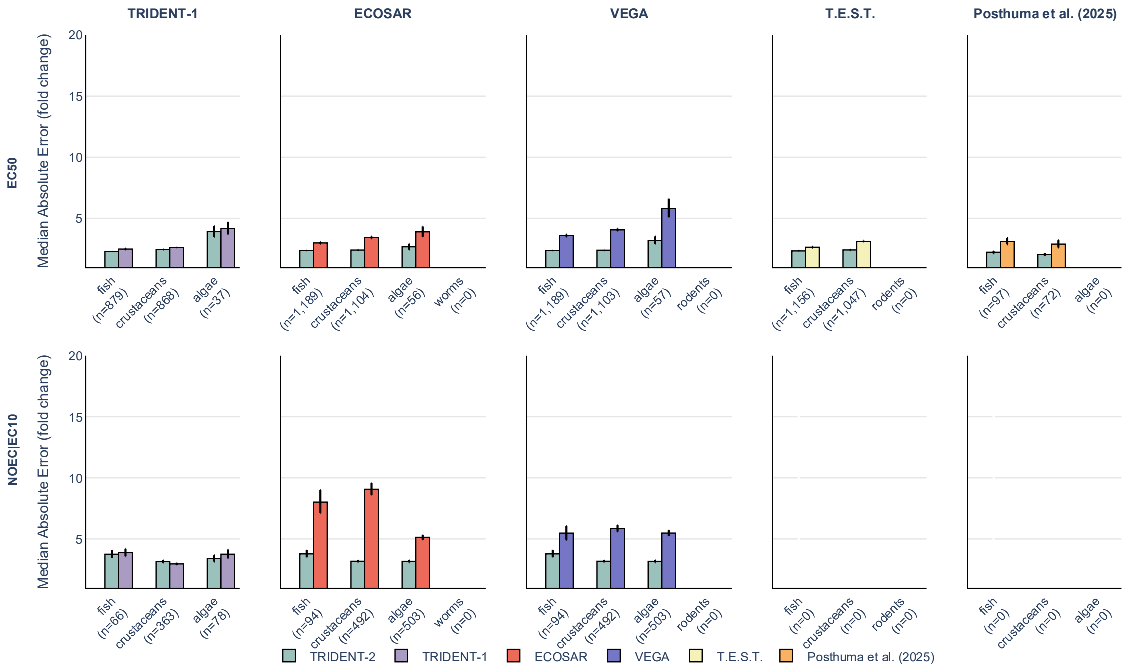

**Figure S16. Pairwise Median Absolute Error fold change (MAE) comparison for EC50 and NOEC/EC10 predictions in the QSAR applicability domain intersection.** Prediction were obtained from each compared model (see Methods, Supplementary Text 2, and Supplementary Table 13) to estimate NOEC/EC10 MAE for each species/groups applicable for each model. Chemicals where either ECOSAR or VEGA flagged predictions as outside of their applicability domain for fish, crustaceans and algae were removed and only chemicals inside the tools' joint applicability domain were included for benchmarking. ChV predictions were converted into NOEC by division with the square root of 2.

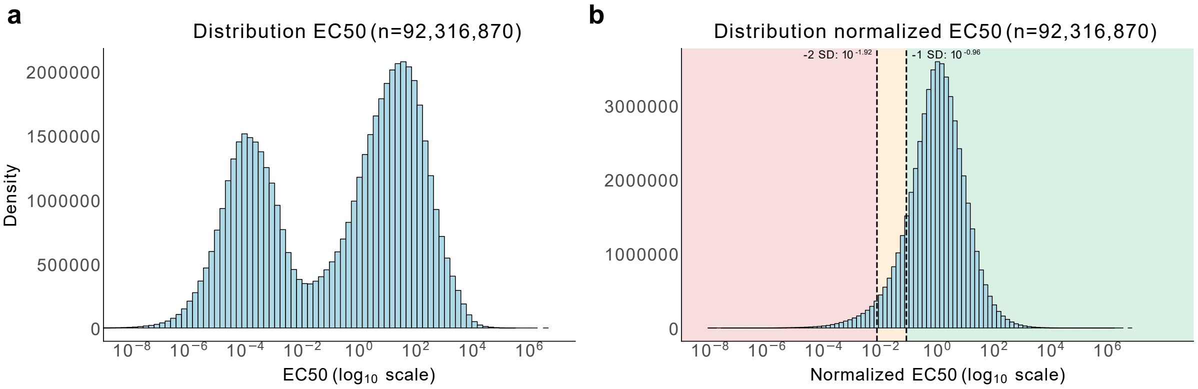

**Figure S17. Predicted EC50 distribution across all species and 13,590 chemicals in the dataset before and after normalization. a)** Predicted EC50 distribution (n=92,316,870) across all species (n=6,793) and 12,890+2,144 chemicals (chemicals in the dataset for which there were not solely rodent data available + chemicals in combined PPDB and BPDB, yielding in total 13,590 unique chemicals after canonicalization and deduplication). **b)** Predicted EC50 distribution after nomalizing each prediction by its species' median EC50 predictio. Associated toxicity intervals are indicated by vertical lines and shaded areas at -1 standard deviation (SD) and -2 SD (distribution mean -0.028 and standard deviation 0.956).

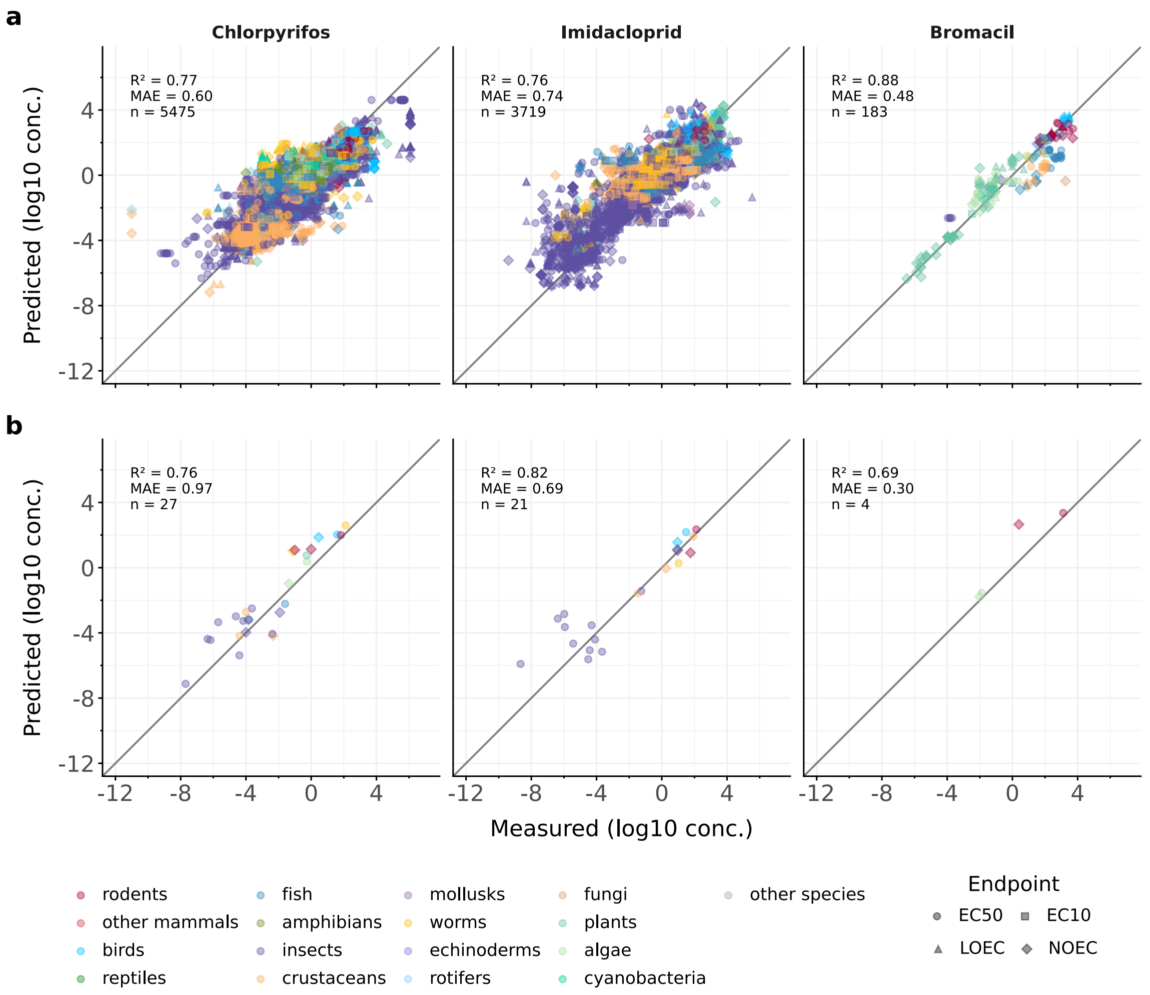

**Figure S18. Predicted toxicity versus measured values for chlorpyrifos, imidacloprid, and bromacil.** Predicted versus measured toxicity (EC50, EC10, LOEC, and NOEC) for the three chemicals with associated coefficient of determination (R2), Median Absolute Error (MAE) and number of samples (n). Predictions show the mean of the nine cross-validation models that had not seen the chemical during training from the 0.2 Butina distance dataset split. Predictions thus reflect the a model that has not seen each chemical nor its close analogues. **a)** Predicted and measured toxicity for records in the dataset. **b)** Same as a) but showing predicted versus measured toxicity for records in PPDB and BPDB.

### 2 Supplementary Methods

**Table S6. Eukaryotic group assignment of species based on taxonomic groups.** Species were semi manually assigned to eukaryotic groups, partly based on harmonized U.S. EPA ECOTOXKnowledgebase species groups. Species not falling into a category were grouped as "other species". The table contains NCBI taxids used to assign species at specified taxonomic ranks.

| | | **Taxonomic rank** | | | | |
|---|---|---|---|---|---|---|
| **Group** | **Comment** | **order** | **class** | **subphylum** | **phylum** | **kingdom** |
| **rodents** | | 9989 | 40674 | | | |
| **other mammals** | All mammals except rodents | | 40674 | | | |
| **birds** | | | 8782 | | | |
| **reptiles** | | | 8504 | | | |
| | | 8459 \| 1294634 | | | | |
| **amphibians** | | | 8292 | | | |
| **fish** | | | 186623 \| 7777 \| 2682552 \| 117569 | | | |
| **insects** | insects, spiders, and other arthropods | | | | 6656 | |
| **crustaceans** | crustacea | | | 6657 | 6656 | |
| | horseshoe crabs | | 6844 | | 6656 | |
| **mollusks** | Mollusca and other shells and snails | | | | 6447 \| 7568 | |
| **worms** | | | | | 33310 \| 6157 \| 43120 \| 10229 \| 6340 \| 6231 \| 6217 \| 6178 | |
| **echinoderms** | | | | | 7586 | |
| **rotifers** | | | | | 10190 | |
| **fungi** | | | | | | 4751 |
| **plants** | | | | | 35493 | |
| **algae** | | | | | 2836 \| 3041 \| 2830 \| 2763 | |

| | | | | | |
|---|---|---|---|---|---|
| | | | 2870 \| 2864 \| 3035 \| 3027 \| 2825 \| 5747 \| 38410 \| 33859 \| 35675 \| 304573 | | |
| **protozoans** | | 2605435 \| 136419 \| 5878 \| 6020 \| 5977 \| 6015 \| 37471 \| 5988 \| 33827 \| 194287 \| 422676 \| 1280412 \| 33829 \| 5653 \| 2779609 \| 6000 \| 1485085 \| 2681632 \| 555280 \| 2497438 | | | |
| **cyanobacteria** | | | | | 1117 |

**Table S7. Stepwise endpoint harmonization strategy.** Endpoints were assigned according to a tiered framwork based on exact string matches, wildcard matches, and series of regex matches.

| Tier | Matching Strategy | Endpoint Pattern | Target |
|---|---|---|---|
| 1 | Exact String Match (case-insensitive) | EC50, EC10, NOEC, LOEC | |
| 1a | Direct match to EC50 family | EC50, LC50, IC50, ID50, LD50, ED50 | EC50 |
| 1b | Direct match to EC10 family | EC10, LC10, IC10, EC15, EC05, EC16, EC17, EC12 | EC10 |
| 1c | Direct match to NOEC family | NOAEL, NOAEC, NOEL, NOEC | NOEC |
| 1d | Direct match to LOEC family | LOAEL, LOAEC, LOEC, LOEL | LOEC |
| 2 | Regex Pattern Matching: Two-Digit Suffixes | ^(EC\|LC\|IC\|ID\|LD\|ED)(\d{2}) | |
| 2a | First digit = 1 | EC1X, LC1X, LD1X, etc. | EC10 |
| 2b | First digit = 5 | EC5X, LC5X, LD5X, etc. | EC50 |
| 2c | First digit = 0, second digit ≥5 | EC0X (X≥5) | EC10 |
| 3 | Regex Pattern Matching: Single-Digit Suffixes | ^(EC\|LC\|IC\|ID\|LD\|ED)(\d{1}) | |
| 3a | Single digit ≥5 | ECX, LCX, LDX (X≥5) | EC10 |

**Table S8. Stepwise toxicological effect harmonization strategy.** Effects were assigned according to a tiered framework based on exact string matches, prefix wildcard matches and regex matches. Lethality endpoints were used to assign missing mortality effects.

| Tier | Matching Strategy | Effect Pattern | Target | Examples |
|---|---|---|---|---|
| 1 | Prefix Matching | | | |
| 1a | Starts with | MOR, LETH | MOR | mortality → MOR |
| 1b | Starts with | CAR, TUM, MUT | CAR | carcinogenic → CAR |
| 1c | Starts with | BEH, FBH, AVO | BEH | behavioral → BEH |
| 1d | Starts with | REP | REP | reprotoxic → REP |
| 2 | Substring Matching | | | |
| 2a | Contains | mor, leth, mort | MOR | effect mortality → MOR |
| 2b | Contains | tumor, mutation data | CAR | tumor formation → CAR |
| 2c | Contains | beh, fbh, behaviour, avo | BEH | avoidance response → BEH |
| 2d | Contains | rep | REP | reprotoxicity experiment → REP |
| 3 | Endpoint-Based Effect Inference | | | |
| 3a | Lethal endpoints with missing effect | lc, ld + NA effect | MOR | LC50 with no effect → MOR |

**Table S9. General summary of eukaryotic group organism life stage categorization.** Life stages were harmonized based on ECOTOX definitions. Each eukaryotic group could be assigned to a maximum of fours life stages; however, not all groups could be assigned to all three. The table describes which life stages were applicable for categorization for each group.

| Species group | # life stages | Life stages | Motivation |
|---|---|---|---|
| Plants | 1 | early | plants have two stages, seed and blank (other and plant) |
| Insects | 3 | early/mid/adult | early (larvae/egg), mid (pupa), adult (adult) |
| Fish | 3 | early/mid/adult | early (egg or yolk sack carry), mid (jv/fingerling etc, small fish), adult (adult) |
| Algae | None | | algae have only one stage so their are not assigned a life stage |
| Amphibians | 3 | early/mid/adult | early (egg), mid (tadpole), adult (adult) |
| Mollusks | 3 | early/mid/adult | early (egg), mid (jv), adult (sexually mature) |
| Rodents and Other mammals | 3 | early/mid/adult | early (before birth), mid (newborn-premature), adult (sexually mature) |
| Fungi | 3 | early | early (spore=early) |
| Birds | 2 | early/adult | early (egg), adult (adult) |

**Table S10. Detailed assignment of ECOTOX organism life stages into categories.** Life stages were harmonized based on ECOTOX definitions. In the case of ambiguous categories or missing information, we assigned NA (blanks). The table details assignment per ECOTOX life stage.

| ECOTOX life stage | ECOTOX description | Assigned category |
|---|---|---|
| -- | Unspecified | |
| AD | Adult | adult |
| AL | Alevin | early |
| BD | Bud or Budding | |
| BL | Blastula | early |
| BS | Bud blast stage | |
| BT | Boot | |
| CC | Cocoon | mid |
| CM | Corm | |
| CO | Copepodid | early |
| CP | Copepodite | mid |
| CS | Cleavage stage | early |
| CY | Cyst | |
| EG | Egg | early |
| EL | Elver | mid |
| EM | Embryo | early |
| EX | Exponential growth phase | |
| EY | Eyed egg or stage | early |
| F0 | F0 generation | |
| F1 | F1 generation | |
| F11 | F11 generation | |
| F2 | F2 generation | |
| F3 | F3 generation | |
| F6 | F6 generation | |
| F7 | F7 generation | |
| FB | Mature (full-bloom stage) organism | |
| FG | Female gametophyte | |
| FI | Fingerling | mid |
| FO | Flower opening | |
| FT | Froglet | mid |
| FY | Fry | mid |
| GA | Gastrula | early |
| GE | Gestation | early |
| GL | Glochidia | mid |
| GM | Gamete | early |
| GP | Lag growth phase | |
| GPS | Grain or seed formation stage | |
| GS | Germinated seed | |
| HD | Heading | |
| IB | Incipient bud | |

| | | |
|---|---|---|
| IE | Internode elongation | |
| IG | Imago | adult |
| IM | Immature | mid |
| IN | Instar | mid |
| IT | Intermolt | mid |
| JN | Jointing | |
| JV | Juvenile | mid |
| LC | Lactational | adult |
| LE | Egg laying | adult |
| LP | Larva-pupa | mid |
| LR | Prolarva | early |
| LV | Larva | early |
| MA | Mature | adult |
| MD | Mature dormant | adult |
| ME | Megalopa | mid |
| MG | Male gametophyte | |
| ML | Morula | early |
| MN | Mid-neurula | early |
| MO | Molt | mid |
| MX | Multiple | |
| MY | Mysis | early |
| NB | Newborn | mid |
| ND | Naiad | mid |
| NE | Neonate | mid |
| NH | New | mid |
| NL | Neurula | early |
| NOINT | Not intact | |
| NR | Not reported | |
| NU | Nauplii | early |
| NY | Nymph | mid |
| OO | Oocyte | early |
| PA | Parr | mid |
| PB | Mature | |
| PC | Pre-hatch | early |
| PD | Pre-molt | mid |
| PE | Post-emergence | |
| PG | Post-spawning | |
| PH | Mature | |
| PHT | Post-hatch | mid |
| PI | Post-molt | |
| PJ | Pre- | adult |
| PK | Post-smolt | adult |
| PL | Pullet | mid |
| PN | Post-nauplius | early |

| | | |
|---|---|---|
| PO | Pollen | |
| PP | Postpartum | adult |
| PPU | Prepupal | early |
| PQ | Pre-larva | early |
| PRB | Prebloom | |
| PS | Pre-smolt | mid |
| PT | Protolarvae | early |
| PU | Pupa | mid |
| PV | Post-larva | mid |
| PW | Pre-spawning | |
| PY | Post-embryo | |
| PZ | Protozoea | early |
| RC | Rooted cuttings | |
| RH | Rhizome | |
| RP | Mature reproductive | adult |
| RST | Rootstock | |
| SA | Subadult | mid |
| SB | Shoot | |
| SC | Yolk sac larvae | early |
| SCS | Senescence | adult |
| SD | Seed | early |
| SE | Scape elongation | |
| SF | Sac fry | early |
| SG | Mature | |
| SI | Sexually immature | mid |
| SL | Seedling | |
| SM | Sexually mature | adult |
| SMT | Smolt | mid |
| SN | Sapling | |
| SO | Sporeling | |
| SP | Sperm | early |
| SR | Spore | early |
| ST | Spat | mid |
| SU | Swim-up | mid |
| SW | Spawning | adult |
| SY | Stationary growth phase | |
| TA | Tadpole | mid |
| TC | Tissue culture callus | |
| TLS | Tiller stage | |
| TU | Tuber | |
| TZ | Trophozoite | |
| UY | Underyearling | mid |
| VE | Veliger | early |
| VG | Mature vegetative | |

| | | |
|---|---|---|
| VI | Virgin | adult |
| WN | Weanling | mid |
| YA | Young adult | adult |
| YE | Yearling | |
| YO | Young | mid |
| YY | Young of year | mid |
| ZO | Zoea | early |
| ZS | Zygospore | early |
| ZY | Zygote | early |

232

**Table S11. Detailed assignment of ECOTOX and RTECS administration route harmonization and categorization.** Administration routes were harmonized using ECOTOX definitions and by frequency. Due to the large number of possible administration routes, we harmonized and categorized them to the reduce risk of overfitting the models. The resulting categories were: static, oral, renewal, intraperitoneal, environmental, flow_through, spray, intravenous, food, direct_application, subcutaneous, gavage, inhalation, culture_media, granular, topical, intramuscular, soaking, dermal, drinking, in_vitro, injection, and parenteral. In the case of multiple reported routes, ambiguous assignment, or missing information, we assigned NA (blanks).

| ECOTOX life stage | ECOTOX description | Assigned category |
|---|---|---|
| -- | Unspecified | |
| S | | static |
| oral | | oral |
| R | | renewal |
| intraperitoneal | | intraperitoneal |
| EN | | environmental |
| F | | flow_through |
| SP | | spray |
| intravenous | | intravenous |
| FD | | food |
| DA | | direct_application |
| HS | | spray |
| subcutaneous | | subcutaneous |
| GV | | gavage |
| inhalation | | inhalation |
| CM | | culture_media |
| GS | | spray |
| FS | | spray |
| E | Applied in static waters. | static |
| GG | | granular |
| TP | | topical |
| FU | | inhalation |
| DT | | food |
| IP | | intraperitoneal |
| IM | | intramuscular |
| SO | | soaking |
| dermal | | dermal |
| OR | | oral |
| DR | | drinking |
| CH | | oral |
| DR | | oral |
| LC | | oral |
| OM | | oral |
| DM | | dermal |
| IV | | intravenous |
| P | Pulse | renewal |

| | | |
|---|---|---|
| PR | | environmental |
| SC | | subcutaneous |
| intramuscular | | intramuscular |
| IVT | | in_vitro |
| O | Flowing water system, such as streams | environmental |
| AS | | spray |
| IJ | | injection |
| GM | | culture_media |
| L | drain away from soil, ash, or similar material by the action of percolating liquid, especially rainwater | environmental |
| IC | Exposed via the air sac or air cell of an organism | injection |
| parenteral | | parenteral |
| AQ | Aqueous | static |
| MM | immersion | soaking |
| oral: gavage | | gavage |
| oral: feed | | food |
| inhalation: vapour | | inhalation |
| oral: drinking water | | drinking |
| administration onto the skin | | dermal |
| oral: unspecified | | oral |
| oral: diet | | food |
| inhalation: aerosol | | inhalation |
| inhalation: unspecified | | inhalation |
| inhalation: gas | | inhalation |
| chemical added to tank with water (dissolved in water) | | static |

**Table S12. Summary of harmonized toxicology metadata.** The table describes the harmonized endpoints, effects, concentration units, exposure duration units, organism life stages, and administration routes.

| **Category** | **Values** |
|---|---|
| Endpoints | NOEC (No Observed Effect Concentration),<br>LOEC (Lowest Observed Effect Concentration),<br>EC10 (10% Effect Concentration),<br>EC50 (50% Effect Concentration) |
| Effects | MOR (Mortality),<br>DVP (Developmental),<br>BEH (Behavioral),<br>REP (Reproductive),<br>CAR (Carcinogenicity),<br>GRO (Growth),<br>ITX (Intoxication),<br>PHY (Physiological),<br>MPH (Morphological),<br>POP (Population) |
| Concentration Units | mg/L,<br>mg/kg,<br>kg/m$^2$,<br>%,<br>ppm,<br>mg/kg bw,<br>mg/kg soil,<br>mg/org,<br>mg/L air,<br>mg/kg diet,<br>mg/kg org,<br>mL/kg bw,<br>mg/org/d,<br>mg/kg bw/d |
| Exposure duration units | h |
| Organism life stages | early, mid, adult, NA |
| Administration routes | static, oral, renewal, intraperitoneal, environmental, flow_through, spray, intravenous, food, direct_application, subcutaneous, gavage, inhalation, culture_media, granular, topical, intramuscular, soaking, dermal, drinking, in_vitro, injection, and parenteral, NA |

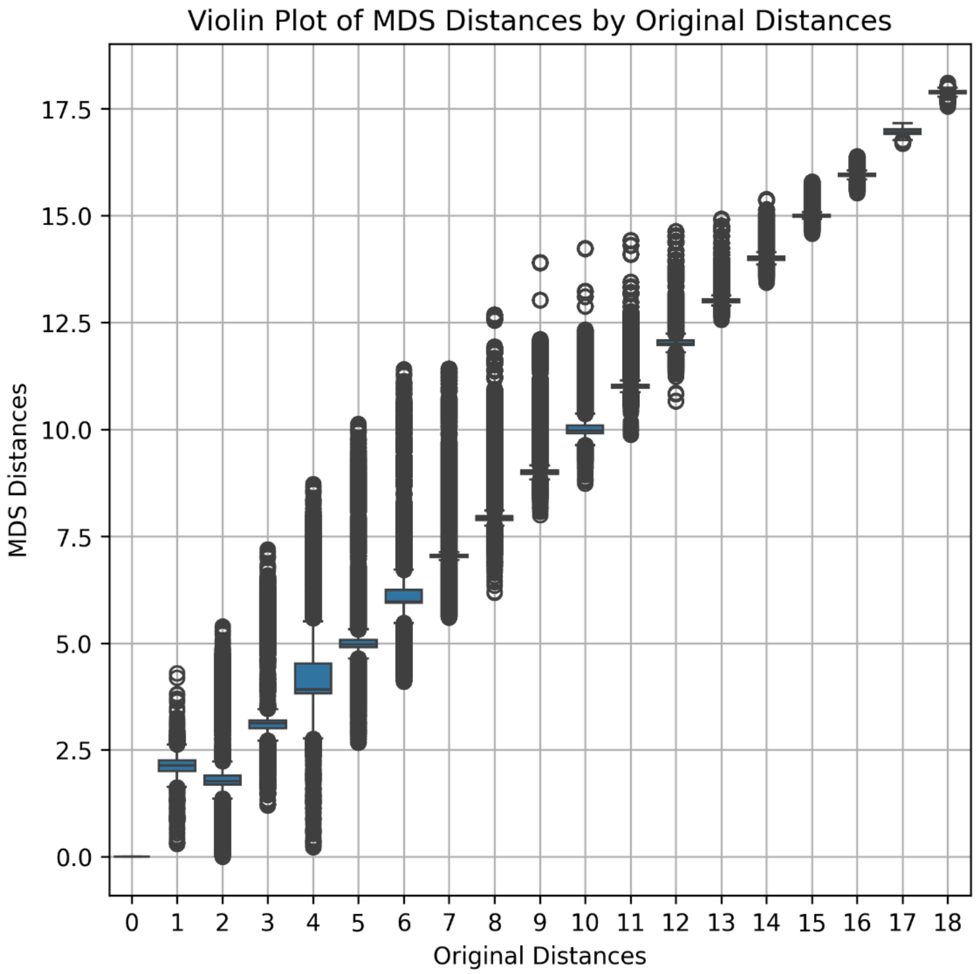

**Figure S19. Taxonomic embedding distances compared to taxonomic distances.** Metric Multidimensional Scaling (MDS) generated taxonomic embedding distances versus taxonomic distances for the taxonomic tree. MDS was used to obtain taxonomic embeddings for each taxa in the dataset as 768-dimensional vectors for each taxonomic ID in the data (including species, genus, family, order, class, subphylum, phylum, kingdom, superkingdom) preserving taxonomic distances. The MDS algorithm optimizes a stress value (MSE of recreated distance matrix) iteratively. Distances were calculated as the number of edges between two nodes and not distance to common ancestor. The maximum distance is 9+9=18, reflecting a walk from one species in Eukaryota to a species in (cyano)Bacteria. The minimum distance, 1, reflects a walk from one node to its parent node (e.g., species to genus or phylum to kingdom). A value of 2 reflects a walk between two sister nodes or walking to the grandparent node (e.g., *Danio rerio* to *Danio margaritatus* or *Danio rerio* to *Danionidae*). The Pearson and Spearman correlation between the staxonomic distance matrix and the recreated MDS vector distances were 0.999 and 0.990 respectively.

**Supplementary Text 1. Calculation of prediction errors.**

To assess model performance, we implemented a hierarchical aggregation strategy. First, raw predictions were aggregated to the level of unique experimental setups. This approach involves: (1) computing the median across technical replicates for each unique combination of endpoint, effect, chemical, concentration unit, taxonomic rank, exposure duration, administration route, and organism life stage; (2) calculating the absolute error for each experimental setup; (3) calculating the mean across different experimental effects for each endpoint-chemical-species combination. This results in one absolute error per endpoint-chemical-species combination.

Median Aggregation to reduce technical variability:

$$\tilde{y}_{i,c,s,e,d} = \text{median}\{y_{i,c,s,e,d,m}\}$$

where $i$ indexes endpoint, $e$ indexes effect, $c$ indexes chemical, $s$ indexes species, $d$ indexes exposure duration, and $m$ indexes replicate, administration route and life stage.

Absolute error for each experimental setup:

$$\epsilon_{i,c,s,e,d} = \left|\hat{y}_{i,c,s,e,d} - \tilde{y}_{i,c,s,e,d}\right|$$

Mean Error Aggregation (experimental setup → Endpoint-Effect-Chemical-Species):

$$\bar{\epsilon}_{i,c,s,e} = \frac{1}{\left|\mathcal{M}_{i,c,s,e}\right|} \sum_{m \in \mathcal{M}_{i,c,s,e}} \epsilon_{i,c,s,e,d}$$

For effect-specific performance estimation, aggregation stops here. This results in one error per unique endpoint, effect, chemical, and species combination.

Weighted Mean Error Aggregation (effects → Endpoint-Chemical-Species):

$$\bar{\epsilon}_{i,c,s} = \frac{\sum_{e \in \mathcal{E}_{i,c,s}} \omega_{i,c,s,e} \cdot \bar{\epsilon}_{i,c,s,e}}{\sum_{e \in \mathcal{E}_{i,c,s}} \omega_{i,c,s,e}}$$

where $\omega_{i,c,s,e}$ is the number of observations for each combination of $i$, $e$, $c$, $s$ in the data. This results in one error per unique endpoint, chemical, and species combination.

Subsequently, we aggregated the errors to endpoint-effect-eukaryotic group and endpoint-eukaryotic group combinations respectively by

(i) averaging, which first averages across both chemicals and species within each endpoint-species group, then computes the final metric. This was used for estimations where we did not want to estimate performance for novel chemicals or species.
Macro-Averaged Median Absolute Error (every observation has equal weight):

$$\text{MAE}_{\text{macro}}^{(i,g)} = \text{median}\left\{\frac{1}{|\mathcal{C}||\mathcal{S}_g|}\sum_{c \in \mathcal{C}}\sum_{s \in \mathcal{S}_g} \bar{\epsilon}_{i,c,s}\right\}$$

(ii) taxon-averaging, which averages across species before computing metrics. This was used to estimate the performance of novel species.
Taxon-Averaged Median Absolute Error (every taxon has equal importance):

$$\mathrm{MAE}_{\mathrm{taxon}}^{(i,g)} = \mathrm{median}\left\{\frac{1}{|\mathcal{C}|}\sum_{c\in\mathcal{C}}\frac{1}{|\mathcal{S}_g|}\sum_{s\in\mathcal{S}_g}\bar{\epsilon}_{i,c,s}\right\}$$

(iii) chemical-averaging, which averages across chemicals before computing metrics. This was used to estimate performance of novel chemicals.
Chemical-Averaged Median Absolute Error (every chemical has equal importance):

$$\mathrm{MAE}_{\mathrm{chem}}^{(i,g)} = \mathrm{median}\left\{\frac{1}{|\mathcal{S}_g|}\sum_{s\in\mathcal{S}_g}\frac{1}{|\mathcal{C}|}\sum_{c\in\mathcal{C}}\bar{\epsilon}_{i,c,s}\right\}$$

where $\mathcal{S}_g$ are all species in species group $g$ and $\mathcal{C}$ the set of all chemicals.

Finally, spread was measured as the Median Absolute Deviation Standard Error, define as:

$$\text{MAD-SE} = \frac{\mathrm{MAD}}{\sqrt{n}} = \frac{\mathrm{median}\{|x_i - \tilde{x}|\}}{\sqrt{\mathrm{n}}}$$

where $\tilde{x}$ is the median of the distribution and $n$ is the sample size.

**Supplementary Text 2. Obtaining model predictions from ECOSAR, VEGA, and T.E.S.T..**

83,970 SMILES (and CAS) were assigned internal identification numbers, and 58 SMILES were removed due to causing errors or soft locks (very large molecules, multiple bindings, multiple metals, high ring closure numbers) for at least one of the model software (see separate SMILES list in Supplementary Appendix, List of removed SMILES from QSAR comparison). The identifiers were separated into two sets: chemicals with no rat data (13,421), and the full dataset (83,970).

For ECOSAR 2.2, the no rat data subset was further divided into 14 files containing up to 1,000 SMILES each (to improve software performance) and subsequently fed into the model software. No additional CAS-based predictions were generated. Predictions for molecules with molecular weight above 1,000 and predictions flagged "LogKowCutOff" were removed. Additionally, ECOSAR 2.0 predictions were generated from the same set of SMILES, to capture "Should not be profiled"-tagged molecules, which were then removed from the ECOSAR 2.2 prediction set. This is necessary since ECOSAR 2.2 has removed this feature in batch mode. Finally, predicted ChV values were converted into NOEC by division with the square root of 2, following recommendations in ECHA (European Chemicals Agency, 2016).

For VEGA, the full 83 970 SMILES set was used, divided into 4 files containing up to 25 000 SMILES each (to account for java memory limits) and subsequently fed into the model software. For VEGA the following models were selected for predictions:

VEGA fish models:

- VEGA_Fish Acute (LC50) Toxicity model (KNN-Read-Across)
- VEGA_Fish Acute (LC50) Toxicity model (NIC)
- VEGA_Fish Acute (LC50) Toxicity model (IRFMN)
- VEGA_Fish Acute (LC50) Toxicity model (IRFMN-Combase)
- VEGA_Fish Chronic (NOEC) Toxicity model (IRFMN)
- VEGA_Fathead Minnow LC50 96h (EPA)
- VEGA_Fathead Minnow LC50 model (KNN-IRFMN)
- VEGA_Guppy LC50 model (KNN-IRFMN)

VEGA aquatic invertebrate models:

- VEGA_Daphnia Magna LC50 48h (EPA)
- VEGA_Daphnia Magna LC50 48h (DEMETRA)
- VEGA_Daphnia Magna Acute (EC50) Toxicity model (IRFMN)
- VEGA_Daphnia Magna Acute (EC50) Toxicity model (IRFMN-Combase)
- VEGA_Daphnia Magna Chronic (NOEC) Toxicity model (IRFMN)

VEGA algae models:

- VEGA_Algae Acute (EC50) Toxicity model (IRFMN)
- VEGA_Algae Acute (EC50) Toxicity model (ProtoQSAR-Combase)
- VEGA_Algae Chronic (NOEC) Toxicity model (IRFMN)

VEGA rat models (human toxicity):

- VEGA_Acute Toxicity (LD50) model (KNN)
- NOAEL (under the "other" category of human toxicity) (IRFMN)

Failed predictions (output "[Error]") were removed.

For T.E.S.T., the full SMILES set was used, separated into 28 files with 3,000 SMILES each. The consensus acute toxicity models for *Daphnia magna*, fish and rat were selected.

**Supplementary Text 3. Estimating predictability of eukaryotic taxonomic orders, eukaryotic species and chemicals for ECOSAR, VEGA, T.E.S.T., and TRIDENT-1.**

For each tool, predictability was quantified along three axes—eukaryotic taxonomic order, species, and chemicals—separately for each toxicity endpoint (EC50, NOEC/EC10) and, within each endpoint, separately for each possible species group (fish, crustaceans, algae, worms, rodents, plants, insects, protozoans). For every axis, coverage was expressed as the number of unique reference values (from the training dataset) that were predictable with the tool, divided by the total number of unique reference values.

For ECOSAR, VEGA, and T.E.S.T., chemical coverage was determined empirically: a SMILES was considered "covered" if the tool returned a prediction for it. Species- and order coverage were determined from each tool's documentation (QMRF/model documentation), i.e., a fixed curated list of species (or taxa) that each tool is validated to predict for a given species group (Supplementary Table S13). At the taxonomic-order level, this was applied generously: any NCBI order present in the reference dataset under an applicable species group was counted as "covered" by the tool, except for ECOSAR's algae and worm domains, which were additionally restricted to the phyla *Chlorophyta* (green algae) and *Annelida* (segmented worms), respectively, and ECOSAR's crustacean species domain, which was restricted to daphnids and mysids (though order-level coverage for crustaceans was not further restricted).

TRIDENT-1 predicts EC50 and NOEC/EC10 for three species groups without species separation—fish, crustaceans, and algae; all other species groups and endpoints were assigned zero coverage. Within the three species groups and two endpoints, TRIDENT-1 covers 100 % of the chemicals, and exposure durations present in the reference dataset for that group. For taxonomic orders and species, TRIDENT-1 metrics were treated similarly to ECOSAR.

**Table S13. Assigned predictable species for each tool.**

| Tool | Species group | Species/taxa considered within applicability domain |
|---|---|---|
| T.E.S.T. | protozoans | *Tetrahymena pyriformis* |
| T.E.S.T. | crustaceans | *Daphnia magna* |
| T.E.S.T. | fish | *Pimephales promelas* |
| T.E.S.T. | rodents | *Rattus norvegicus* |
| VEGA | fish | *Poecilia reticulata*, *Oncorhynchus mykiss*, *Pimephales promelas*, *Oryzias latipes* |
| VEGA | crustaceans | *Daphnia magna* |
| VEGA | algae | *Raphidocelis subcapitata*, *Selenastrum capricornutum*, *Pseudokirchneriella subcapitata* (synonymous taxa) |
| VEGA | rodents | *Rattus norvegicus* |
| ECOSAR | fish | All fish species present in the reference dataset |
| ECOSAR | crustaceans | Daphnids and mysids only |
| ECOSAR | algae | Green algae (phylum Chlorophyta) |
| ECOSAR | worms | Earthworms (phylum Annelida, generously assigned) |
| ECOSAR | plants | *Lemna gibba* |
| TRIDENT-1 | fish | All fish species present in the reference dataset |
| TRIDENT-1 | crustaceans | All crustacean species present in the reference dataset |
| TRIDENT-1 | algae | All algae species present in the reference dataset |

**Supplementary Text 4. Assigning inference settings.**

To be able to compare predictions from TRIDENT-2 across species, concentration units, effects, durations, administration routes, etc., we harmonized the inference settings so that each species was assigned its "standard input". Each species therefore has a specific combination of administration route, organism life stage, concentration unit, toxicological effect, and exposure duration. Instead of taking the most frequent value by species from the data, we aimed to harmonize the settings at the highest possible taxonomic level, since poorly studied species would otherwise risk deviating from their close relatives. We used a tiered assignment of these settings, starting at the eukaryotic group level and, where a group is not sufficiently uniform, splitting it by ECOTOX habitat (aquatic or terrestrial) or falling back on the most frequent value within each taxonomic family. For eukaryotic groups where standards, like the OECD guidelines, are typically used, we assigned these settings. See Supplementary Table S14-S17 for details. For organism life stage we assigned NA for all species.

Administration route and concentration unit are strongly coupled in the training data and were therefore assigned together. Aquatic test designs (static, renewal and flow-through) were reported in mg/L in 81-93% of records (the remainder mostly in ppm, approximately equal to mg/L), culture media in mg/L in 75%, spray applications in $kg/m^2$ in 81%, and oral, intraperitoneal and intravenous administration in mg/kg in 85-99%. The route "environmental" was not a consistent predictor of unit across groups (for example, $kg/m^2$ for plants but mg/kg soil for worms) and was therefore only assigned where the pairing is group-specific (terrestrial worms, Table S15). Missing route was assigned only to species with the “other species” species group.

Habitat (aquatic or terrestrial) was assigned per taxon, since the habitat recorded in the data is a property of the individual record and is frequently missing. Each species was assigned the majority habitat (water, soil or non-soil; ties resolved in that order) of its taxonomic family. If the family had no habitat data, the majority habitat of its order, then class, and finally its species group was used. Species with habitat "water" were treated as aquatic, all others as terrestrial. This assignment was manually corrected for 19 plant species whose ecology clearly contradicted the family majority: 12 terrestrial ornamentals and crops in Araceae (a family dominated by duckweed) were assigned soil, and 7 aquatic and emergent grasses in Poaceae (Glyceria, Sporobolus/Spartina, Echinochloa oryzicola and Oryza punctata) were assigned water. Remaining discrepancies between the assigned habitat and the habitat recorded for individual species (about 1% of species) mainly reflect the test medium or life stage (for example aquatic mosquito larvae tested as adults, or nematodes tested in liquid culture) rather than the ecology of the organism, and were left unchanged.

Acute effects were determined from EC50 records only, since chronic endpoints (NOEC, LOEC and EC10) are dominated by other effects, such as population growth of pest insects, that do not represent the acute standard. Species groups with a clear dominant acute effect in the EC50 data were assigned a group-level standard (Table S16). Immobilization (ITX), which is the standard acute endpoint for daphnids, was treated as equivalent to mortality (MOR) for crustaceans and other invertebrates. Plants were assigned growth (GRO), except floating plants (duckweeds—the genera Lemna, Spirodela, Landoltia and Wolffia—and Salviniaceae—Salvinia and Azolla), which were assigned population (POP). Habitat alone was not a suitable rule, since aquatic plants were split roughly evenly between GRO and POP. Species assigned to the “other species” group were assigned by family frequency.

**Table S14. Tiered assignment of standardized units, effects, and duration to taxonomic identifiers (TaxIDs) for toxicity prediction.** To keep consistent with standardized tests, we use species group assignment as a primary assignment. However, for many species groups data is ambiguous and requires other strategies.

| Input | Step | Decision Basis | Species Covered | Rule | Tie-Breaking |
|---|---|---|---|---|---|
| **Administration route** | 1 | Eukaryotic group and habitat | All eukaryotic groups except unmapped species | Route assigned together with the concentration unit according to predefined species-group standards (Table S15); plants, insects, mollusks and worms are split into aquatic and terrestrial by habitat | Not applicable |
| | 2 | Unmapped species group | Species with an unmapped species group | Assigned as missing route | Not applicable |
| **Concentration unit** | 1 | Eukaryotic group and habitat | All eukaryotic groups except unmapped species | Unit assigned together with the administration route according to predefined species-group standards (Table S15; e.g., aquatic taxa → mg/L; terrestrial vertebrates → mg/kg) | Not applicable |
| | 2 | Family frequency | Species not identified in Step 1 | The most frequent unit within each taxonomic family was assigned | In case of ties, the unit most consistent with the species-group standard was selected |
| **Effects** | 1 | Eukaryotic group | All eukaryotic groups except unmapped species | Acute effects were assigned using predefined species-group standards (Table S16: POP for algae, cyanobacteria, fungi and protozoans; GRO for plants; MOR for all other groups); floating plants (duckweeds and Salviniaceae) were assigned POP | Not applicable |
| | 2 | Family frequency | Species not identified in Step 1 (unmapped species group) | The most frequent acute effect (chosen from: MOR, ITX, POP, GRO) among EC50 records within each family was assigned | Ties were resolved by selecting the effect most consistent with the species-group assignment; if no family-level data were available, the species-group effect was used |
| **Duration** | 1 | Eukaryotic group | Algae, cyanobacteria, fish, crustaceans, amphibians, rodents, other mammals, reptiles, birds | Standardized acute durations (including NA) were assigned at the species-group level | Not applicable |
| | 2 | Family frequency with bounds | Species not identified in Step 1 | The most frequent duration within each family was assigned, constrained to a minimum of 1 hour | Ties or missing family data were resolved using the species-group duration |

| | | | | | |
|---|---|---|---|---|---|
| | | | | and a maximum of 504 hours | |
| **Habitat (used in Step 1 above)** | 1 | Manual correction | 19 plant species (Araceae, Poaceae) contradicting the family majority | Habitat set by hand according to the ecology of the species | Not applicable |
| | 2 | Family majority | All other species | The most frequent recorded habitat (water, soil, non-soil) within the family was assigned | Ties resolved in the order water, soil, non-soil |
| | 3 | Order, class, species-group majority | Species whose family has no habitat data | The most frequent recorded habitat within the order, then class, then species group was assigned | Same as above |

**Table S15. Assignment of administration route and concentration unit based on species group and habitat.** Route and unit are assigned together. Groups with identical aquatic and terrestrial settings were not split by habitat. Species groups not present in the table were assigned a missing route and treated separately for the unit (Table S14).

| Eukaryotic group | Aquatic route | Aquatic unit | Terrestrial route | Terrestrial unit |
|---|---|---|---|---|
| algae, cyanobacteria, fish, crustaceans, amphibians, echinoderms, cnidarians & bryozoans, protozoans, rotifers, tunicates & sponges | static | mg/l | static | mg/l |
| rodents, other mammals, reptiles, birds | oral | mg/kg | oral | mg/kg |
| fungi | culture media | mg/l | culture media | mg/l |
| plants | static | mg/l | spray | $kg/m^2$ |
| insects | static | mg/l | spray | $kg/m^2$ |
| mollusks | static | mg/l | spray | $kg/m^2$ |
| worms | static | mg/l | environmental | mg/kg soil |

**Table S16 Assignment of acute toxic effect based on species group.** Eukaryotic groups not present in the table were treated separately.

| Eukaryotic group | Acute effect |
|---|---|
| algae, cyanobacteria, fungi, protozoans | POP |
| plants (floating plants: duckweeds and Salviniaceae) | GRO (POP) |
| fish, crustaceans, amphibians, birds, reptiles, rodents, other mammals, insects, worms, mollusks, rotifers, cnidarians & bryozoans, echinoderms, tunicates & sponges | MOR |

**Table S17. Assignment of acute exposure duration based on species group.** Eukaryotic groups not present in the table were treated separately.

| **Eukaryotic group** | **Duration (hours)** |
|---|---|
| Algae | 96 |
| Cyanobacteria | 96 |
| Fish | 96 |
| Crustaceans | 48 |
| Amphibians | 96 |
| Rodents | NA |
| Other mammals | NA |
| Reptiles | NA |
| Birds | NA |

### 4 Supplementary appendix

#### List of removed SMILES from QSAR comparison

Removed molecules:

[Tb].[Tb].[Tb].[Tb].[Tb].[Tb].[Tb].[Tb].[Tb].[Tb].[Tb].[Tb].[Tb].[Tb].[Tb].[Tb].[Tb].[Tb].[Tb].[Tb].[Tb].[Tb].[Tb].[Tb].[Tb].[Tb].[Tb].[Tb].[Tb].[Tb].[Tb].[Tb].[Tb].[Tb].[Tb].[Tb].[Tb].[Tb].[Tb].[Tb].[Tb].[Tb].[Tb].[Tb].[Tb].[Tb].[Tb].[Tb].[Tb].[Tb].[Tb].[Tb].[Tb].[Tb].[Tb].[Tb].[Tb].[Tb].[Tb].[Tb].[Tb].[Tb].[Tb].[Tb].[Tb].[Tb].[Tb].[Tb].[Tb].[Tb].[Tb].[Tb].[Tb].[Tb].[Tb].[Tb].[Tb].[Tb].[Tb].[Tb].[Tb].[Tb].[Tb].[Tb].[Tb],

[B].[B].[B].[B].[B].[Mo].[Mo],

CCC(C)C1NC(=O)C(CCC(=O)O)NC(=O)C(CCC(N)=O)NC(=O)CNC(=O)C(CO)NC(=O)C2CCCN2C(=O)C2CSSCC(C(=O)NC(C)C(=O)NC(C)C(=O)NC(C(=O)NC3CSSCC(C(=O)NC4CSSCC(C(=O)NC(CC(N)=O)C(=O)NC(C(=O)NC(Cc5ccc(O)cc5)C(=O)N5CCCC5C(=O)O)C(C)O)NC(=O)C(CCCCN)NC(=O)C(CC(=O)O)NC(=O)C(C)NC(=O)C(CO)NC4=O)NC(=O)C(CCCCN)NC(=O)C(C(C)CC)NC(=O)C(CCC(=O)O)NC(=O)C(C(C)O)NC(=O)CNC(=O)C(C(C)O)NC(=O)C(CC(N)=O)NC(=O)C(C(C)C)NC(=O)C(CO)NC(=O)C4CCCN4C3=O)C(C)O)NC(=O)CNC(=O)C(Cc3ccccc3)NC(=O)C(CCC(=O)O)NC(=O)C(CC(C)C)NC(=O)C(C(C)C)NC(=O)C(CCCCN)NC(=O)CNC(=O)C(CCCNC(=N)N)NC(=O)C(CO)NC(=O)C(CO)NC(=O)C3CSSCC(NC(=O)C(Cc4c[nH]c5ccccc45)NC(=O)C(CO)NC(=O)C(CCCCN)NC(=O)C(C(C)C)NC(=O)C(Cc4ccc(O)cc4)NC(=O)C(CSSCC(NC(=O)C(CCC(=O)O)NC(=O)C(N)CCCNC(=N)N)C(=O)NC(Cc4ccc(O)cc4)C(=O)NC(CC(C)C)C(=O)NC(CC(N)=O)C(=O)N4CCCC4C(=O)NC(Cc4cnc[nH]4)C(=O)NC(CC(=O)O)C(=O)NC(C(C)O)C(=O)NC(CCC(N)=O)C(=O)NC(C(C)O)C(=O)N2)NC1=O)C(=O)NC(CC(N)=O)C(=O)NC(C)C(=O)NC(Cc1c[nH]c2ccccc12)C(=O)N3,

C=C1CCC(N2C(=O)c3ccc(OCC#Cc4cc(N5CCC(OC6CC(Oc7ccc(-c8ccc9c%10cnccc%10n(C)c9c8)cn7)C6)CC5)ccn4)cc3C2=O)C(=O)N1.C=C1CCC(N2C(=O)c3ccc(OCC#Cc4ccnc(N5CCC(OC6CC(Oc7ccc(-c8ccc9c%10cnccc%10n(C)c9c8)cn7)C6)CC5)n4)cc3C2=O)C(=O)N1.C=C1CCC(N2C(=O)c3ccc(OCC#Cc4nccc(N5CCC(OC6CC(Oc7ccc(-c8ccc9c%10cnccc%10n(C)c9c8)cn7)C6)CC5)n4)cc3C2=O)C(=O)N1.C=C1CCC(N2Cc3cc(OCC#Cc4cc(N5CCC(OC6CC(Oc7ccc(-c8ccc9c%10cnccc%10n(C)c9c8)cn7)C6)CC5)ccn4)ccc3C2=O)C(=O)N1.C=C1CCC(N2Cc3cc(OCC#Cc4cccc(N5CCC(OC6CC(Oc7ccc(-c8ccc9c%10cnccc%10n(C)c9c8)cn7)C6)CC5)c4)ccc3C2=O)C(=O)N1.C=C1CCC(N2Cc3cc(OCC#Cc4nccc(N5CCC(OC6CC(Oc7ccc(-c8ccc9c%10cnccc%10n(C)c9c8)cn7)C6)CC5)n4)ccc3C2=O)C(=O)N1.C=C1CCC(N2Cc3cc(OCCOCC#Cc4ccc(OC5CC(Oc6ncc(-c7ccc8c9cnccc9n(C)c8c7)cc6C)C5)cn4)ccc3C2=O)C(=O)N1,

CC=Cc1cccc(N(Cc2ccc(-c3ccc4c(c3)CC=N4)cc2)C(=O)C2CCCCC2)c1.CC=Cc1cccc(N(Cc2ccc(-c3ccc4nc(C)oc4c3)cc2)C(=O)C2CCCCC2)c1.COC(=O)C=Cc1cccc(N(Cc2ccc(-c3ccc4c(c3)OCCN4)cc2)C(=O)C2CCCCC2)c1.COC(=O)C=Cc1cccc(N(Cc2ccc(-c3ccc4c(c3)OCCO4)cc2)C(=O)C2CCCCC2)c1.COC(=O)C=Cc1cccc(N(Cc2ccc(-c3ccc4c(c3)S(=O)(=O)CCN4)cc2)C(=O)C2CCCCC2)c1.COC(=O)C=Cc1cccc(N(Cc2ccc(-c3ccc4c(c3)SCCN4)cc2)C(=O)C2CCCCC2)c1.COC(=O)C=Cc1cccc(N(Cc2ccc(-c3ccc4nccnc4c3)cc2)C(=O)C2CCCCC2)c1,

CCCN(CCC1CCC(CC(=O)C=Cc2ccc(OC)cc2)CC1)C1CCc2nc(N)sc2C1.CCCN(CCC1CCC(CC(=O)C=Cc2cccc(C)c2)CC1)C1CCc2nc(N)sc2C1.CCCN(CCC1CCC(CC(=O)C=Cc2cccc(OC)c2)CC1)C1CCc2nc(N)sc2C1.CCCN(CCC1CCC(CC(=O)C=Cc2ccccc2C)CC1)C1CCc2nc(N)sc2C1.CCCN(CCC1CCC(CC(=O)C

=Cc2ccccc2OC)CC1)C1CCc2nc(N)sc2C1.CCCN(CCC1CCC(CC(=O)C=Cc2ccccn2)CC1)C1CCc2nc(N)sc2C1.CCCN(CCC1CCC(CC(=O)C=Cc2cccnc2)CC1)C1CCc2nc(N)sc2C1,

O.O.O.O.O.O.O.O.O.O.O.O.O.O.O.O.O.O.O.O.O.O.O.O.O.O.O.[Al+3].[Al+3].[Al+3].[Al+3].[Al+3].[Al+3].[Al+3].[Al+3].[Al+3].[Al+3].[Al+3].[Al+3].[Na+].[Na+].[Na+].[Na+].[Na+].[Na+].[Na+].[Na+].[Na+].[Na+].[Na+].[Na+].[O-2].[O-2].[O-2].[O-2].[O-2].[O-2].[O-2].[O-2].[O-2].[O-2].[O-2].[O-2].[O-2].[O-2].[O-2].[O-2].[O-2].[O-2].[O-2].[O-2].[O-2].[O-2].[O-2].[O-2].[Si+4].[Si+4].[Si+4].[Si+4].[Si+4].[Si+4].[Si+4].[Si+4].[Si+4].[Si+4].[Si+4].[Si+4],

CCC(C)C(NC(=O)CN)C(=O)NC(C(=O)NC(CCC(=O)O)C(=O)NC(CCC(N)=O)C(=O)NC1CSSCC2NC(=O)C(C(C)CC)NC(=O)C(CO)NC(=O)C(C(C)O)NC(=O)C(CSSCC(NC(=O)C(CC(C)C)NC(=O)C(Cc3c[nH]cn3)NC(=O)C(CCC(N)=O)NC(=O)C(NC(=O)C(NC(=O)C(N)Cc3ccccc3)C(C)C)C(N)=O)C(=O)NCC(=O)NC(CO)C(=O)NC(Cc3c[nH]cn3)C(=O)NC(CC(C)C)C(=O)NC(C(C)C)C(=O)NC(CCC(=O)O)C(=O)NC(C)C(=O)NC(CC(C)C)C(=O)NC(Cc3ccc(O)cc3)C(=O)NC(CC(C)C)C(=O)NC(C(C)C)C(=O)NC(C(=O)NCC(=O)NC(CCC(=O)O)C(=O)NC(CCCNC(=N)N)C(=O)NCC(=O)NC(Cc3ccccc3)C(=O)NC(Cc3ccccc3)C(=O)NC(Cc3ccc(O)cc3)C(=O)NC(C(=O)N3CCCC3C(=O)NC(CCCCN)C(=O)NC(C(=O)O)C(C)O)C(C)O)CSSCC(C(=O)NC(CC(N)=O)C(=O)O)NC(=O)C(Cc3ccc(O)cc3)NC(=O)C(CC(N)=O)NC(=O)C(CCC(=O)O)NC(=O)C(CC(C)C)NC(=O)C(CCC(N)=O)NC(=O)C(Cc3ccc(O)cc3)NC(=O)C(CC(C)C)NC(=O)C(CO)NC2=O)NC1=O)C(C)C,

CSCCC(NC(=O)C(CO)NC(=O)C(Cc1ccc(O)cc1)NC(=O)C(N)CO)C(=O)NC(CCC(=O)O)C(=O)NC(Cc1cnc[nH]1)C(=O)NC(Cc1ccccc1)C(=O)NC(CCCNC(=N)N)C(=O)NC(Cc1c[nH]c2ccccc12)C(=O)NCC(=O)NC(CCCCN)C(=O)N1CCCC1C(=O)NC(C(=O)NCC(=O)NC(CCCCN)C(=O)NC(CCCCN)C(=O)NC(CCCNC(=N)N)C(=O)NC(CCCNC(=N)N)C(=O)N1CCCC1C(=O)NC(C(=O)NC(CCCCN)C(=O)NC(C(=O)NC(Cc1ccc(O)cc1)C(=O)N1CCCC1C(=O)NC(CC(N)=O)C(=O)NCC(=O)NC(C)C(=O)NC(CCC(=O)O)C(=O)NC(CC(=O)O)C(=O)NC(CCC(=O)O)C(=O)NC(CO)C(=O)NC(C)C(=O)NC(CCC(=O)O)C(=O)NC(C)C(=O)NC(Cc1ccccc1)C(=O)N1CCCC1C(=O)NC(CC(C)C)C(=O)NC(CCC(=O)O)C(=O)NC(Cc1ccccc1)C(=O)O)C(C)C)C(C)C)C(C)C,

CCC(C)C(NC(=O)CNC(=O)C(CO)NC(=O)C(CC(C)C)NC(=O)C(NC(=O)C(CC(N)=O)NC(=O)CNC(=O)C(CCCNC(=N)N)NC(=O)C(NC(=O)C(CO)NC(=O)C1CCCN1C(=O)CNC(=O)C(CC(=O)O)NC(=O)C(CO)NC(=O)C(CC(=O)O)NC(=O)C(CS)NC(=O)C(CC(C)C)NC(=O)C(CS)NC(=O)C1CCCN1C(=O)C(NC(=O)CN)C(C)C)C(C)C)C(C)O)C(=O)NC(C(=O)NC(Cc1c[nH]c2ccccc12)C(=O)NC(CC(C)C)C(=O)NC(C)C(=O)NCC(=O)NC(CS)C(=O)N1CCCC1C(=O)NC(CO)C(=O)NCC(=O)NC(Cc1c[nH]c2ccccc12)C(=O)NC(Cc1cnc[nH]1)C(=O)NC(CC(N)=O)C(=O)NC(CS)C(=O)NC(CCCCN)C(=O)NC(CCCCN)C(=O)NC(Cc1cnc[nH]1)C(=O)NCC(=O)N1CCCC1C(=O)NC(C(=O)NC(C(=O)NCC(=O)NC(Cc1c[nH]c2ccccc12)C(=O)NC(CS)C(=O)NC(CS)C(=O)NC(CCCCN)C(=O)NC(CCC(N)=O)C(=O)O)C(C)CC)C(C)O)C(C)CC,

COC(=O)C1NC(=O)C2NC(=O)C(NC(=O)C3NC(=O)C4NC(=O)C(Cc5ccc(cc5)Oc5cc3cc(c5OC3OC(CO)C(O)C(O)C3OC3OC(CO)C(O)C(O)C3O)Oc3ccc(cc3Cl)C2OC2CC(N)C(O)C(C)O2)NC(=O)C(N)c2ccc(O)c(c2)Oc2cc4cc(OC3OC(CO)C(O)C(O)C3O)c2C)c2ccc(O)c(c2)-c2c(OC3OC(CO)C(O)C(O)C3O)cc(O)cc21,

COC1C(N)CC(OC2c3ccc(c(Cl)c3)Oc3cc4cc(c3OC3OC(CO)C(O)C(O)C3OC3CC(N)C(O)C(C)O3)Oc3ccc(cc3)C(O)C(NC(=O)C(N)c3ccc(O)cc3)C(=O)NC(Cc3ccccc3)C(=O)NC4C(=O)NC3C(=O)NC2C(=O)NC(C(=O)O)c2cc(O)cc(OC4OC(CO)C(O)C(O)C4O)c2-c2cc3ccc2O)OC1C,

CC(CCC1(O)OC2CC3C4CCC5CC(OC6OC(CO)C(OC7OC(CO)C(OC8OC(CO)C(O)C(OC9OC(C)C(OC%10OC(C)C(O)C(O)C%10O)C(O)C9OC9OC(C)C(O)C(O)C9O)C8OC8OCC(O)C(O)C8O)C(O)C7O)C(O)C6O),

CC(CCC1(O)OC2CC3C4CCC5CC(OC6OC(CO)C(OC7OC(CO)C(OC8OC(CO)C(O)C(OC9OC(C)C(OC%10OC(C)C(O)C(O)C%10O)C(O)C9OC9OC(C)C(O)C(O)C9O)C8OC8OCC(O)C(O)C8O)C(O)C7O)C(O)C6O)CCC5(C)C4CC(=O)C3(C)C2C1C)COC1OC(CO)C(O)C(O)C1O,

N#C[S-].[Br-].[Sn+4].[c-]1ccccc1.[c-]1ccccc1.[c-]1ccccc1.c1ccc([As+](c2ccccc2)(c2ccccc2)c2ccccc2)cc1,

COC(=O)C1c2cc3c(c(O)c2C(OC2CC(OC)C(OC4CC(O)C(OC5CC(OC)C(OC6CC(OC)C(OC(=O)C=CC(=O)O)C(C)O6)C(C)O5)C(C)O4)C(C)O2)CC1(C)O)C(=O)c1c(O)cc2c(c1C3=O)OC1OC2(C)C(OC2CC(O)C(OC3CC(C)([N+](=O)[O-])C(OC4CC(OC)C(O)C(C)O4)C(C)O3)C(C)O2)C(N(C)C)C1O,

CNC(C(=O)NC1C(=O)NC(c2ccc(O)c(Cl)c2)C(=O)NC2C(=O)NC3C(=O)NC(Cc4cc(Cl)c(cc4OC4CC(N)C(O)C(C)O4)Oc4cc2cc(c4OC2OC(CO)C(O)C(O)C2OC2CC(N)C(O)C(C)O2)Oc2ccc(cc2)C1OC1OC(CO)C(O)C(O)C1O)C(=O)NC(C(=O)O)c1cc(O)cc(O)c1-c1cc3ccc1O)c1ccc(OC2OC(C)C(O)C(O)C2O)cc1,

CNC(C(=O)NC1C(=O)NC(c2ccc(O)c(Cl)c2)C(=O)NC2C(=O)NC3C(=O)NC(C(=O)NC(C(=O)O)c4cc(O)cc(O)c4-c4cc3ccc4O)C(OC3CC(N)C(O)C(C)O3)c3ccc(c(Cl)c3)Oc3cc2cc(c3OC2OC(CO)C(O)C(O)C2OC2CC(N)C(O)C(C)O2)Oc2ccc(cc2)C1OC1OC(CO)C(O)C(O)C1O)c1ccc(O)cc1,

CNC(C(=O)NC1C(=O)NC(c2ccc(O)c(Cl)c2)C(=O)NC2C(=O)NC3C(=O)NC(C(=O)NC(C(=O)O)c4cc(O)cc(O)c4-c4cc3ccc4O)C(OC3CC(N)C(O)C(C)O3)c3ccc(c(Cl)c3)Oc3cc2cc(c3OC2OC(CO)C(O)C(O)C2OC2CC(N)C(O)C(C)O2)Oc2ccc(cc2)C1O)c1ccc(OC2OC(C)C(O)C(O)C2OC)cc1,

CNC(C(=O)NC1C(=O)NC(c2ccc(O)c(Cl)c2)C(=O)NC2C(=O)NC3C(=O)NC(C(=O)NC(C(=O)O)c4cc(O)cc(O)c4-c4cc3ccc4O)C(OC3CC(N)C(O)C(C)O3)c3ccc(c(Cl)c3)Oc3cc2cc(c3OC2OC(CO)C(O)C(O)C2O)Oc2ccc(cc2Cl)C1OC1OC(CO)C(O)C(O)C1O)c1ccc(OC2OC(C)C(O)C(O)C2O)cc1,

CNC(C(=O)NC1C(=O)NC(c2ccc(O)c(Cl)c2)C(=O)NC2C(=O)NC3C(=O)NC(C(=O)NC(C(=O)O)c4cc(O)cc(O)c4-c4cc3ccc4O)C(OC3CC(N)C(O)C(C)O3)c3ccc(c(Cl)c3)Oc3cc2cc(c3OC2OC(CO)C(O)C(O)C2O)Oc2ccc(cc2Cl)C1OC1OC(CO)C(O)C(O)C1O)c1ccc(O)cc1,

O=C1c2ccccc2C(=O)c2c1ccc1c2[nH]c2ccc3c4ccc5c6c(ccc(c(=O)c3c21)c46)c1ccc2c(=O)c3ccccc3c3nn5c1c23,

CC1=C2N=C(C=C3N=C(C(C)=C4[N-]C(C(CC(N)=O)C4(C)CCC(=O)NCC(C)OP(=O)([O-])OC4C(CO)OC(n5cnc6cc(C)c(C)cc65)C4O)C4(C)N=C1C(CCC(N)=O)C4(C)CC(N)=O)C(CCC(N)=O)C3(C)C)C(CCC(N)=O)C2(C)CC(N)=O.[CH2-]C1OC(n2cnc3c(N)ncnc32)C(O)C1O.[Co+3],

CCC(C)C(NC(=O)CNC(=O)C(CC(N)=O)NC(=O)C(CCCCN)NC(=O)C(NC(=O)C(CO)NC(=O)C1CCCN1C(=O)C(CS)NC(=O)CNC(=O)C(CS)NC(=O)CNC(=O)C(CCCNC(=N)N)NC(=O)C(CCC(=O)O)NC(=O)C(NC(=O)C(CCCNC(=N)N)NC(=O)C(Cc1ccc(O)cc1)NC(=O)CNC(=O)C(CCCNC(=N)N)NC(=O)C(Cc1cnc[nH]1)NC(=O)C(CC(=O)O)NC(=O)C(CCCNC(=N)N)NC(=O)C(Cc1c[nH]c2ccccc12)NC(=O)C(CCCNC(=N)N)NC(=O)C(CCCCN)NC(=O)C(CCCCN)NC(=O)C(Cc1ccc(O)cc1)NC(=O)C(CS)NC(=O)C(CC(N)=O)NC(=O)C(NC(=O)C(CCC(=O)O)NC(=O)CNC(=O)CNC(=O)C(CO)NC(=O)C(CS)NC(=O)CNC(=O)C(NC(=O)C(NC(=O)C(NC(=O)C1CCCN1C(=O)C(NC(=O)C(CCC(N)=O)NC(=O)C(CO)NC(=O)C(CO)NC(=O)C(CCC(N)=O)NC(=O)C(CCC(N)=O)NC(=O)C(CC(N)=O)NC(=O)C(Cc1cnc[nH]1)NC(=O)C(CS)NC(=O)C(CCC(=O)O)NC(=O)C(N)CC(C)C)C(C)O)C(C)O)C(C)O)C(C)O)C(C)O)C(C)O)C(C)C)C(=O)NC(CCC(=O)O)C(=O)NC(C(=O)NC(CC(N)=O)C(=O)NC(CS)C(=O)NC(CS)C(=O)NC(C(=O)NC(C(

=O)NC(CC(=O)O)C(=O)NC(CCCNC(=N)N)C(=O)NC(CS)C(=O)NC(CC(N)=O)C(=O)NC(CC(N)=O)C(=O)O)C(C)O)C(C)O)C(C)CC,

CCC(C)C1NC(=O)CNC(=O)C2CCCN2C(=O)C(CCCCN)NC(=O)C(C(C)C)NC(=O)C(C(C)O)NC(=O)C2CCCN2C(=O)C(NC(=O)CNC(=O)C2CSSCC3NC(=O)C(C(C)O)NC(=O)C(CCCCN)NC(=O)C(C(C)O)NC(=O)C(C(C)O)NC(=O)C(CCC(N)=O)NC(=O)C4CCCN4C(=O)C(CCC(N)=O)NC(=O)C(CO)NC(=O)C(CO)NC(=O)C(Cc4cnc[nH]4)NC(=O)C(CCC(N)=O)NC(=O)C(CC(N)=O)NC(=O)C(Cc4ccccc4)NC(=O)C(NC(=O)C(NC(=O)C(N)CCCNC(=N)N)C(C)CC)CSSCC(NC(=O)C(CO)NC(=O)C(CCC(=O)O)NC(=O)C(CO)NC(=O)CNC(=O)C(CO)NC(=O)C4CCCN4C3=O)C(=O)NC(Cc3ccc(O)cc3)C(=O)NC(CC(N)=O)C(=O)NC(CCCCN)C(=O)NC(CCC(N)=O)C(=O)NC(Cc3c[nH]c4ccccc34)C(=O)NC(CO)C(=O)NC(CC(=O)O)C(=O)NC(Cc3ccccc3)C(=O)NC(CCCNC(=N)N)C(=O)NCC(=O)NC(C(C)O)C(=O)NC(C(C)CC)C(=O)NC(C(C)CC)C(=O)NC(CCC(=O)O)C(=O)NC(CCCNC(=N)N)C(=O)NCC(=O)N2)CSSCC(C(=O)NC2CSSCC(C(=O)NC(CC(N)=O)C(=O)NC(CC(N)=O)C(=O)O)NC(=O)C(C(C)C)NC(=O)C(CCC(=O)O)NC(=O)C(CO)NC(=O)C(CCC(=O)O)NC2=O)NC(=O)C(CO)NC(=O)C(CC(C)C)NC(=O)C(CCCCN)NC1=O,

CCC(C)C(NC(=O)CNC(=O)C1CCCN1C(=O)C(CCCCN)NC(=O)C(NC(=O)C(NC(=O)C1CCCN1C(=O)C(CS)NC(=O)CNC(=O)C(CS)NC(=O)CNC(=O)C(CCCNC(=N)N)NC(=O)C(CCC(=O)O)NC(=O)C(NC(=O)C(NC(=O)C(NC(=O)CNC(=O)C(CCCNC(=N)N)NC(=O)C(Cc1ccccc1)NC(=O)C(CC(=O)O)NC(=O)C(CO)NC(=O)C(Cc1c[nH]c2ccccc12)NC(=O)C(CCC(N)=O)NC(=O)C(CCCCN)NC(=O)C(Cc1cnc[nH]1)NC(=O)C(Cc1ccc(O)cc1)NC(=O)C(CS)NC(=O)C(CO)NC(=O)C(CO)NC(=O)C(CCC(=O)O)NC(=O)CNC(=O)C(CO)NC(=O)C1CCCN1C(=O)C(CS)NC(=O)C(NC(=O)C(CCCCN)NC(=O)C(NC(=O)C(NC(=O)C(CCC(N)=O)NC(=O)C1CCCN1C(=O)C(CCC(N)=O)NC(=O)C(CO)NC(=O)C(CO)NC(=O)C(Cc1cnc[nH]1)NC(=O)C(CCC(N)=O)NC(=O)C(CC(N)=O)NC(=O)C(Cc1ccccc1)NC(=O)C(CS)NC(=O)C(NC(=O)C(N)CCCNC(=N)N)C(C)CC)C(C)O)C(C)O)C(C)O)C(C)O)C(C)CC)C(C)CC)C(C)O)C(C)C)C(=O)NC(CCCCN)C(=O)NC(CC(C)C)C(=O)NC(CO)C(=O)NC(CS)C(=O)NC(CS)C(=O)NC(CCC(=O)O)C(=O)NC(CO)C(=O)NC(CCC(=O)O)C(=O)NC(C(=O)NC(CS)C(=O)NC(CC(N)=O)C(=O)NC(CC(N)=O)C(=O)O)C(C)C,

CC(CCC1(O)OC2CC3C4CC=C5CC(OC6OC(CO)C(OC7OC(CO)C(O)C(O)C7OC7OC(CO)C(OC8OC(C)C(OC9OC(C)C(O)C(O)C9O)C(O)C8O)C(OC8OCC(O)C(O)C8O)C7O)C(O)C6O)CCC5(C)C4CCC3(C)C2C1C)COC1OC(CO)C(O)C(O)C1O,

CCC(C)C1NC(=O)C(CC(N)=O)NC(=O)C(CCCNC(=N)N)NC(=O)C(C)NC(=O)C(C(C)O)NC(=O)C(C(C)O)NC(=O)C(CO)NC(=O)C2CCCN2C(=O)C2CSSCC(NC(=O)CNC(=O)C(CO)NC(=O)C(CC(C)C)NC(=O)C(CO)NC(=O)C(C)NC(=O)C3CSSCC(NC(=O)C(C(C)O)NC(=O)C(CC(N)=O)NC(=O)C(Cc4ccc(O)cc4)NC1=O)C(=O)NC(CCCNC(=N)N)C(=O)NC(CC(C)C)C(=O)NC(C(C)O)C(=O)NCC(=O)NC(C)C(=O)NC(CO)C(=O)NC(CCCNC(=N)N)C(=O)NC(CO)C(=O)NC(C(C)C)C(=O)N3)C(=O)NC(CCCCN)C(=O)NC(C(C)CC)C(=O)NC(C(C)CC)C(=O)NC(CO)C(=O)NCC(=O)NC(CO)C(=O)NC(C(C)O)C(=O)NC(C(=O)NC(CC(=O)O)C(=O)NC(CO)C(=O)NCC(=O)NC(Cc1c[nH]c3ccccc13)C(=O)NC(CC(N)=O)C(=O)NC(Cc1cnc[nH]1)C(=O)O)CSSCC(NC(=O)C(CO)NC(=O)C(N)CCCCN)C(=O)N2,

COCC1OC(OC2OCC3OC4(OCC(O)(C(C)OC)C5OCOC54)OC3C2OC)C(OC)C(O)C1OC1OC(C)C(OC)C(OC2OC(C)C3OC4(CC(O)C(OC5CC(OC6CC(C)([N+](=O)[O-])C(OC)C(C)O6)C(OC(=O)c6c(C)c(Cl)c(O)c(Cl)c6OC)C(C)O5)C(C)O4)OC3(C)C2O)C1O,

COCC1OC(OC2OCC3OC4(OCC(O)(C(C)OC)C5OCOC54)OC3C2OC)C(OC)C(O)C1OC1OC(C)C(OC)C(OC2CC3(C)OC4(CC(O)C(OC5CC(OC6CC(C)([N+](=O)[O-])C(OC)C(C)O6)C(OC(=O)c6c(C)c(Cl)c(O)c(Cl)c6OC)C(C)O5)C(C)O4)OC3C(C)O2)C1O,

CCC(C)C1NC(=O)C(C(C)O)NC(=O)CNC(=O)C(CCCNC(=N)N)NC(=O)C(Cc2cnc[nH]2)NC(=O)C(CC(=O)O)NC(=O)C(CO)NC(=O)C(Cc2c[nH]c3ccccc23)NC(=O)C(Cc2c[nH]c3ccccc23)NC(=O)C(CCCCN)NC(=O)C(CCCCN)NC(=O)C(Cc2ccc(O)cc2)NC(=O)C2CSSCC(NC(=O)C(CCC(=O)O)NC(=O)C(N)CC(C)C)C(=O)NC(Cc3cnc[nH]3)C(=O)NC(CC(N)=O)C(=O)NC(CCC(N)=O)C(=O)NC(CCC(N)=O)C(=

O)NC(CO)C(=O)NC(CO)C(=O)NC(CCC(N)=O)C(=O)NC(C)C(=O)N3CCCC3C(=O)NC(C(C)O)C(=O)NC(C(C)O)C(=O)NC(CCCCN)C(=O)NC(C(C)O)C(=O)NC(CSSCC(C(=O)NCC(=O)NC3CSSCC(C(=O)NC4CSSCC(C(=O)NC(CC(N)=O)C(=O)NC(CC(N)=O)C(=O)O)NC(=O)C(CCCNC(=N)N)NC(=O)C(CC(=O)O)NC(=O)C(C(C)O)NC(=O)C(CCCNC(=N)N)NC4=O)NC(=O)C(CC(N)=O)NC(=O)C(CC(C)C)NC(=O)C(CCCCN)NC(=O)C(C(C)C)NC(=O)CNC(=O)C4CCCN4C(=O)C(CCCCN)NC(=O)C(C(C)C)NC(=O)C(CCCCN)NC(=O)C4CCCN4C3=O)NC(=O)CNC(=O)C(CCCNC(=N)N)NC(=O)C(CCC(=O)O)NC(=O)C(C(C)CC)NC1=O)C(=O)NC(CO)C(=O)NCC(=O)NC(CCC(=O)O)C(=O)NC(C(C)O)C(=O)NC(CC(N)=O)C(=O)N2,

CCC(C)C(NC(=O)C(Cc1ccc(O)cc1)NC(=O)C(C)NC(=O)C(CC(=O)O)NC(=O)C(CCCNC(=N)N)NC(=O)CN)C(=O)NC(C)C(=O)NC(CCC(N)=O)C(=O)N1CCCC1C(=O)NC(CCC(=O)O)C(=O)NC(CC(N)=O)C(=O)NC(CS)C(=O)NC(C(=O)NC(Cc1ccc(O)cc1)C(=O)NC(CCC(=O)O)C(=O)NC(CS)C(=O)NC(C)C(=O)NC(CCC(N)=O)C(=O)NC(CC(N)=O)C(=O)NC(CO)C(=O)NC(Cc1ccc(O)cc1)C(=O)NC(CS)C(=O)NC(CC(N)=O)C(=O)NC(CC(=O)O)C(=O)NC(CC(C)C)C(=O)NC(CS)C(=O)NC(C(=O)NC(CCCCN)C(=O)NC(CC(N)=O)C(=O)NCC(=O)NC(C)C(=O)NC(C(=O)NC(CO)C(=O)NCC(=O)NC(Cc1ccc(O)cc1)C(=O)NC(CS)C(=O)NC(CCC(N)=O)C(=O)NC(Cc1c[nH]c2ccccc12)C(=O)NC(CC(C)C)C(=O)NCC(=O)NC(CCCCN)C(=O)NC(Cc1ccc(O)cc1)C(=O)NCC(=O)NC(CC(N)=O)C(=O)NC(C)C(=O)NC(CS)C(=O)NC(Cc1c[nH]c2ccccc12)C(=O)NC(CS)C(=O)NC(CCCCN)C(=O)NC(CC(=O)O)C(=O)NC(CC(C)C)C(=O)N1CCCC1C(=O)NC(CC(=O)O)C(=O)NC(CC(N)=O)C(=O)NC(C(=O)N1CCCC1C(=O)NC(C(=O)NC(CCCNC(=N)N)C(=O)NC(C(=O)N1CCCC1C(=O)NCC(=O)NC(CCCCN)C(=O)NC(CS)C(=O)NC(Cc1cnc[nH]1)C(=O)NC(Cc1ccccc1)C(=O)O)C(C)CC)C(C)CC)C(C)C)C(C)O)C(C)O)C(C)C,

CCC(C)C1NC(=O)C(C(C)O)NC(=O)CNC(=O)C(CCCNC(=N)N)NC(=O)C(Cc2cnc[nH]2)NC(=O)C(CC(=O)O)NC(=O)C(CO)NC(=O)C(Cc2c[nH]c3ccccc23)NC(=O)C(Cc2c[nH]c3ccccc23)NC(=O)C(CCCCN)NC(=O)C(CCCCN)NC(=O)C(Cc2ccc(O)cc2)NC(=O)C2CSSCC(NC(=O)C(CCC(=O)O)NC(=O)C(N)CC(C)C)C(=O)NC(Cc3cnc[nH]3)C(=O)NC(CC(N)=O)C(=O)NC(CCC(N)=O)C(=O)NC(CCC(N)=O)C(=O)NC(CO)C(=O)NC(CO)C(=O)NC(CCC(N)=O)C(=O)N3CCCC3C(=O)N3CCCC3C(=O)NC(C(C)O)C(=O)NC(C(C)O)C(=O)NC(CCCCN)C(=O)NC(C(C)O)C(=O)NC(CSSCC(C(=O)NCC(=O)NC3CSSCC(C(=O)NC4CSSCC(C(=O)NC(CC(N)=O)C(=O)NC(CC(N)=O)C(=O)O)NC(=O)C(CCCNC(=N)N)NC(=O)C(CC(=O)O)NC(=O)C(C(C)O)NC(=O)C(CCCNC(=N)N)NC4=O)NC(=O)C(CC(N)=O)NC(=O)C(CC(C)C)NC(=O)C(CC(N)=O)NC(=O)C(C(C)C)NC(=O)CNC(=O)C4CCCN4C(=O)C(CCCCN)NC(=O)C(C(C)C)NC(=O)C(CCCCN)NC(=O)C4CCCN4C3=O)NC(=O)CNC(=O)C(CCCNC(=N)N)NC(=O)C(CCC(=O)O)NC(=O)C(C(C)CC)NC1=O)C(=O)NC(CO)C(=O)NCC(=O)NC(CCC(=O)O)C(=O)NC(C(C)O)C(=O)NC(CC(N)=O)C(=O)N2,

CCC(C)C(NC(=O)C(Cc1ccc(O)cc1)NC(=O)CNC(=O)C(CC(=O)O)NC(=O)C(CCCCN)NC(=O)C(N)C(C)C)C(=O)NC(C(=O)NC(CC(=O)O)C(=O)NC(CC(=O)O)C(=O)NC(CCCNC(=N)N)C(=O)NC(CC(N)=O)C(=O)NC1CSSCC(C(=O)NC(CC(N)=O)C(N)=O)NC(=O)C(CCCNC(=N)N)NC(=O)CNC(=O)C2CCCN2C(=O)CNC(=O)C(CCCCN)NC(=O)C(C(C)O)NC(=O)C(CCCNC(=N)N)NC(=O)C(C(C)C)NC(=O)C(Cc2cnc[nH]2)NC(=O)C(CC(=O)O)NC(=O)C2CCCN2C(=O)C(C(C)C)NC(=O)C(CCCCN)NC(=O)C(Cc2ccc(O)cc2)NC(=O)C2CSSCC3NC(=O)C(CCC(=O)O)NC(=O)C(CCC(=O)O)NC(=O)C(CC(N)=O)NC(=O)C4CSSCC(NC(=O)C(C)NC(=O)C(CC(N)=O)NC(=O)CNC(=O)C(Cc5ccc(O)cc5)NC(=O)C5CCCN5C(=O)C(CO)NC(=O)C(C)NC(=O)C(Cc5c[nH]c6ccccc56)NC(=O)C(CCC(N)=O)NC(=O)C(CSSCC(NC(=O)C(Cc5ccccc5)NC(=O)C(Cc5ccc(O)cc5)NC(=O)C(C(C)O)NC1=O)C(=O)NCC(=O)NC(CCCNC(=N)N)C(=O)NC(CC(N)=O)C(=O)NC(C)C(=O)NC(Cc1ccc(O)cc1)C(=O)N4)NC(=O)C(Cc1ccc(O)cc1)NC(=O)CNC(=O)C(CO)NC(=O)C(CCC(=O)O)NC(=O)CNC(=O)C(CCCCN)NC(=O)C(CC(C)C)NC(=O)C(CCCCN)NC(=O)C(C(C)O)NC3=O)C(=O)NC(Cc1ccc(O)cc1)C(=O)N2)C(C)C,

CCC(C)C(NC(=O)C(Cc1ccc(O)cc1)NC(=O)CNC(=O)C(CC(=O)O)NC(=O)C(CCCNC(=N)N)NC(=O)C(NC(=O)CN)C(C)C)C(=O)NC(C)C(=O)NC(CCC(N)=O)C(=O)N1CCCC1C(=O)NC(Cc1cnc[nH]1)C(=O)NC(CC(N)=O)C(=O)NC1CSSCC(C(=O)NC(Cc2cnc[nH]2)C(=O)O)NC(=O)C(CCCCN)NC(=O)C(CCC(

=O)O)NC(=O)CNC(=O)C(CC(=O)O)NC(=O)C(C(C)C)NC(=O)C(C(C)CC)NC(=O)C(C(C)C)NC(=O)CNC(=O)C(C(C)C)NC(=O)C(CCCNC(=N)N)NC(=O)C(CC(=O)O)NC(=O)C2CCCN2C(=O)C(CC(C)C)NC(=O)C(CC(=O)O)NC(=O)C(CCCCN)NC(=O)C2CSSCC3NC(=O)C(CC(C)C)NC(=O)C(C(C)O)NC(=O)C(CC(=O)O)NC(=O)C4CSSCC(NC(=O)C(C)NC(=O)C(C(C)O)NC(=O)CNC(=O)C(CCCNC(=N)N)NC(=O)CNC(=O)C(CC(C)C)NC(=O)C(C(C)CC)NC(=O)C(Cc5ccccc5)NC(=O)C(CSSCC(NC(=O)C(Cc5cnc[nH]5)NC(=O)C(Cc5ccc(O)cc5)NC(=O)C(C(C)C)NC1=O)C(=O)NC(Cc1ccccc1)C(=O)N1CCCC1C(=O)NCC(=O)NC(CO)C(=O)NCC(=O)NCC(=O)N4)NC(=O)C(CO)NC(=O)C(CO)NC(=O)CNC(=O)C(CCC(N)=O)NC(=O)C(C(C)O)NC(=O)C(C)NC(=O)CNC(=O)C(CC(N)=O)NC(=O)C(CCC(=O)O)NC(=O)C(CCCCN)NC3=O)C(=O)NC(Cc1c[nH]c3ccccc13)C(=O)N2,

C=C(C=CCC(O)C(O)C(O)CC=CC=CC(O)CC1OC(CC(O)C(O)CC2OC(C(O)C(O)CCC(O)C=CC(C)C(O)CC3(O)OC(CC(O)CCCCCCCC45CC(C)CC(C)(O4)C(CC(C)CCCCCC(O)C(O)C(O)C(O)C(O)C4OC(CC(O)C(O)C(C)=CC(O)CC(C)C(O)C(=O)NC=CC(=O)NCCCO)C(O)C(O)C4O)O5)C(O)C(O)C3O)CC(O)C2O)C(O)C(O)C1O)CCC(O)C(O)C(O)C(C)CC1OC(C=CC(O)C(O)CC2CC3CC(O2)C(CCC2OC(CN)CC2O)O3)C(O)C(O)C1O,

CC(CCC1(O)OC2CC3C4CC=C5CC(OC6OC(CO)C(O)C(OC7OC(CO)C(OC8OC(CO)C(O)C(OC9OC(CO)C(O)C(OC%10OC(CO)C(O)C(O)C%10O)C9O)C8O)C(O)C7O)C6O)CCC5(C)C4CCC3(C)C2C1C)COC1OC(CO)C(O)C(O)C1O,

CCC(C)C1NC(=O)C(CCCCN)NC(=O)C2CSSCC(NC(=O)C(NC(=O)C(CO)NC(=O)C(N)CCCCN)CSSCC(C(=O)N3CCCC3C(=O)NC(CCCCN)C(=O)NCC(=O)NC(Cc3ccccc3)C(=O)N3CCCC3C(=O)NC(CCCCN)C(=O)O)NC(=O)C(CO)NC(=O)C(CC(C)C)NC(=O)CNC(=O)C(CO)NC(=O)C(CO)NC1=O)C(=O)NC(CCCNC(=N)N)C(=O)NC(CO)C(=O)NC(C(C)O)C(=O)NC(CC(C)C)C(=O)NCC(=O)NC(CCCNC(=N)N)C(=O)NC(CC(N)=O)C(=O)NC1CSSCC(NC(=O)C(C(C)C)NC(=O)CNC(=O)C(C)NC(=O)C3CSSCC(NC(=O)C(CC(C)C)NC(=O)C(CC(N)=O)NC(=O)C(Cc4ccc(O)cc4)NC1=O)C(=O)NC(CCCNC(=N)N)C(=O)NC(C)C(=O)NC(CCCNC(=N)N)C(=O)NCC(=O)NC(C)C(=O)NC(CCC(N)=O)C(=O)NC(CCCCN)C(=O)NC(CC(C)C)C(=O)N3)C(=O)NC(CCCNC(=N)N)C(=O)N2,

CC1(C)CCC2(C(=O)OC3OC(CO)C(O)C(OC4OC(CO)C(O)C(OC5OC(CO)C(O)C(OC6OC(CO)C(O)C(OC7OC(CO)C(O)C(O)C7O)C6O)C5O)C4O)C3O)C(O)CC3(C)C(=CCC4C5(C)CCC(OC6OC(CO)C(O)C(OC7OC(CO)C(O)C(OC8OC(CO)C(O)C(O)C8O)C7O)C6O)C(C)(C=O)C5CCC43C)C2C1,

COC(=O)C1NC(=O)C2NC(=O)C(NC(=O)C3NC(=O)C4NC(=O)C(NC(=O)C(N)c5ccc(O)c(c5)Oc5cc4cc(O)c5C)C(O)c4ccc(cc4)Oc4cc3cc(c4OC3OC(COC4OC(C)C(O)C(O)C4O)C(O)C(O)C3O)Oc3ccc(cc3)C2OC2CC(N)C(O)C(C)O2)c2ccc(O)c(c2)-c2c(OC3OC(CO)C(O)C(O)C3O)cc(O)cc21,

COC(=O)C1NC(=O)C2NC(=O)C(NC(=O)C3NC(=O)C4NC(=O)C(NC(=O)C(N)c5ccc(O)c(c5)Oc5cc4cc(O)c5C)C(O)c4ccc(cc4)Oc4cc3cc(c4OC3OC(COC4OC(C)C(O)C(O)C4O)C(O)C(O)C3OC3OC(CO)C(O)C(O)C3OC3OCC(O)C(O)C3O)Oc3ccc(cc3)C2OC2CC(N)C(O)C(C)O2)c2ccc(O)c(c2)-c2c(OC3OC(CO)C(O)C(O)C3O)cc(O)cc21,

COC(=O)C1NC(=O)C2NC(=O)C(NC(=O)C3NC(=O)C4NC(=O)C(NC(=O)C([NH3+])c5ccc(O)c(c5)Oc5cc4cc(O)c5C)C(O)c4ccc(cc4)Oc4cc3cc(c4OC3OC(COC4OC(C)C(O)C(O)C4O)C(O)C(O)C3OC3OC(CO)C(O)C(O)C3OC3OCC(O)C(O)C3O)Oc3ccc(cc3)C2OC2CC([NH3+])C(O)C(C)O2)c2ccc(O)c(c2)-c2c(OC3OC(CO)C(O)C(O)C3O)cc(O)cc21.O=S(=O)([O-])[O-],

COC(=O)C1NC(=O)C2NC(=O)C(NC(=O)C3NC(=O)C4NC(=O)C(NC(=O)C(N)c5ccc(O)c(c5)Oc5cc4cc(O)c5C)C(O)c4ccc(cc4)Oc4cc3cc(c4O)Oc3ccc(cc3)C2OC2CC(N)C(O)C(C)O2)c2ccc(O)c(c2)-c2c(O)cc(O)cc21,

CCCCCC=CCCC(=O)NC1C(Oc2c3cc4cc2Oc2ccc(cc2Cl)C(OC2OC(CO)C(O)C(O)C2NC(C)=O)C2NC(=O)C(NC(=O)C4NC(=O)C4NC(=O)C(NC(=O)C(N)c5ccc(O)c(c5)Oc5cc4cc(O)c5C)C(O)c4ccc(c(Cl)c4)O3)c3ccc(O)c(c3)-c3c(OC4OC(CO)C(O)C(O)C4O)cc(O)cc3C(C(=O)OC)NC2=O)OC(CO)C(O)C1O,

CCCCCC=CCCC(=O)NC1C(Oc2c3cc4cc2Oc2ccc(cc2Cl)C(OC2OC(CO)C(O)C(O)C2NC(C)=O)C2NC(=O)C(NC(=O)C4NC(=O)C4NC(=O)C(Cc5ccc(c(Cl)c5)O3)NC(=O)C(N)c3ccc(O)c(c3)Oc3cc(O)cc4c3)c3ccc(O)c(c3)-c3c(OC4OC(CO)C(O)C(O)C4O)cc(O)cc3C(C(=O)O)NC2=O)OC(CO)C(O)C1O,

CC(=O)NC1C(OC2c3ccc(c(Cl)c3)Oc3cc4cc(c3OC3OC(CO)C(O)C(O)C3NC(=O)CCCCCCC(C)C)Oc3ccc(cc3Cl)CC3NC(=O)C(N)c5ccc(O)c(c5)Oc5cc(O)cc(c5)C(NC3=O)C(=O)NC4C(=O)NC3C(=O)NC2C(=O)NC(C(=O)O)c2cc(O)cc(OC4OC(CO)C(O)C(O)C4O)c2-c2cc3ccc2O)OC(CO)C(O)C1O,

CCCCCCCCCC(=O)NC1C(Oc2c3cc4cc2Oc2ccc(cc2Cl)C(OC2OC(CO)C(O)C(O)C2NC(C)=O)C2NC(=O)C(NC(=O)C4NC(=O)C4NC(=O)C(Cc5ccc(c(Cl)c5)O3)NC(=O)C(N)c3ccc(O)c(c3)Oc3cc(O)cc4c3)c3ccc(O)c(c3)-c3c(OC4OC(CO)C(O)C(O)C4O)cc(O)cc3C(C(=O)O)NC2=O)OC(CO)C(O)C1O,

CCC(C)CCCCCCC(=O)NC1C(Oc2c3cc4cc2Oc2ccc(cc2Cl)C(OC2OC(CO)C(O)C(O)C2NC(C)=O)C2NC(=O)C(NC(=O)C4NC(=O)C4NC(=O)C(Cc5ccc(c(Cl)c5)O3)NC(=O)C(N)c3ccc(O)c(c3)Oc3cc(O)cc4c3)c3ccc(O)c(c3)-c3c(OC4OC(CO)C(O)C(O)C4O)cc(O)cc3C(C(=O)O)NC2=O)OC(CO)C(O)C1O,

CC(=O)NC1C(OC2c3ccc(c(Cl)c3)Oc3cc4cc(c3OC3OC(CO)C(O)C(O)C3NC(=O)CCCCCCCC(C)C)Oc3ccc(cc3Cl)CC3NC(=O)C(N)c5ccc(O)c(c5)Oc5cc(O)cc(c5)C(NC3=O)C(=O)NC4C(=O)NC3C(=O)NC2C(=O)NC(C(=O)O)c2cc(O)cc(OC4OC(CO)C(O)C(O)C4O)c2-c2cc3ccc2O)OC(CO)C(O)C1O,

CCC(C)C(NC(=O)C(C)NC(=O)C(CCCNC(=N)N)NC(=O)C(CO)NC(=O)C(NC(=O)C(NC(=O)C(NC(=O)C(Cc1cnc[nH]1)NC(=O)C(CO)NC(=O)C(Cc1ccc(O)cc1)NC(=O)C(CS)NC(=O)C(NC(=O)C(N)C(C)O)C(C)CC)C(C)O)C(C)O)C(C)O)C(=O)NC(CC(C)C)C(=O)NC(CCCCN)C(=O)NC(CC(=O)O)C(=O)NC(CS)C(=O)NCC(=O)NC(CCC(=O)O)C(=O)NC(CC(N)=O)C(=O)NC(CO)C(=O)NC(CS)C(=O)NC(Cc1ccc(O)cc1)C(=O)NC(CCCNC(=N)N)C(=O)NC(CCCCN)C(=O)NC(CO)C(=O)NC(CCCNC(=N)N)C(=O)NC(CCCNC(=N)N)C(=O)NC(Cc1cnc[nH]1)C(=O)N1CCCC1C(=O)N1CCCC1C(=O)NC(CCCCN)C(=O)NC(CCSC)C(=O)NC(C(=O)NC(CC(C)C)C(=O)NCC(=O)NC(CCCNC(=N)N)C(=O)NCC(=O)NC(CS)C(=O)NCC(=O)NC(CS)C(=O)N1CCCC1C(=O)N1CCCC1C(=O)NCC(=O)NC(CC(=O)O)C(=O)NC(CC(=O)O)C(=O)NC(Cc1ccc(O)cc1)C(=O)NC(CC(C)C)C(=O)NC(CCC(=O)O)C(=O)NC(C(=O)NC(CCCCN)C(=O)NC(CS)C(=O)NC(CS)C(=O)NC(C(=O)NC(CO)C(=O)N1CCCC1C(=O)NC(CC(=O)O)C(=O)NC(CCCCN)C(=O)NC(CS)C(=O)NC(CC(N)=O)C(=O)NC(Cc1ccc(O)cc1)C(=O)O)C(C)O)C(C)C)C(C)C,

C=C(CO)CC1CC(O)C2(C)OC3CC4OC5CC6(C)OC7(C)CCC8OC9CC%10(C)OC%11C(C)=CC(=O)OC%11CC%10OC9CC(C)C8OC7CC6OC5(C)CC=CC4OC3CC2O1,

C=C(C=O)CC1CC(O)C2(C)OC3CC4OC5CC6(C)OC7(C)CCC8OC9CC%10(C)OC%11C(C)=CC(=O)OC%11CC%10OC9CC(C)C8OC7CC6OC5(C)CC=CC4OC3CC2O1,

CCC(C)C(NC(=O)C(CS)NC(=O)CNC(=O)C(CCCNC(=N)N)NC(=O)C(CCCCN)NC(=O)C(NC(=O)C1CCCN1C(=O)C(NC(=O)C(NC(=O)C(CO)NC(=O)C(NC(=O)C(CO)NC(=O)C(NC(=O)C(CCSC)NC(=O)C(Cc1ccccc1)NC(=O)C(CCSC)NC(=O)C(CCCCN)NC(=O)C(Cc1ccc(O)cc1)NC(=O)C(CS)NC(=O)C(CC(C)C)NC(=O)C(CC(N)=O)NC(=O)C(CCCCN)NC(=O)CNC(=O)C(CCC(=O)O)NC(=O)C1CCCN1C(=O)C(CS)NC(=O)C(NC(=O)C(CCCCN)NC(=O)C(Cc1c[nH]c2ccccc12)NC(=O)C(NC(=O)C1CCCN1C(=O)C1CCCN1C(=O)C(NC(=O)C(CC(C)C)NC(=O)C(CCCCN)NC(=O)C(Cc1cnc[nH]1)NC(=O)C(CS)NC(=O)C(CCCCN)NC(=O)C(N)CC(C)C)C(C)C)C(C)C)C(C)O)C(C)C)C(C)O)C(C)O)C(C)C)C(C)C)C(=O)NC(CC(=O)O)C(=O)NC(C(=O)NC(CS)C(=O)N1CCCC1C(=O)NC(CCCCN)C(=O)NC(CC(=O)O)C(=O)NC(CO)C(=O)NC(C)C(=O)NC(CC(C)C)C(=O)NC(C(=O)NC(CCCCN)C(=O)NC(Cc1ccc(O)cc1)C(=O)NC(C(=O)NC(CS)C(=O)NC(CS)C(=O)NC(CO)C(=O)NC(C(=O)NC(CC(=O)O)C(=O)NC(CCCCN)C(=O)NC(CS)C(=O)NC(CC(N)=O)C(=O)O)C(C)O)C(C)C)C(C)C)C(C)C,

CCC(C)C(NC(=O)C(CS)NC(=O)CNC(=O)C(CCCNC(=N)N)NC(=O)C(CCCCN)NC(=O)C(NC(=O)C1CCCN1C(=O)C(NC(=O)C(NC(=O)C(CO)NC(=O)C(NC(=O)C(CO)NC(=O)C(NC(=O)C(CCSC)NC(=O)C(Cc1ccc(O)cc1)NC(=O)C(CCSC)NC(=O)C(CC(N)=O)NC(=O)C(Cc1ccc(O)cc1)NC(=O)C(CS)NC(=O)C(CC(C)C)NC(=O)C(CC(N)=O)NC(=O)C(CCCCN)NC(=O)CNC(=O)C(CCCCN)NC(=O)C1CCCN1C(=O)C

(CS)NC(=O)C(NC(=O)C(CCCCN)NC(=O)C(Cc1c[nH]c2ccccc12)NC(=O)C(Cc1ccccc1)NC(=O)C1CCCN1C(=O)C1CCCN1C(=O)C(NC(=O)C(CC(C)C)NC(=O)C(CCC(N)=O)NC(=O)C(CC(N)=O)NC(=O)C(CS)NC(=O)C(CCCCN)NC(=O)C(N)CC(C)C)C(C)CC)C(C)O)C(C)C)C(C)O)C(C)O)C(C)C)C(C)C)C(=O)NC(CC(=O)O)C(=O)NC(C(=O)NC(CS)C(=O)N1CCCC1C(=O)NC(CCCCN)C(=O)NC(CC(N)=O)C(=O)NC(CO)C(=O)NC(C)C(=O)NC(CC(C)C)C(=O)NC(C(=O)NC(CCCCN)C(=O)NC(Cc1ccc(O)cc1)C(=O)NC(C(=O)NC(CS)C(=O)NC(CS)C(=O)NC(CC(N)=O)C(=O)NC(C(=O)NC(CC(=O)O)C(=O)NC(CCCNC(=N)N)C(=O)NC(CS)C(=O)NC(CC(N)=O)C(=O)O)C(C)O)C(C)C)C(C)C)C(C)C,

COC(=O)C1c2cc3c(c(O)c2C(OC2CC(O)C(OC4CC(O)C(OC5CC(C)(N=O)C(OC6CC(OC)C(OC(=O)C(C)=CC(=O)O)C(C)O6)C(C)O5)C(C)O4)C(C)O2)CC1(C)O)C(=O)c1c(O)cc2c(c1C3=O)CC1CC(N(C)C)C(OC3CC(O)C(OC4CC(C)(N=O)C(OC5CC(O)C(O)C(C)O5)C(C)O4)C(C)O3)C2(C)O1,

CCC(C)C(NC(=O)C(CC(N)=O)NC(=O)C(CCCNC(=N)N)NC(=O)CNC(=O)C(NC(=O)C(NC(=O)C(CC(N)=O)NC(=O)C1CCCN1C(=O)C(CS)NC(=O)C(CS)NC(=O)C(CO)NC(=O)C(N)CCCCN)C(C)O)C(C)O)C(=O)NC(Cc1ccc(O)cc1)C(=O)NC(CC(N)=O)C(=O)NCC(=O)NC(CS)C(=O)NC(CCCNC(=N)N)C(=O)NCC(=O)NCC(=O)NCC(=O)NCC(=O)NCC(=O)NC(CCCNC(=N)N)C(=O)NCC(=O)NCC(=O)NC(CS)C(=O)NC(C)C(=O)NCC(=O)NCC(=O)NC(CO)C(=O)NCC(=O)NC(CS)C(=O)NC(CCCCN)C(=O)NC(C(=O)NC(C(=O)NC(CO)C(=O)NCC(=O)NC(CO)C(=O)NC(C(=O)NC(CS)C(=O)N1CCCC1C(=O)NC(CO)C(=O)NC(Cc1ccc(O)cc1)C(=O)N1CCCC1C(=O)NC(CC(=O)O)C(=O)NC(CCCCN)C(=O)O)C(C)O)C(C)CC)C(C)CC,

C=CCC(=C)C=CC(C)(O)C1OC2CC3OC4CC5OC6CC7OC8CC9OC%10CC%11OC(C)(CCOS(=O)(=O)O)C(OS(=O)(=O)O)CC%11OC%10CC9OC8CCC7(C)OC6(C)CCC(C)C5OC4C(O)C3(C)OC2CC1=C,

O=S(=O)(O)c1cc(Nc2nc(Cl)nc(Nc3ccc(C=Cc4c(Nc5nc(Cl)nc(Nc6cc(S(=O)(=O)O)cc7cc(S(=O)(=O)O)c(N=Nc8ccc9ccccc9c8S(=O)(=O)O)c(O)c67)n5)cccc4S(=O)(=O)O)c(S(=O)(=O)O)c3)n2)c2c(O)c(N=Nc3ccc4ccccc4c3S(=O)(=O)O)c(S(=O)(=O)O)cc2c1,

CC1(C)Nc2cccc3c(N=Nc4ccc(N=Nc5ccccc5)c5ccccc45)ccc(c23)N1
